# The *Drosophila melanogaster* Genetic Reference Panel, Version 3 (DGRP3)

**DOI:** 10.64898/2026.05.29.728783

**Authors:** Richard F. Lyman, Vijay Shankar, Rachel A. Lyman, Lakshmi Sunkara, Nestor O. Nazario Yepiz, Maria E. Adonay, Jordyn Brooks, Roberta L. Lyman, Yuefan Huang, Wen Huang, Robert R. H. Anholt, Trudy F. C. Mackay

**Author notes:** Corresponding author Contact Information: Dr. Trudy F. C Mackay Institute for Human Genetics Clemson University 114 Gregor Mendel Circle Greenwood, SC 29646. Equal contributions (Richard F. Lyman, Vijay Shankar, Rachel A. Lyman).

## Abstract

*Drosophila melanogaster* is a leading animal model for understanding basic genetic principles for biomedical research and for quantitative, population and evolutionary genetics and genomics. The *D. melanogaster* Genetic Reference Panel (DGRP) of ∼200 inbred strains with full genome sequences is a publicly available resource for genome wide association (GWA) studies of quantitative traits, systems genetics, and population genomic analyses. However, the small size of the DGRP limits the power to map rare variants or variants with small effect sizes. Here, we describe the DGRP Version 3 (DGRP3), consisting of full genome sequences from 1,233 inbred lines, of which 1,037 are currently extant. The DGRP3 harbors extensive genetic diversity, with 8,760,339 molecular polymorphisms including single nucleotide polymorphisms, insertions, deletions, inversions, tandem duplications and complex rearrangements. We identified 80,894 damaging variants and 4,550 genes with at least one loss-of-function haplotype. We inferred 16 mitochondrial haplotype groups, determined the microbial composition, and quantified the numbers of sensory bristles for the DGRP3 lines. We performed GWA analyses for inbreeding depression, host control of microbiome composition, and numbers of sensory bristles, identifying novel candidate variants and genes for each. The accompanying online tool (flydgrp.org) implements state-of-the-art association mapping methods and provides results for users’ DGRP3 data.

## Introduction

*Drosophila melanogaster* has been used for over a century to understand the genetic principles of mutation, linkage and recombination and their underlying mechanisms; and to use induced mutations to map and clone genes affecting development, growth, differentiation, behaviors and other complex traits. These include canonical cell signaling pathways (Notch, Wingless/Wnt, Hedgehog, Toll) named after genes first described in *D. melanogaster*. The genome sequences of *D. melanogaster* and humans were both published at the beginning of the 21^st^ century^1,2^, leading to the realization that 75% of human disease genes have *Drosophila* orthologs. Many publicly available resources are available for *D. melanogaster* to manipulate genes and proteins, including mutations in nearly all genes in the genome^3,4^, reagents for reducing or increasing gene expression in specific tissues and cell types with temporal control^5–8^, and databases of gene expression for tissues, developmental stages and single cells^9,10^. Combined with low maintenance costs, a short generation time and few regulatory restrictions, these features make *D. melanogaster* a leading animal model for biomedical research.

*D. melanogaster* is also a leading animal model for quantitative, population and evolutionary genetics and genomics. This species has been used to test quantitative genetics theory^11–16^ and estimate the spontaneous mutation rate for fitness^17–19^ and other quantitative traits^20–25^. The *D. melanogaster* model has also been used to pioneer molecular population genetic analyses to estimate local nucleotide diversity and linkage disequilibrium (LD) near the first genes that were cloned in this species and determine the relationships between nucleotide diversity, LD and local recombination rate^26–31^. Some of the first experiments to map quantitative trait loci (QTLs) in linkage^32–38^ and association^39–43^ mapping paradigms were performed in *D. melanogaster*. Molecular population genetic analyses showed that LD decays rapidly in regions of normal recombination in this species, necessitating complete genome sequences to perform genome-wide association (GWA) mapping analyses.

The *D. melanogaster* Genetic Reference Panel (DGRP) was developed as a publicly available community resource for GWA and population genomic analyses. The first version of the DGRP (DGRP1) consisted of 168 lines derived by 20 generations of full sib mating from progeny of single inseminated females collected between 1998 and 2002 from the Raleigh, NC USA Farmer’s Market^44^. These lines were sequenced by a combination of 454 and Illumina Genome Analyzer IIx (GAII) chemistry, and single nucleotide polymorphisms (SNPs), polymorphic microsatellites and transposable element insertion sites were reported. The DGRP2 added 37 more inbred lines derived from the same population (for a total of 205 lines), sequenced to an average depth of 27X by Illumina GAII or HiSeq 2000 platforms^45^. SNPs, insertion and deletion polymorphisms, tandem duplications, complex variants, karyotypes of polymorphic inversions and *Wolbachia pipientis* infection status were determined for all 205 lines. The DGRP2 harbors over 6 million molecular variants, of which nearly 2 million are common, with minor allele frequencies (MAF) ≥ 0.05.

Because the DGRP lines are inbred, large numbers of individuals of the same genotype can be measured, reducing the pooled environmental variance and increasing the precision of estimates line means, which increases mapping power. Replicated genotypes also facilitate the detection of pleiotropic effects on multiple traits, including the organismal and molecular traits required for systems genetics analyses^46–49^, and for quantifying the magnitude of genotype by environment interaction^50,51^. Each inbred line is nearly homozygous, but the additive genetic variance between the inbred lines is twice that of the outbred population from which they were derived^52^. The contribution of any additive × additive epistatic variance is increased four-fold and three-way additive epistatic interaction variance by a factor of eight^53^, so the DGRP can be exploited to map epistatic interactions^54,55^. A freely available online tool for mapping molecular variants associated with any quantitative trait measured on the DGRP lines made using this resource widely accessible to the community, including for classroom teaching. There are over 200 publications to date using the DGRP to map variants and genes associated with morphological, behavioral, physiological, life history and molecular quantitative traits, including mapping modifiers of *Drosophila* models of human disease^56–59^. The rapid decline in LD with physical distance in the DGRP gives high mapping resolution, often to a few base pairs, and mutations and RNA interference constructs can be used to functionally validate associations at the levels of genes.

However, the small size of the DGRP2 limits the power to map rare variants or variants with small effect sizes associated with quantitative traits^57^. Further, the DGRP lines must be propagated by living animals; therefore, the published sequence data no longer match the extant genomes due to the accumulation of new spontaneous mutations and changes in allele frequency of any segregating sites from random genetic drift over the 15 years of stock maintenance since the original sequences were obtained. Here, we describe the DGRP Version 3 (DGRP3), consisting of sequences from 1,233 inbred lines derived from the same Raleigh, NC population. The DGRP3 harbors extensive genetic diversity, with 8,760,339 molecular polymorphisms and structural variants, including SNPs, insertions, deletions, inversions, tandem duplications and complex rearrangements. Inbreeding depression resulted in the loss of 196 lines. The 1,037 viable DGRP3 lines comprise one of the largest publicly available eukaryotic association mapping reference panels. The DGRP3 has greater statistical power to detect associations than the DGRP2^45^ and enables evaluation of effects of variants with minor allele frequencies in the range of 0.01-0.05, which tend to have larger effects^44,45^. The accompanying online tool implements state-of-the-art association mapping methods accounting for relatedness and provides results to users based on their own DGRP3 data and links to external resources.

## Results

### Molecular Polymorphisms in the DGRP3

The DGRP3 consists of 1,058 lines derived by 20 generations of full-sib inbreeding of progeny from inseminated females collected at the Raleigh, NC USA Farmer’s Market between 2015-2017, and 175 DGRP2 lines that were re-inbred by 5 generations of full-sib inbreeding (Table S1A). The original DGRP2 lines were collected between 1998-2002 from the same location and were also established by 20 generations of full-sib inbreeding. All 1,233 lines were sequenced using Illumina NovaSeq 6000 technology to an average depth of 46.5X (Table S1B). The DGRP3 lines are available from the Bloomington, Indiana USA Drosophila Stock Center and the sequences are available from the National Center for Biotechnology Information Short Read Archive (SRA). SRA accession numbers are given in Table S1B.

### Nuclear Genomic Variants

We identified a total of 8,692,287 biallelic short nucleotide variants (SNVs, ≤ 50 bp in length), of which 7,105,829 (81.7%) were single nucleotide variants (SNPs), 978,710 (11.3%) were deletions and 607,748 (7.0%) were insertions (collectively, indels). A total of 18,273 genes and 19,235 transcripts overlapped with SNV coordinates. The most common variant site class was intronic (44.2%) followed by intergenic (19.6%) (Table S2A). Among coding variants, 52.2% were synonymous and 42.3% were missense (Table S2B). SNVs and indels were distributed throughout the genome proportionate to chromosome size (Table S2C, Figure 1A). There were fewer SNVs near telomeres, centromeres and pericentromeric heterochromatic regions (Figure 1A), consistent with previous observations that polymorphism scales positively with recombination rate in this species^31,44,60^. This pattern is also evident from estimates of nucleotide diversity^61^ (x), calculated separately for SNPs and indels in 10 Mb windows across each chromosome arm (Figure 2A, Table S3). The average nucleotide diversity for SNPs is 0.00311 for the *X* chromosome and 0.00436 for the major autosome arms. The average nucleotide diversity for indels is 0.00058 over the whole genome (excluding the small *4*^th^ chromosome), with little difference between the *X* chromosome and autosomes (Table S3). The estimate of x for SNPs over the whole genome (x = 0.411) is somewhat lower than estimated in the DGRP1^44^, likely attributable to more rare variants in the larger DGRP3 population. The patterning of x for indels across the major chromosome arms is strongly positively correlated with that of SNPs (average r = 0.747, Table S3), as previously observed in the DGRP2^45^.

**Figure 1.**
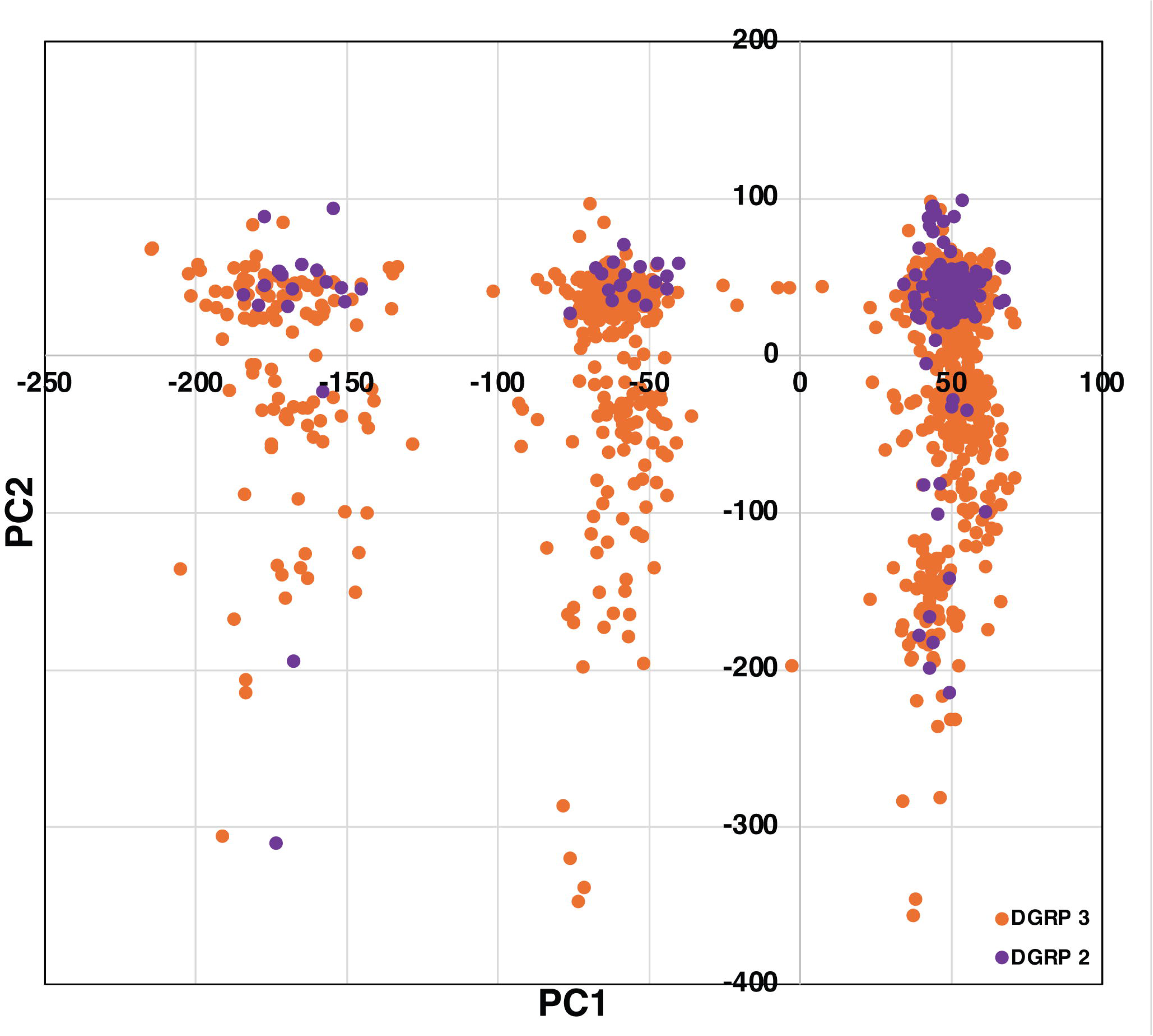
Genomic distributions of polymorphic variants. (A) Nuclear genome. The Circos plot depicts the locations of genes, SNVs (SNPs and indels) and common large polymorphic inversions on the *D. melanogaster* nuclear genome. The major chromosome arms and genomic positions are shown in the outer ring. The same data for the small *4*^th^ chromosome are depicted in the inset. (B) Mitochondrial genome. The Circos plot depicts the locations of genes and SNVs (SNPs and indels) on the mitochondrial genome.

**Figure 2.**
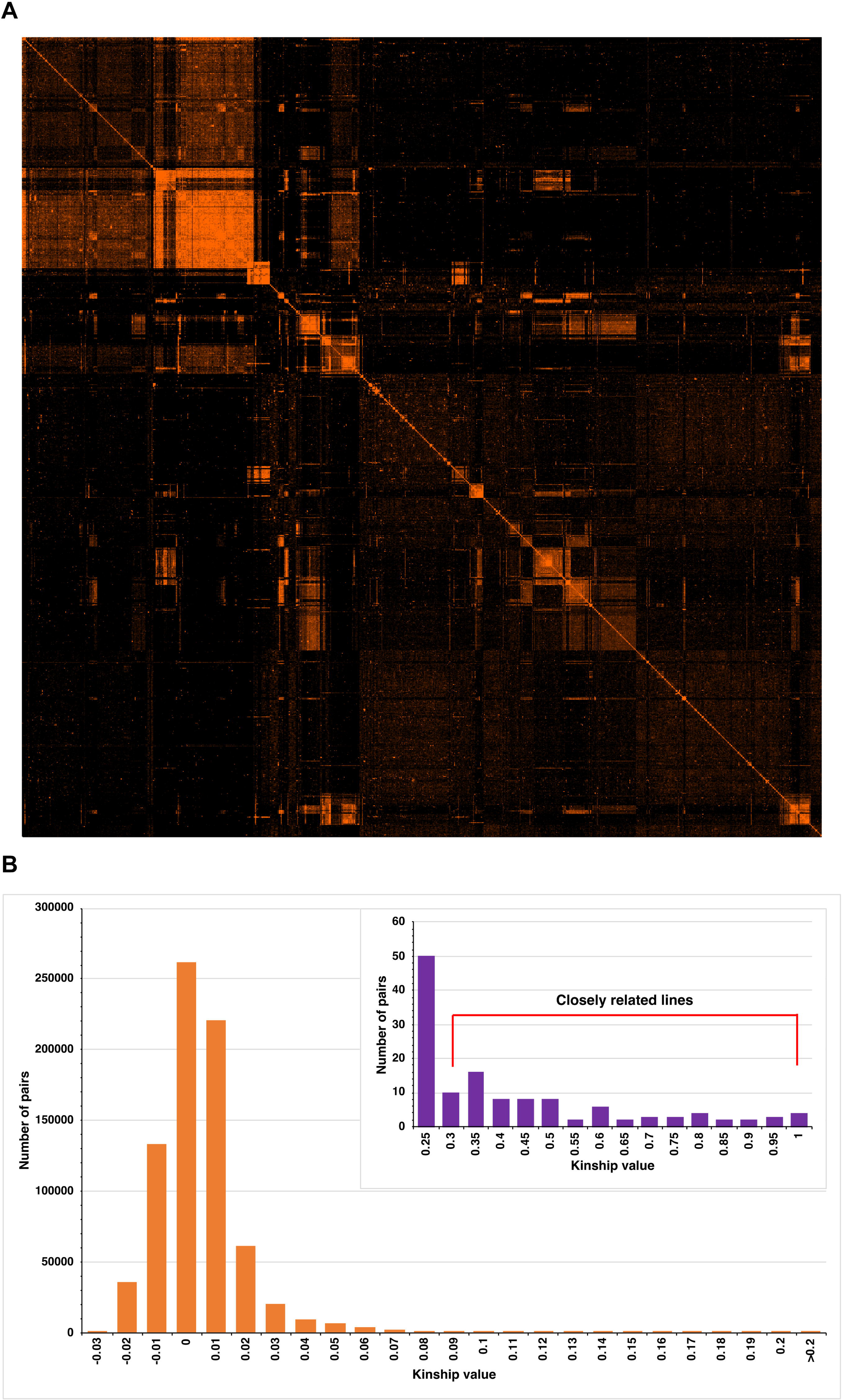
**Distribution of nucleotide diversity (**n^61^**) and Tajima’s** D^68^ **estimates.** (A) Left coumn. Average values of x in adjacent 10 kb windows are shown for SNPs (purple dots) and indels (orange dots) along each major chromosome arm. (B) Average values of D in adjacent 10 kb windows are shown for SNPs (purple dots) and indels (orange dots) along each major chromosome arm.

We also identified multiple polymorphic structural variants (> 50 bp). These include 28,851 insertions, 30,184 deletions, 3,521 tandem duplications, 4,443 interchromosomal translocations and 1,053 inversions (Table S4). Many of these structural variants co-occur in heterochromatic genomic regions: cytobands 1A-2F and 19A-20F on the *X* chromosome; 21A-22F and 39A-40F on chromosome *2L*; 41A-42F and 59A-60F on chromosome *2R*; 61A-62F and 79A-80F on chromosome *3L*; and 81A-82F and 99A-100F on chromosome *3R* (Table S4). These regions are characterized by repetitive DNA regions, which appear to be susceptible to the formation of structural variation. Indeed, 176 of the 286 chromosome *3L* inversions (61%) and 91 of the 209 chromosome *3R* inversions (43.5%) have one or both breakpoints in cytobands 79A-80F and 81A-82F, respectively. However, repeated regions do not account for breakpoints of most structural variants (Table S4).

Large polymorphic chromosome inversions were among the first genetic polymorphisms studied in *Drosophila*, since they could be identified using polytene salivary chromosome banding patterns^62,63^. Nearly 400 polymorphic inversions have been identified in natural populations of *D. melanogaster* worldwide based on their cytogenetic breakpoints^64^. Most of these inversions are rare and endemic to a single population. Four polymorphic inversions are found in nearly all populations (*In(2L)t*, *In(2R)NS*, *In(3L)P*, *In(3R)P*) and six additional inversions (*In(2R)Cy*, *In(3L)M*, *In(3R)C*, *In(3R)K*, *In(3R)Mo*, *In(3R)M*) are widespread^64^. We identified 31 large (> 1 Mb) common (inversion frequency > 0.03) polymorphic inversions in the DGRP3, of which three are historically found in most populations (*In(2L)t*, *In(2R)NS*, *In(3L)P*) and two are widespread (*In(3R)K*, *In(3R)Mo*) (Table S5). Compared to the DGRP2, the frequencies of *In(2L)t* and *In(3R)Mo* decreased. The remaining 26 common large inversion polymorphisms are new and did not map to the cytological locations of any previously described inversion^45,64^. These observations are consistent with previous observations that most inversions are young, rare and population-specific^64,65^. The large inversions are not randomly distributed in the genome and tend to overlap, particularly for chromosomes *3L* and *3R* (Figure 1A).

Most molecular variants are rare, with minor allele frequencies (MAF) < 0.01 (Figure S1). Note that we calculated MAFs assuming that lines with a segregating variant are heterozygous for the variant, which is not necessarily true, since we sequenced pools of flies and therefore do not know the genotypes of individuals for segregating variants. Rare alleles are more likely to have deleterious effects on fitness than common alleles^66,67^. We inferred that 35,509 of the single nucleotide variants were deleterious due to frameshift, splice acceptor, splice donor, start lost, stop gained, stop lost, and ablated transcript effects (Table S2, Table S6). The structural variants often disrupted genes (*i.e.*, occurred in the gene body), with potential deleterious effects. Of the 28,851 insertion variants, 19,215 (66.6%) disrupted coding genes. Similarly, 19,283 of 30,184 deletions (63.9%), 2,696 of 3,521 duplications (76.8%), 728 of 1,053 inversions (69.4%) and 3,463 of 4,443 translocations (77.9%) disrupted coding genes (*i.e*., if the start and/or end occurred in the gene body). We also identified loss-of-function (LoF) effects of protein coding genes and found that each DGRP line harbored on average 103.9 LoF genes, with a wide range between 18 and 229 per line (Figure S2A, Table S6). Among the 13,840 protein coding genes that were assessed, 4,550 (32.9%) contained at least one LoF haplotype in the population. The frequency distribution was highly skewed, with the majority of LoF genes present in one (1,576), two (660), or three (392) lines (Figure S2B, Table S6).

Prediction of deleterious effects of variants is constrained to variants that impact protein coding regions. However, most molecular variation observed in the DGRP occurs in non-coding regions. Another signature of deleterious effects that applies genome-wide are variants that only occur as segregating variants within inbred lines. This can be an indication that natural selection opposes the fixation of these presumably deleterious recessive variants during inbreeding, so there are many fewer homozygotes than expected given the allele frequency. For example, 21 of the 26 new common inversions had appreciable frequencies of segregating standard and inverted karyotypes but no or few inversion homozygote(s) (Table S5). In addition, there were no homozygous inversion karyotypes for *In(2R)NS* and *In(3R)K*.

To assess global patterns of selection and demography, we calculated Tajima’s D statistics^68^ across each chromosome arm in 10 kb windows. As expected from previous analyses on smaller samples, Tajima’s D for SNPs (*D_SNP_*) is negative averaged over all major chromosome arms (*D_SNP_* = −1.102) (Table S7A-F), consistent with an excess of low frequency molecular polymorphisms caused by population bottlenecks when colonizing North America, followed by population expansion^69,70^. However, *D_SNP_* is lower on the *X* chromosome (*D_SNP_* = −1.210) than major autosomes (*D_SNP_* = −1.104), and lower still on the *4*^th^ chromosome (*D_SNP_* = −2.109) (Figure 2B, Table S7A-F). *D_SNP_* is also reduced in regions of lower recombination near the telomeres and centromeres of the major chromosome arms, less so for the *X* chromosome than autosomes (Figure 2B, Table S7A-F), as previously observed^44^. The decrease in *D_SNP_* in regions of reduced recombination is likely attributable to background selection^66^, where linked neutral loci are reduced in frequency by selection against rare deleterious alleles in these regions, while heterogeneity in estimates of *D_SNP_* in normal regions of recombination may be due to either selection against deleterious mutations or local selective sweeps for adaptive mutations^71^. D for indels (*D_indel_*) is on average lower than *D_SNP_* (Figure 2B, Table S7A-F). *D_indel_* = −1.627 averaged over all major chromosome arms, indicating that indels are more deleterious than SNPs, as observed previously^45^. In addition, we computed *D_SNP_* and *D_indel_* by gene as the average of sequences 1 kb from the transcription start and end site and in 1 kb intervening windows (Table S7G). Of the 306 genes with *D_SNP_* < −2.000, the most negative values were for noncoding RNAs: 5S rRNAs, tRNAs, asRNAs, miRNAs and lncRNAs, possibly suggesting functional constraint. However, 173 protein coding genes were also included in this list.

### Mitochondrial Genome Variation

We identified 1,240 biallelic short nucleotide variants in the mitochondrial genome, consisting of 1,067 SNVs, 100 deletions and 73 insertions. A significant proportion (51.3%) of these variants were annotated as upstream gene variants, suggesting that regulatory regions contribute to the largest source of mitochondrial genetic variation. Coding variants contributed 47.6% of the total SNVs with 25.2% and 18.3% of the variants classified as missense and synonymous, respectively (Table S8A). High impact variants (frameshift, stop gained, stop lost and splice acceptor variants) were rare and only contributed 3.4% of the total SNVs identified in the mitochondrial genome.

We plotted the distribution of the 1,240 biallelic short nucleotide mitochondrial variants (Figure 1B). The insertions and deletions were largely in gene-rich regions, whereas SNPs were more evenly distributed across the mitochondrial genome. There was a significant excess of SNPs (366/4,259 *vs*. 701/14,198; Fisher’s Exact Test *P* = 8.48 × 10^-16^) and indels (113/4,512 *vs*. 60/14,839, Fisher’s Exact Test *P* = 2.16 × 10^-31^) in the AT-rich displacement loop (D-loop), non-coding control region of the mitochondrial genome (15kb – 19.5kb). These SNVs contributed to the predominance of upstream gene variant classification and the rarity of high-impact coding variants in the variant annotations.

To identify the mitochondrial haplotype groups in the DGRP3, we masked hypervariable regions, normalized the variants across the panel, and aligned them to conduct population genetics analyses. Variant site-only analysis produced a collection of 442 segregating sites with a nucleotide diversity (x) of 0.018 and 240 parsimony-informative sites. We visualized the parsimony network using PopART^72^ and clustered them to identify haplotype groups using the *Mclust* R package^73^. The PopART parsimony analysis identified 14 – 16 distinct haplogroups, with a single dominant haplotype group consisting of more than 400 members (Figure S3). The *Mclust* unsupervised cluster analysis identified 16 clusters, confirming the cluster membership distribution identified using the PopArt parsimony network. The most dominant cluster in the parsimony network (Figure S3) was cluster 14 in the *Mclust* results, with 463 members (Table S8B).

### Microbiome variation

Whole genome sequencing also yields information on the composition of microbiota present in and on the flies^48^. The presence/absence of the endosymbiont *Wolbachia pipientis* is polymorphic in *D. melanogaster* populations. We determined the *Wolbachia* infection status of each DGRP3 line by genotyping two *Wolbachia* genes and found that 66.3% of the DGRP3 lines tested positive for *Wolbachia* (Table S9A). The DGRP3 core microbiome is represented by all three kingdoms: Bacteria, Viruses and Eukarya (mostly yeast) (Figure S4, Table S9B, C).

Most of the community members are acetic acid bacteria (*Acetobacter* (51.85%), *Lactobacillus* (30.76%), *Komagataeibacter* (7.21%) and *Corynebacterium* (1.45%)) (Table S9C). Species of these commensal bacteria have been associated with positive effects on *D. melanogaster* fitness. *Acetobacter pomorum* modulates insulin and insulin-like signaling through acetic acid production^74^, promotes gut epithelial renewal and homeostasis^74^, has positive effects on lifespan and healthspan^74,75^ and modulates the immune system by interacting with immune deficiency signaling pathways^76^. *Lactobacillus plantarum* participates in the breakdown of complex sugars and polysaccharides into simple sugars^77^ and the acquisition of amino acids and vitamins^78,79^, promotes juvenile growth under nutrient-constrained conditions by modulating the insulin signaling pathway^80,81^, and has positive effects on lifespan and resistance to nutritional stresses^82^. *Komagataeibacter* is positively associated with egg-to-adult viability and resistance to heavy metal toxicity or environmental stress^83^. *Corynebacterium nuruki* affects thermoregulatory behavior^84^.

We used alpha diversity metrics to summarize the structure and complexity of the microbial community within each line. Shannon’s diversity^85^ captures both taxon richness (number of unique taxa) and abundance evenness (distribution of the taxa), Simpson’s D^86^ captures the degree of dominance by one or a few abundant taxa, and Simpson’s evenness^87^ (E) reflects how evenly abundance is distributed among observed taxa, with particular sensitivity to dominant members of the community. Across the DGRP3 population, these alpha diversity indices show a distribution of values rather than a single common profile, indicating that host lines vary in the complexity and dominance structure of their associated microbiomes (Figure S5). Lines with higher Shannon diversity were characterized by communities with greater richness and/or more even abundance distributions, whereas lines with higher Simpson’s D were more strongly dominated by a smaller number of microbial taxa. Variation in Simpson’s evenness further suggests that the balance among dominant microbial taxa differs across host genetic backgrounds.

We used beta diversity analyses to assess differences in community composition among lines. The Bray-Curtis dissimilarity metric^88^ quantifies pairwise differences in microbial abundance profiles, and Multidimensional Scaling (MDS) and Uniform Manifold Approximation and Projection (UMAP) visualizations of this metric revealed small groups of lines with similar microbial profiles (Figure S6). These patterns suggest that, in addition to variation in alpha diversity, subsets of lines share more similar microbiome compositions. Overall, these results indicate measurable heterogeneity in the Drosophila-associated microbiome across the population, both in terms of within-line diversity and between-line compositional structure.

We used the abundances of the microbial species to construct a Spearman rank-based co-occurrence network (Figure S7). The co-occurrence network revealed strong positive relationships between *A. pomorum* and multiple species of Lactobacillus (especially *L. plantarum*), and a strong negative relationship between *Lactobacillus fructivorans* and *Acetobacter senegalensis*. The strong positive association between *A. pomorum* and *Lactobacillus* is supported by the well-documented metabolic mutualism and syntrophy that exists between members of these clades of microorganisms in the *Drosophila* gut. *Lactobacillus* produces lactate from the *Drosophila* diet that feeds *Acetobacter,* and *Acetobacter* produces amino acids and vitamins that promotes the growth of *Lactobacillus* species^80,89^.

### Quantitative Genetic Analyses of Bristle Numbers

We quantified the numbers of bristles on the abdominal sternites (AB) and sternopleural plates (SB) bristles, classic *Drosophila* quantitative traits, for 1,124 DGRP3 lines (Table S10A, B). Our quantitative genetic analyses of these traits show that both have similarly high broad sense heritabilities (*H*^2^) for the total number of bristles (*H*^2^ = 0.677 for SB and *H*^2^ = 0.685 for AB) (Figure S8, Figure S9). The genetic and phenotypic correlations between the two bristle traits are low (∼0.17-0.18), albeit significantly different from zero (Table S10C). There is significant variation in sexual dimorphism for both traits (*i.e*., significant sex × line interaction variance), albeit of greater magnitude for AB than SB. Both AB and SB are spatially repeated characters, enabling us to test whether there is genetic variation in the magnitude of developmental homeostasis^52^ for these traits, estimated as variation among lines in the difference between the numbers of left and right SB or between the AB sternites. There is no significant genetic variation for developmental homeostasis for SB. However, the DGRP3 lines do harbor significant genetic variance for AB developmental homeostasis (*H*^2^ = 0.056), which is greater in females (*H*^2^ = 0.081) than males (*H*^2^ = 0.020) (Figure S10). The cross-sex genetic correlation (*r_GS_*) for AB developmental homeostasis is low: *r_GS_* = 0.190 (Table S10C). The estimates of environmental (residual within line) variance from the ANOVAs are pooled across all lines. However, several studies have shown that there can be genetic variance in within line (micro-environmental) variance^90–92^ for many quantitative traits. We find significant genetic variance in micro-environmental variance for total bristle number for both traits, which is greater for AB than SB, and greater for female than male AB (Table S10D).

The phenotypic correlation (*r_P_*) between micro-environmental variance (parameterized as in(*ln*(*σ_ε_*)) and the sum of SB number is positive and significant in both females (*r_P_* = 0.464, *P* = 4.31 × 10^-61^) and males (*r_P_* = 0.434, *P* = 7.01 × 10^-53^) (Table S10C, Figure S11). However, *r_P_* between in *ln*(*σ_ε_*) and the sum and difference of AB number are only significant for females: *r_P_* = – 0.249 (*P* = 2.41 × 10^-17^) for AB sum and *r_P_* = –0.267 (*P* = 7.59 × 10^-20^) for AB difference (Table S10C, Figure S11). Therefore, it appears that an “abnormal abdomen” phenotype is segregating in the DGRP3. This phenotype is characterized by large differences in the numbers of bristles on the two most posterior abdominal sternites (decreased developmental homeostasis) and increased micro-environmental variance of some lines that is more prevalent for lines with lower numbers of abdominal bristles. This phenotype has been observed repeatedly in *D. melanogaster* since its first description in 1952^93^, including in spontaneous mutation accumulation lines^22,23,94^ and among isogenic chromosome lines derived from natural populations^95^.

A total of 83 DGRP lines died after their numbers of bristles were quantified, giving a unique opportunity to assess the association of bristle numbers with reproductive fitness. Directional natural selection is associated with a change of mean and reduction of phenotypic variance following selection, while stabilizing selection is associated with reduced phenotypic variance after selection, but not a change of mean. We performed these analyses for differences between extant and extinct DGRP3 lines for AB and SB sum as well as in *ln*(*σ_ε_*) of AB and SB sum, separately for males and females (Table S10E-L). We observed selection for both AB and *ln*(*σ_ε_*) AB in females. The lines that died had lower mean bristle numbers (*t*_1,1116_ = 3.509, *P* = 0.00047) and higher among-line phenotypic variance (Levene’s test, *P* = 0.0125) than the extant lines; and higher micro-environmental variance (*ln*(*σ_ε_*)) (*t*_1,1116_ = −2.886, *P* = 0.004) and higher, but not formally significant, among-line phenotypic variance of micro-environmental variance (Levene’s test, *P* = 0.065). These observations are consistent with both directional and stabilizing natural selection against the abnormal abdomen phenotype in females. In males, the difference between the mean AB sum between the extant and extinct lines was in the same direction as for females, but not formally significant (*t*_1,1116_ = 1.867, *P* = 0.062), and the difference in among-line variance between the extinct and extant lines was not significant (Levene’s test, *P* = 0.172). However, lines with higher micro-environmental variance were also selected against (*t*_1,1116_ = −2.945, *P* = 0.0033) and the extinct lines had higher among line variance in micro-environmental variance than the extant lines (Levene’s test, *P* = 0.0021). Therefore, there appears to be directional and stabilizing selection for micro-environmental variance aspect of the abnormal abdomen phenotype in males. We found no evidence of selection for SB sum in either sex or *ln*(*σ_ε_*) SB in females, while there was directional selection against lines with higher male *ln*(*σ_ε_*) SB (*t*_1,1116_ = −2.366, *P* = 0.018).

### Genomic Relationships Among DGRP3 Lines

We performed Principal Components (PC) analyses to assess the extent of population structure in the DGRP3. We performed these analyses separately for the new DGRP3 lines, and for the subset of the re-inbred and re-sequenced DGRP2 lines that are part of the DGRP3. The first two PCs group the lines into three clusters that overlap between the DGRP3 and DGRP2 (Figure 3), indicating that this structure has been maintained for over 15 years in the outbred population from which the DGRP lines were derived. We performed a Random Forest machine-learning discriminant analysis to determine which variants separated the three sub-populations based on biallelic genotypes of 758,001 molecular polymorphisms (Figure S12, A-C), of which the top 216 nearly perfectly discriminated between the three sub-populations (Table S11A, Figure 12, D-F). The genomic distribution of these polymorphisms is within or near the breakpoints of several of the large polymorphic inversions (Table S11A, Figure S13). This is clear from examining the frequencies of the inversion karyotypes (Table S5) with the minor allele frequencies of the discriminating variants (Table S11A): the inverted karyotypes have lower frequencies than those of the discriminating variants. Therefore, these variants must also segregate in lines that are homozygous for the standard karyotype. Possibly, the inversions and discriminating variants are in inherently unstable genomic regions. However, the cause of the population structure is not known, even though it can be explained nearly perfectly.

**Figure 3.**
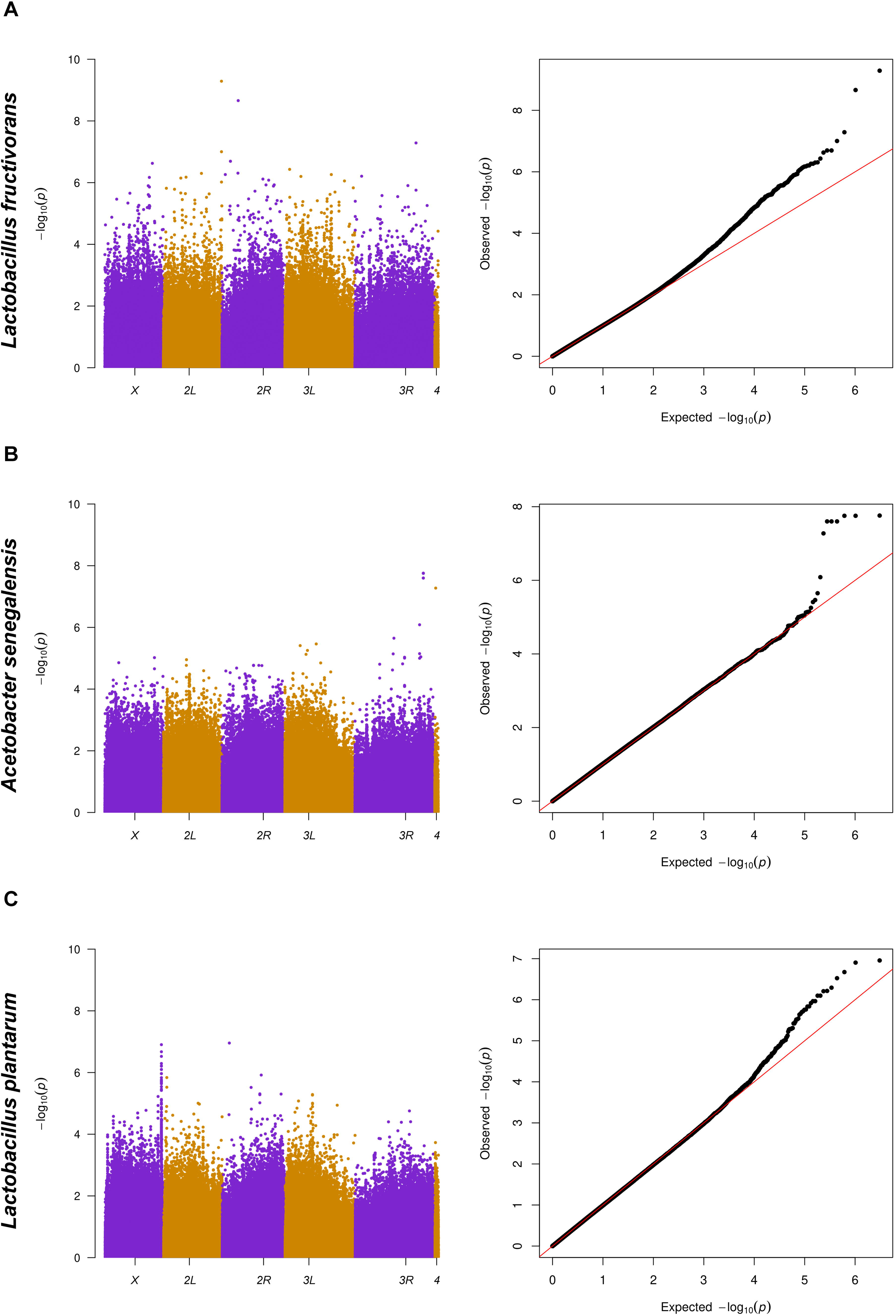
Principal Component analysis of structure in the DGRP3. The first two principal components from the analysis are depicted as scatter plots. Lines that were re-inbred and re-sequenced from the DGRP2 set are colored in purple, whereas the remaining DGRP3 lines are colored in orange.

We computed the genomic relationship matrix (GRM) between all pairs of DGRP3 lines^96,97^ (Figure 4A). The distribution of relatedness is skewed. Most lines have an average relatedness between −0.02 and 0.02, with the mode at zero. Therefore, most DGRP3 lines are unrelated, consistent with sampling from a large, random mating population and with the distribution of average pairwise relatedness observed in the DGRP2^45^. However, 71 pairs among 126 lines have a genomic relationship greater than 0.3 (Figure 4B, Table S11B). The lines with high genomic relatedness could be caused by sampling siblings from the natural population (many related pairs were collected on the same sampling date), shared karyotypes for genetically divergent inversions, or stock-keeping errors.

**Figure 4.**
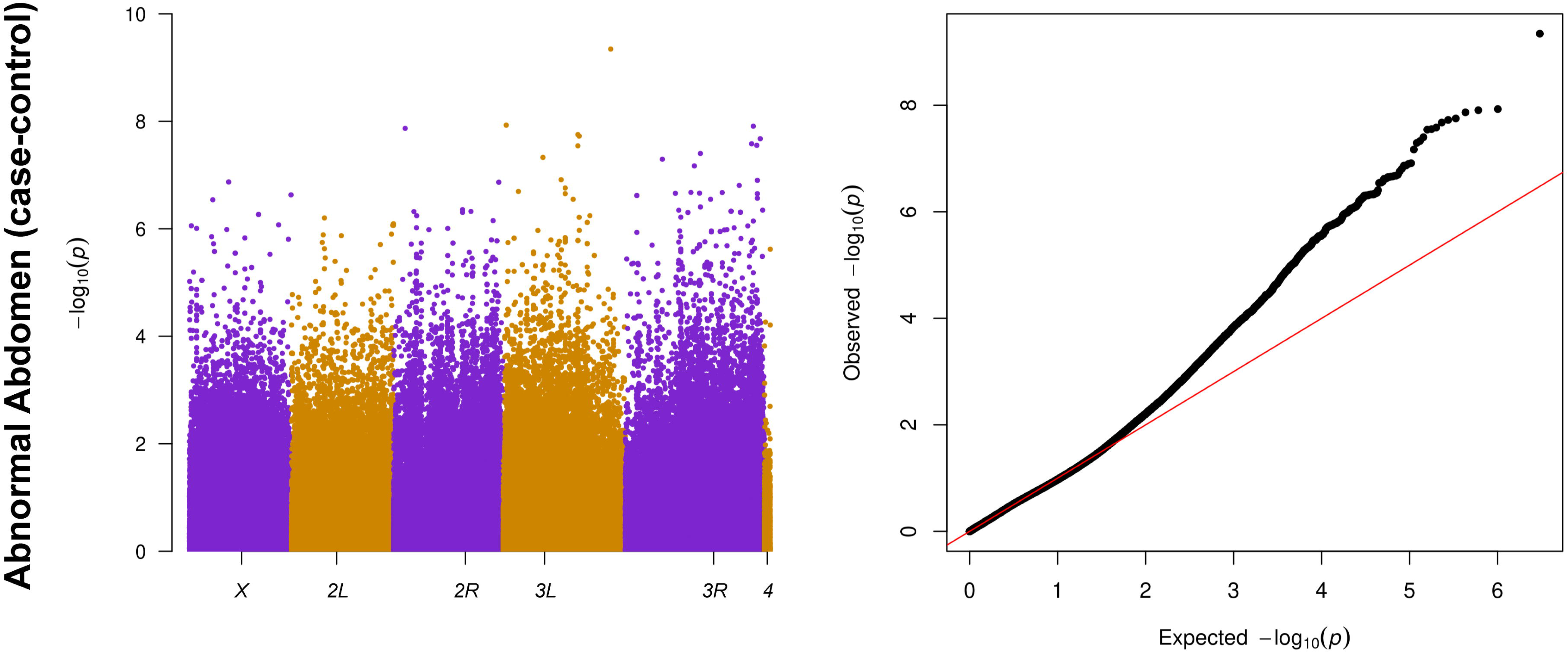
Genomic relationships among the DGRP3 lines. (A) Heat map of the bi-clustered pairwise relatedness among the DGRP3 lines. The diagonal shows the relationship of each line to itself and the symmetrical off-diagonals quantify the pairwise relatedness. (B) Histogram of pairwise relatedness. The insert magnifies the right tail (closely related lines) of the distribution.

The GRM indicates that there is variation in local LD across the genome. We computed the mean *r*^2^ between variants 50-150 bp apart in 100 kb sliding windows in each 1 Mb region for all major chromosome arms^98^ (Figure S14, Table S11C-G). Consistent with previous findings from smaller DGRP populations^44,45^, LD is small in regions of normal recombination and increases near centromeres and telomeres. In addition, there is greater LD on the *X* chromosome than autosomes, likely because the effective population size of the *X* chromosome is three quarters that of autosomes and there is greater purifying selection on the *X* chromosome. Removing the highly related lines would further decrease local LD.

### Genotype-Phenotype Associations

The DGRP3 is a community resource for GWA mapping of quantitative traits. We have developed a publicly available association mapping pipeline that uses line means as input and performs a linear mixed effects model that includes the GRM as a covariate to account for relatedness and population structure as well as covariates for *Wolbachia* infection status and large common inversions for which the *P*-value for association with the trait is < 0.1. We performed GWA analyses for variants affecting loss of DGRP3 lines during inbreeding, host control of microbiome composition, and numbers of abdominal and sternopleural bristles.

### Genomic signature of inbreeding depression

Inbreeding reduces the frequency of heterozygous genotypes with a concomitant increase in homozygous genotypes^52^ and reduces reproductive fitness by exposing recessive or partially recessive deleterious alleles in homozygous condition^52^. Inbreeding depression is high in *D. melanogaster*: only ∼35% of DGRP3 lines survived 20 generations of full-sib inbreeding. In addition, 196 DGRP3 lines died after they were sequenced. This gave us the opportunity to assess genetic features associated with inbreeding depression.

First, we asked whether the lines that died had more damaging variants than those that survived. We counted the number of homozygous high impact variants (frameshift, stop gained, stop lost, start lost, spice acceptor, splice donor and transcript ablation) in each extant and extinct line (Figure S15) and compared the two distributions. Indeed, lines that died harbored on average more homozygous high impact variants (613.5) than extant lines (564.5) (Welch’s *t*-test, 200 df; one-tail *P*-value = 0.013).

Second, we performed a GWA analysis using death status as a case-control phenotype to determine whether there were any individual alleles associated with inbreeding depression. We tested 1,501,062 variants with frequencies ≥ 0.01 for association with survival and identified 183 molecular polymorphisms in or near (± 1 kb) 117 genes at a nominal *P*-value < 10^-5^ (Table S12, Figure S16). Polymorphisms in two loci, *jim* (*3L*_22800626_A_G, *P* = 7.26 × 10^-11^) and *Nlg4* (*Neuroligin 4*) (*3R*_20243469_A_T, *P* = 2.51 x 10^-9^) were significant following a Bonferroni correction for multiple tests. *jim* encodes a zinc finger transcription factor involved in dendrite morphogenesis and chromatin silencing and *Nlg4* encodes a synaptic adhesion molecule involved in synapse formation and synaptic transmission^10^. Gene set enrichment analyses of all genes with association *P*-values < 10^-5^ showed enriched terms involved in nervous system development and function, implicating the nervous system as a major target in inbreeding depression.

### Host control of microbiome composition

*Drosophila* microbes can benefit the host by producing essential metabolites and nutrients^74,77–79^ and modulating insulin signaling^74,80,81^ and immune^76^ pathways. The host controls pathogenic microbes via Toll signaling^99^ and anti-microbial proteins^100^. We performed GWA analyses to identify variants and genes associated with global measures of microbial diversity and abundance of the most prevalent microbial species. We identified 275 polymorphic variants in or near (± 1 kb) 199 genes associated with alpha diversity (Simpson’s D statistic), and the abundance of six of the most common bacterial species (*Lactobacillus brevis*, *Komagataebacter hansenii*, *A. pomorum*, *L. fructivorans*, *A. senegalensis*, *L. plantarum*) at a nominal *P*-value < 10^-5^ (Table S13A,B; Figures 5, S17). Several molecular polymorphisms had association *P*-values less than the conservative Bonferroni-corrected *P*-value, including a polymorphism associated with *K. hansenii* abundance in a *Trim9* intron and six closely linked intergenic indel variants associated with *A. senegalis* abundance (Table S13A). There were no variants or genes in common between any of these analyses, suggesting that the host control of microbiome abundance is specific for each species. The 199 genes were not enriched for any Gene Ontology or pathway terms. However, many of the associated genes are attractive candidates based on known gene ontology categories (Table S13C). For example, the insect cuticle can confer resistance to pathogenic microbes^101^, and several cuticular genes (*CC-AP-R*, *Cht6*, *Cpr73D*) were implicated in the GWA analyses. *HBS1*, *kay* and *mop* are involved in immune/defense response to microbes; and *Act13E*, *Arms*, *b*, *Cda4*, *CG1486*, *CG99747*, *IP3K2*, *Pde8*, *Pdk* and *Ptp61F* are involved in various aspects of metabolism.

**Figure 5.**
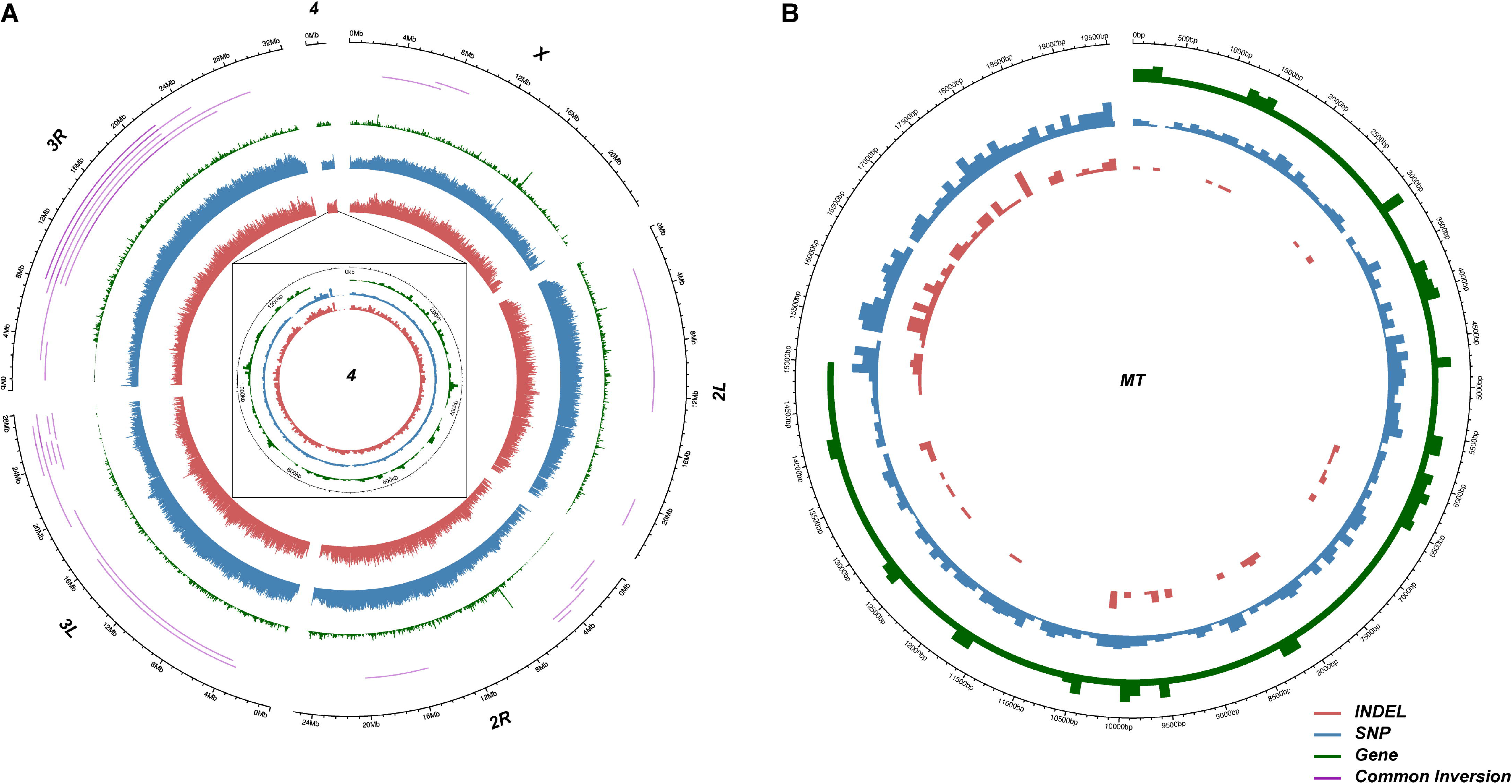
**GWA analyses of host control of microbiome composition**. The plots on the left are Manhattan plots of —*log*_10_(*P*) values, and on the right are quantile-quantile plots of observed and expected —*log*_10_ (*P*) values. (A) *Lactobacillus fructivorans*. (B) *Acetobacter senegalensis*. (C) *Lactobacillus plantarum*.

### Sensory bristle numbers

We performed GWA analyses for the total number of SB for males, females and the average of both sexes (Figure S18, Table S14). We did not perform GWA analyses for the micro-environmental variance of SB number because the Q-Q plots did not show enrichment of low *P*-values. We identified 110 molecular polymorphisms associated with these traits at *P* < 10^-5^, in or near (± 1 kb) 40 genes. An intronic SNP in *fred* was associated with all three analyses at a Bonferroni significance threshold (Table S14). Consistent with the high cross-sex genetic correlation for this trait, most associations are significant for males, females, and the average of the two sexes. Mechanosensory bristles are external organs of the peripheral nervous system and *fred* is involved in sensory organ development, which makes it a plausible candidate gene. The genes associated with total SB number are not enriched for any gene ontology terms or pathways; however, many are involved in peripheral nervous system development (*C901*, *hth*, *Lar*, *trol*) and/or nervous system function (*apt*, *hth*, *Lar*, *PiK59F*, *spin*, *trol*) (Table S14).

We also performed GWA analyses for the sum and difference of AB number on the two posterior sternites for males, females, the average of the two sexes and the difference between the sexes; and for micro-environmental variance^92^ of AB sum in males and females, parameterized as *ln*(*σ_ε_*) (Figures S19-S21, Table S15A,B). We identified 389 polymorphic variants in or near (± 1 kb) 240 genes associated (*P* < 10^-5^) with these phenotypes. Several polymorphic variants were significant following a Bonferroni correlation for multiple traits (Table S15A). Most polymorphisms were associated with either males or females, and only one SNP (*3L*_11955355_C_T) was associated with two traits: (*ln*(*σ_ε_*) AB in females and the difference between the sexes for the AB difference between the sternites. One gene (*Pdp1*) was common between the AB and SB GWA analyses, but the associated variants were not the same.

Several studies have identified genes affecting AB and SB numbers using transposable element-induced mutagenesis^102–104^, quantitative complementation tests^105^ and association mapping of naturally occurring variants in candidate genes known to affect bristle development and morphogenesis^39–41,43,106–108^ (Table S16). Of these 194 genes, 11 (*Atf3*, *Bx*, *cpo*, *crol*, *heca*, *l(3)05822*, *l(3)07882*, *Ptp10D*, *tai*, *ttk*, *tyn*) were identified in this study.

The genes associated with any of the AB traits were enriched for general Gene Ontology categories. One more specific category, mechanosensory behavior (GO:0007638), makes sense given that bristles are mechanosensory organs (Table S15C). Of the 240 genes associated with AB number traits, only 45 (18.8%) were associated with nervous system function and/or development and could plausibly have pleiotropic effects on bristle number (Table S15D). Several of these genes affect peripheral nervous system development (*abd-A*, *ttk*), chaeta development or morphogenesis (*Bx*, *chm*, *Cul4*, *fry*, *kirre*, *msi*, *WASp*), sensory organ development (*chm*, *E(spl)m4-BFM*, *WASp*) and mechanosensory behavior or response to mechanical stimulus (*jus*, *Ptp61F*, *Pzl*, *TrpA1*). *Ptp61F* is thus a top candidate gene given both the significance of the association and known function. However, the other top candidate genes based on the strength of the association, *CG4374* and *CG32816*, regulate transcription and have no annotated function, respectively; and the other top variants are intergenic.

Given the segregation of a female-specific abnormal abdomen-like phenotype in the DGRP3, we also performed a GWA analyses for this phenotype. We considered lines to have this phenotype if the mean AB sum was less than 30, and/or the mean AB difference was less than −5, and/or AB *ln*(*σ_ε_*) was greater than 1.8 (Table S17). We identified 317 unique variants in or near (± 1 kb) 184 unique genes at a nominal *P*-value < 10^-5^ (Figure 6). Several of these variants had *P*-values below or near the Bonferroni-corrected significance threshold, including variants in *alpha-Man-1b*, *CG2310*, *CG11317*, *CG46525*, *Diedel*, *ebd1*, *heca*, *Pdhb*, *Rbp6*, *scro*, *SecC1* and *UQCR-Q*. None of these genes is an obvious candidate affecting this phenotype based on their known functions and expression patterns^10^. However, 31 genes associated with the abnormal abdomen phenotype from the abnormal abdomen GWA analysis overlapped with the GWA analyses of AB number phenotypes. A total of 21 of these genes had female-specific effects on AB number (*asRNA:CR5000*, *ATPsynbetaL*, *CG32816*, *CG4374*, *Cic-C*, *heca*, *Lcch3*, *Rbp6*), *ln*(*σ_ε_*) AB number (*CG32637*, *plum*, *sli*), the difference between males and females in AB number (*CG32637*, *CG32816*, *CG42788*, *CG5742*, *fru*, *Lpin*, *Myo81*), and the difference in AB number between the two most posterior sternites (*dah*, *iab8/MSAmiP*, *Octalpha2R*, *Pzl*). The abnormal abdomen phenotype is clearly polygenic, consistent with previous observations that it occurs on both Chromosome *2* and *3* extraction lines derived from the same Raleigh population^95^.

**Figure 6.**
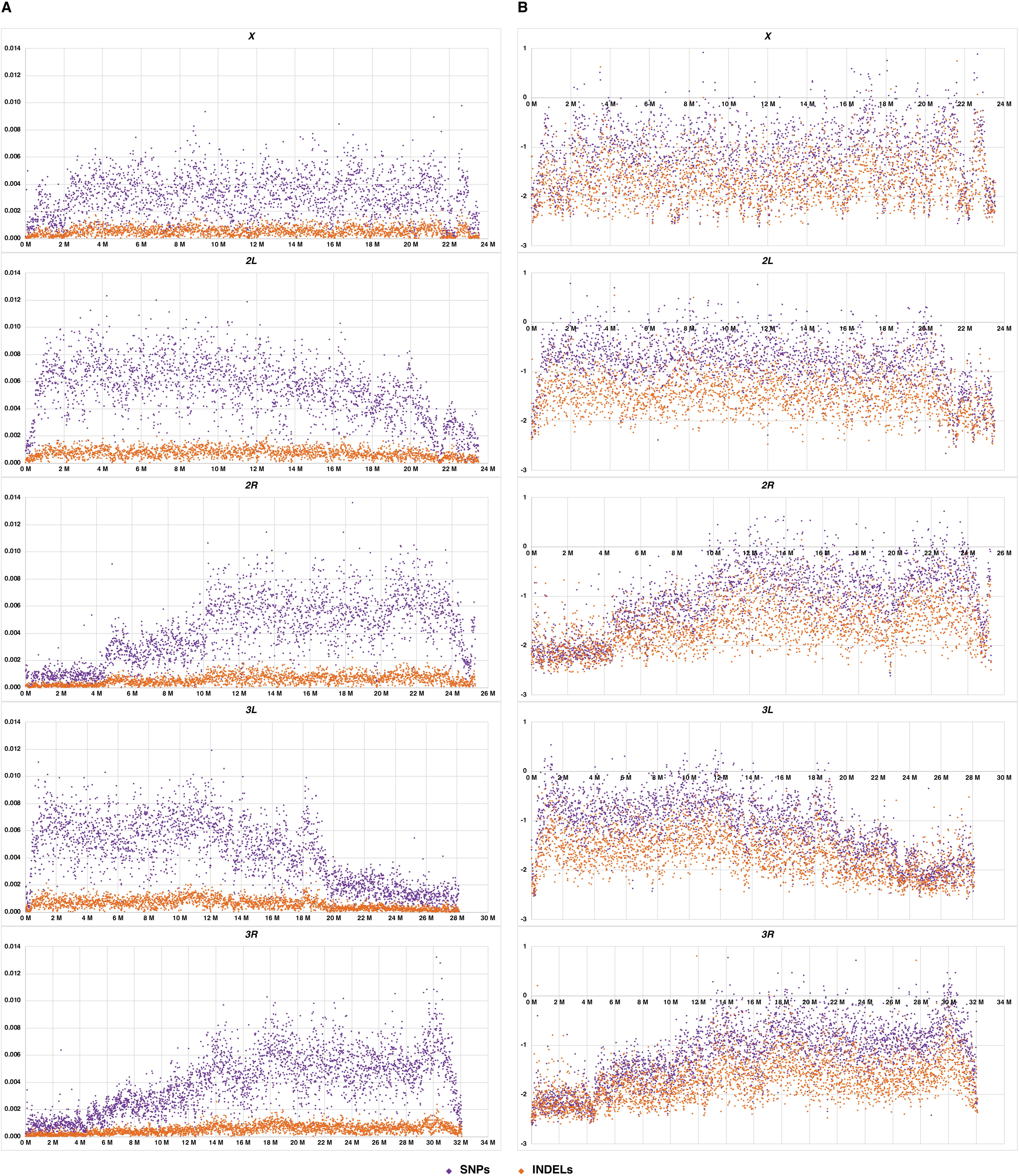
GWA analysis of the Abnormal Abdomen phenotype in females. (A) Manhattan plot of —*log*_10_ (*P*) values. (B) Quantile-quantile plot of observed and expected —*log*_10_(*P*) values.

### Statistical power to detect associations

The substantially increased number of lines in DGRP3 provides significantly higher statistical power to detect genotype-phenotype associations, especially for variants with smaller effects and lower allele frequencies (Figure S22). For example, for a low frequency (MAF = 0.01) variant with a very large effect of one standard deviation, the power to detect this association at a *P*-value threshold of 10^-5^ is 0.8% when there are 200 lines (Figure S22A). At the same *P-*value threshold, the power to detect this association increases to 68% when there are 1,200 lines (Figure S22B).

To demonstrate the increased power in the DGRP3, we performed GWA mapping for female abdominal bristle number, but sub-sampled the full dataset with increasing sample size from 200 to 1,124. It is important to note that this analysis represented only one random subsample, and we ensured that the smaller samples were subsets of larger samples. Among the 42 variants that were significant for the full DGRP3 (*n* = 1124), two were first discovered at a sample size of 200 lines, 19 at 500, 11 at 1000, and 10 for the full dataset (Figure S23A).

Among the 18 genes significant in the full data set, one was first discovered at 200, three at 500, four at 1,000 and 10 at 1,124 (Figure S23B). As sample size increases, a variant may become significant because its MAF passes the threshold for inclusion in association tests and/or increased power at larger sample sizes. This result demonstrates that larger sample sizes validated variants or genes discovered at smaller sample size, likely due to their large effect sizes. More importantly, larger sample sized discovered new associations that were elusive at smaller sample sizes.

## Discussion

The original DGRP2 has been a valuable community resource that has complemented the extensive *D. melanogaster* toolbox by enabling GWA analyses in this widely used model organism. Here, we report the DGRP3, a five-fold expansion of the DGRP2. Given strong orthology between *Drosophila* and human genes and evolutionary conservation of fundamental biological processes, the DGRP3 will be a valuable resource for comparative genomic studies on complex traits. The DGRP lines are available from the Bloomington Drosophila Stock Center and the sequence data are available from the short-read archive of the National Center for Biotechnology Information (accession numbers are given in Table S1). The new DGRP3 website (www.flydgrp.org) hosts the mixed linear model pipeline for GWA analyses using DGRP3 line mean phenotypes that account for any effects of *Wolbachia* infection, large polymorphic inversions and cryptic relatedness among the DGRP3 lines. This pipeline returns the results of all associations less than a nominal *P*-value of 0.001 for allele frequencies ≥ 0.01, complete annotations for each affected transcript, the covariate analyses, figures showing Manhattan and Q-Q plots, and guidance for best practices for reporting GWA results. The website also hosts the phenotype data described in this study. These data and the pipeline will be useful for teaching principles of quantitative genetics.

### Catalog of Naturally Occurring Molecular Polymorphisms

The DGRP3 harbors 42.4% more molecular polymorphisms than the DGRP2 (8,760,339 *vs*. 6,149,882). These variants complement the efforts of the Berkeley *Drosophila* Genome Project Gene Disruption Project^3,4^ to generate mutations in all *D. melanogaster* genes for functional annotation and the *D. melanogaster* modENCODE Project^109^ to identify sequence-based regulatory elements in this species. The naturally occurring polymorphisms in the DGRP represent different functional classes than transposon insertions used by the Gene Disruption Project to generate mutations and represent the spectrum of natural variants in all populations, including humans. We also called short (≤ 50 pb) insertions and deletions, as well as large (> 50 bp) structural variants (insertions, deletions, tandem duplications, intrachromosomal translocations, inversions). All DGRP3 lines contain at least one gene with a loss of function mutation, and there are segregating loss of function mutations in ∼one third of the *Drosophila* coding genome (4,550/14,000 = 32.5%). In addition to nuclear polymorphisms, we genotyped molecular polymorphisms in the mitochondrial genomes and mitochondrial haplotypes, and the composition of microbiota in and on the flies. These features can also be incorporated in future GWA analyses for relevant phenotypes as well as population genetic analyses.

The population structure previously observed in the DGRP2^45^ persists in the DGRP3. It is intriguing that most of the variants that discriminate the three sub-populations are in the same genomic regions as the segregating large inversions and near their breakpoints; however, the inversions themselves are not associated with this structure. Although a Random Forest classifier based on 216 polymorphic variants nearly perfectly discriminates these sub-populations, the underlying cause of the structure remains unknown.

### DGRP3 GWA Analyses

The DGRP3 is five-fold larger than the DGRP2 and thus provides greater power for GWA analyses as well as the ability to determine genotype-phenotype associations for low frequency alleles (MAF ≥ 0.01). Previous DGRP GWA analyses have identified novel genes associated with quantitative traits that defined the first functional annotations of predicted genes and revealed unexpected pleiotropic effects of genes with known roles in development, physiology and/or metabolism; and showed that most associations were in non-coding genomic regions.

Our GWA analyses of inbreeding depression, microbiome composition and numbers of sensory bristles have the same properties. The 71 pairs of related lines in the DGRP3 offer the opportunity for reduced complexity mapping of quantitative traits^110^. If a phenotype is statistically different between pairs of related lines, then the search for associated variants can be restricted to the regions where the related lines have different sequences, in the absence of epistasis.

An advantage of the DGRP3 is that the population is large enough for researchers to measure their trait of interest on a subset of DGRP3 lines and hold out the remaining lines for functional validation of variant associations. Previously, functional tests were largely restricted to knocking down expression of candidate genes using RNA interference (RNAi) to assess the effect(s) on phenotypes. However, RNAi can have unknown off-target effects; a reduction in the amount of transcript may not be the mechanism by which the naturally occurring polymorphisms exert their phenotypic effects; we rarely know in what tissue modulating gene expression is relevant to the trait of interest; and publicly available RNAi constructs do not enable testing of intergenic polymorphisms. Independent validation of effects of molecular polymorphisms in additional samples from the same population is the gold standard. We note that LD declines rapidly with physical distance in the DGRP3, as in the previous DGRP populations^44,45^, but we still observe short haplotypes of associated polymorphisms, restricting functional tests to these local haplotypes. In addition, a single molecular polymorphism can affect multiple overlapping genes as well as multiple transcripts of the same gene. In either of these situations, additional experimental studies will be needed to determine the gene, transcript and relevant tissue responsible for the phenotypic effects.

### Sensory Bristle Numbers

The numbers of sensory bristles on the sternopleural plates and abdominal sternites are classical *Drosophila* quantitative traits. Bristle numbers have been used to test quantitative genetics theory^11–15^, estimate the spontaneous mutation rate for quantitative traits^20–23^, map QTLs^32–37^ and polymorphic variants in candidate genes^39–41,43,108^, and identify novel candidate genes using *P*-element insertional mutagenesis^104,111,112^. Our quantitative genetic analyses of bristle numbers in the DGRP3 reveal that SB and AB have similarly high broad sense heritabilities, but otherwise different genetic architectures. There is genetic variation for developmental homeostasis for AB, but not SB number. Although there is genetic variation for micro-environmental variance for both AB and SB, the magnitude is much greater for AB. These differences are attributable to an abnormal abdomen-like phenotype that segregates in the DGRP3, which is associated with low bristle numbers, high micro-environmental variance and reduced developmental homeostasis for AB.

The question of the relationship between bristle numbers and fitness has been contentious, with some studies claiming strong stabilizing selection^113^ and others claiming neutrality^114–116^. Our analyses of the differences in mean and variance of bristle numbers between lines that died and extant lines indicate directional selection against low AB number and high micro-environmental variance of AB number, as well as stabilizing selection for AB number in females. In males, there is directional selection against high micro-environmental variance as well as stabilizing selection for micro-environmental variance of AB number. SB number appears to be neutral, with some evidence for directional selection against high micro-environmental variance for SB number in males.

Our GWA analyses identified 281 candidate variants and genes associated with bristle traits, only 11 of which were among the 194 genes previously shown to affect bristle numbers. This is not surprising for *P* transposable element insertional mutations, which tend to have larger effects than naturally occurring variants and for which saturation mutagenesis has not been done. However, naturally occurring variants in *achaete*^39,107^, *scute*^39,107^, *scabrous*^40^, *Delta*^41,106^, *hairy*^108^ and *Catsup*^43^ have been associated with numbers of AB and SB, and these genes were not detected in the DGRP3 GWA analyses. The previous association studies used chromosome substitution lines (in which chromosomes from the same Raleigh, NC natural population used to derive the DGRP lines that contained the relevant candidate gene were individually substituted into a common co-isogenic background) or near-isoallelic lines (in which a genomic segment containing candidate gene from each extracted chromosome was further introgressed into the isogenic background genotype). The combined effect of a panel of chromosome substitution lines on quantitative traits is commonly much greater than the difference in the phenotypes between the parental strains, indicating the prevalence of suppressing epistasis for the trait in the parental strains^117,118^. This phenomenon could be the reason for the difference between the GWA results for bristle numbers in the DGRP3 and those from previous studies.

## Methods

### *Drosophila* Maintenance

We maintained all fly stocks at 25°C, 70% humidly, 12:12 hour light-dark cycle on cornmeal-agar-molasses medium (Bloomington Formulation, Nutri-Fly™, #66-112, Genesee Scientific) in 28.5 mm wide polypropylene vials (Genesee Scientific Cat. #32-120). Back-up cultures were maintained at 18°C.

### DGRP3 Lines

We collected flies from peaches using a *Drosophila* sweep net between 6:30 am and 7:30 am at the State Farmers Market in Raleigh, North Carolina (35° 45’ 49.086’’ N, 78° 39’ 45.7488’’ W) from May-September 2015, June-August 2016 and August-September 2017. We placed single inseminated wild-caught females on culture medium to produce progeny, and distinguished *D. melanogaster* females from *D. simulans* females by the morphology of the external genitalia of their male progeny. *D. melanogaster* inbred lines were created by strict full-sib single pair inbreeding for 20 generations. Five single pair matings were set up each generation for each inbred line; males and females from one of these single pair matings were chosen to establish the next generation of inbreeding. Of the ∼3,000 isofemale lines, 1,058 survived the initial inbreeding.

The DGRP2 flies have been maintained by mass mating in vials since 1998-2003 and sequenced between 2010 and 2013^44,45^. The original genotypes are likely to have changed since the initial sequencing due to accumulation of new mutations and fixation of heterozygous variants due to drift. Therefore, we re-inbred the DGRP2 lines for five generations of strict full sib mating. Of the 197 extant DGRP2 lines, 175 survived the inbreeding and are included among the DGRP3 lines, for a total of 1,233 initial DGRP3 lines. To distinguish the DGRP3 from earlier DGRP lines, they are named RP-XXXX, while the DGRP2 lines are named DGRP-XXX. A total of 196 of the original lines were lost due to poor fitness; currently, there are 1,037 DGRP3 lines from the 2015-2017 collection (Table S1_DGRP3 lines).

### Whole Genome DNA Sequences

We extracted DNA from whole flies from all 1,233 DGRP3 lines using the Zymo Research Quick-DNA Tissue/Insect Miniprep Kit and quantified the DNA using the Invitrogen Qubit 4 Fluorometer and Thermo Scientific NanoDropTM 8000 Spectrophotometer. We prepared DNA libraries using a modified NEBNext ® Ultra DNA Library Prep protocol and quantified the libraries using the Invitrogen Qubit 4 Fluorometer and TapeStation 4150 System. We sequenced all samples to ∼40X coverage using Illumina NovaSeq 6000 S1 Reagent Kits v1.5 and 2 x 150 chemistry (300 cycles).

### Variant Calling and Annotation

We filtered the raw sequencing data to remove short reads and low-quality reads and bases and removed adapter sequences using the *fastp* pipeline^119^; and aligned high-quality sequence reads to the *D. melanogaster* reference genome (NCBI, release 6.13) with Burrows-Wheeler Aligner (BWA) v0.7.17^120^. The alignments were sorted and indexed using Samtools (v1.10)^121^ and locally realigned and marked for PCR duplicates with Genome Analysis Toolkit (GATK) v4.1 and Picard tools v2.21.7 before recalibrating base qualities with GATK^122^. We performed variant calling for each individual line using the GATK *haplotypecaller* with the GVCF flag to generate genomic variant call format (VCF) files. We used the GATK joint variant calling workflow with default parameters for joint calling of individual genomic VCF files to generate a combined VCF file, and the Joint Genotyping for Inbred Lines (JGIL) workflow^123^ to account for expected homozygosity following 20 generations of full-sib inbreeding. We separated the resulting variants into SNPs and small (< 50 bp) insertion/deletion molecular polymorphisms and removed low-quality variants using standard filtering criteria recommended by the Broad Institute (GATK Team). For genome-wide association analysis, we filtered the VCF file containing high-quality variants to keep biallelic sites and remove monomorphic and segregating sites and converted the filtered, combined VCF file into PLINK v1.9 format^124^. We annotated the short variants using Variant Effect Predictor (VEP) with Ensembl database v111^125^. To summarize the variant effects, we used the *--pick* option to select a single annotation for each variant based on the standard set of ordered criteria in VEP and the *D. melanogaster* Ensemble R6 version of the reference genome.

We identified large (> 50 bp) structural variants (insertions, deletions, duplications, inversions, translocations) using the processed alignment files as input for MANTA v1.6.0^126^. We converted the break-end coordinates for structural variants using the *convertInversion.py* python script within the MANTA workflow. Individual batches of structural variant VCF files were split into single-sample VCF files and then merged using SURVIVOR (v1.0.7)^127^ with a 1 kb margin for breakpoints and single aligner criteria. We annotated large structural variants by identifying overlap between coordinates of variants and known genomic features which include genes, repeat regions and estimated cytogenetic bands retrieved from the FlyBase database^10^.

We used Block Resolution and Annotation of Integrated DNA (BRAID, https://github.com/YuefanHuang1998/BRAID) to identify DGRP3 lines where the combined effects of variants in each protein coding gene result in loss-of-function (LoF). We defined LoF effects as those that lead to a greater than 10% change in the polypeptide sequence in all splice isoforms. Briefly, we processed the reference sequence and annotation (Flybase 6.46) into DNA blocks corresponding to genes. We read the genotype data in VCF format based on their overlap with the blocks and considered only homozygous variants. We combined alleles in the same line (thus on the same haplotype) to obtain mRNA sequences, which we translated *in silico*. We compared the variant protein sequence with the reference protein sequence and considered divergence greater than 10% between the two protein sequences as LoF.

We represented the distribution of genes, SNPs, insertion/deletion polymorphisms and common polymorphic inversions as histograms across all six chromosome arms by binning frequencies within 50 kb windows and circularizing using the *circlize* R package^128^.

### Mitochondrial Haplotypes

We filtered multi-sample VCF files specifically for the mitochondrial chromosome, normalized to avoid overlapping variants using the bcftools *norm* function, and produced per-sample consensus FASTA sequence using the bcftools *consensus* function^129^. We aligned the per-sample FASTAs using the MAFFT multiple sequence aligner^130^ prior to phylogenetic analysis and masked the aligned FASTA sequences for hypervariable regions^131^. We analyzed the masked, normalized and aligned FASTA file to construct parsimony networks using PopART^72^ and clustered them to identify haplotype groups using the *Mclust* R package^73^. We identified the number of haplotype groups using the Bayesian Information Criterion (BIC) under different covariance model conditions.

### *Wolbachia* Infection Status

We used alignment files to identify the presence of the endosymbiont, *Wolbachia pipientis*. Reads that did not align to the *D. melanogaster* reference genome were aligned to two canonical *Wolbachia*-specific genes (*Wolbachia Surface Protein* and *wAu* ankyrin domain protein) using bbmap aligner v38.73^132^. We used the successful alignment of more than 7,000 reads as the criterion for positive infection status. The threshold was determined by constructing a frequency histogram for the number of aligned reads for the two indicator genes across the DGRP3 population and identifying a global minimum.

### Microbiome Profiling

We filtered alignment files for sequences that did not align to the *D. melanogaster* reference genome and created FASTQ files using the Samtools subcommand *fastq* (v1.10)^121^. We analyzed these unaligned FASTQ files to generate quantitative microbiome community data using the YAMP pipeline^133^ which employs MetaPhlAn2^134^ for taxonomic binning. We aggregated the relative abundance data across all samples using the standard python *pandas* library^135^. We defined the core microbiome by filtering for genera that were present in at least 90% of the DGRP3 lines. We created a sunburst plot using core microbiome relative abundances using the python *plotly* library^136^. We constructed a co-occurrence network by calculating Spearman rank correlations at the species level. We adjusted the correlation *P*-values for multiple hypothesis testing using the Benjamini-Hochberg False Discovery Rate (FDR)^137^. We visualized the co-occurrence network using Cytoscape^138^ and removed edges with FDR >0.05. The *scipy*^139^, *skbio*^140^ and *sklearn*^141^ python libraries were used to calculate alpha and beta diversity metrics. We calculated Multidimensional Scaling (MDS) and Uniform Manifold Approximation and Projection (UMAP) plots and constructed using *sklearn* and *seaborn*^142^ python libraries, respectively.

We computed Shannon’s *H*^85^ (*H* = — ∑*_i_ p_i_ ln*(*p*_*i*_)), Simpson’s *D*^86^ (*D* = ∑*_i_ p_i_* ^2^) and Simpson’s E^87^ (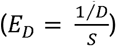) indices of alpha diversity, where S is the number of taxa. We also computed the *S* Bray-Curtis dissimilarity distance metric^88^ 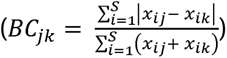 as a measure of beta diversity, where, *x_ij_* is the abundance of taxon i in DGRP3 line *j, x_ik_* is the abundance of taxon i in DGRP3 line k and S is the total number of taxa considered across each pair of lines.

### Population Structure

To estimate population structure within the DGRP3, we imported the filtered VCF file into the *SNPRelate* R package^143^. We estimated LD using the *r*^2^ parameter^98^ and computed the mean *r*^2^ between variants 50-150 bp apart in 100 kb sliding windows in each 1 Mb region for all major chromosome arms. We pruned the SNP data generated from the VCF file for LD using the *snpgdsLDpruning* function and an *r*^2^ threshold of 0.2. We performed Principal Components Analysis (PCA) of the LD-pruned SNP data using the *snpgdsPCA* function. We used the Random Forest machine learning classifier *randomForest* R package^144^ to build a classification model separating the three populations identified in the PCA. We used an elbow plot constructed using variable importance (Mean Decrease in Accuracy) scores from the Random Forest model to identify model-informative SNPs. We calculated an additive kinship matrix using the R package *rrBLUP*^145^ after recoding standard allele-count based genotype code to −1, 0 and 1 and scaling the resulting matrix by dividing by 2. We considered pairs of lines with values greater than 0.3 to be closely related. We visualized the scaled kinship matrix as a heatmap using Genesis software^146^.

### Numbers of sensory bristles

We counted the number of sternopleural bristles (SB) on the left and right sternopleural plates, and the number of abdominal bristles (AB) on the 5^th^ and 6^th^ abdominal sternites in females, and the 4^th^ and 5^th^ abdominal sternites in males. Bristle numbers were recorded for 10 males and 10 females for each of 1,124 DGRP3 lines. We performed factorial mixed effect analyses of variance (ANOVAs) of form *Y* = *μ* + *S* + *L* + *S* x *L* + *ε*, where Y is the phenotype, y is the overall mean, S is the main fixed effect of sex, L is the main random effects of DGRP3 line, S x L is the random sex by line interaction, and z is the residual. We performed these analyses for the sum and the difference of the numbers of sternopleural bristles on the left and right sternopleural plates, and the sums and differences of abdominal bristle numbers on the two most posterior abdominal sternites. We also performed reduced models (*Y* = *μ* + *L* + *ε*) separately for females and males. We estimated the broad sense heritability for each trait as 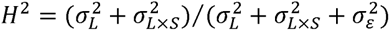, where *σ*^2^ denotes a variance component and the subscripts indicate the random effect terms. We calculated the cross-sex genetic correlations for each trait as *r_GS_* = *Cov_F,M_*/(*σ_LF_σ_LM_*), where *Cov_F,M_* is the product-moment covariance of male and female line means, and p_LF_ and p_LM_ are the standard deviations of the among line variance components from the female and male reduced models, respectively^52^. We also computed the cross-trait genetic correlations, separately for males and females, as *r_GSB,AB_* = *Cov_SB,AB_*/ (*σ_LSB_ σ_LAB_*), where *Cov_SB,AB_* is the product-moment covariance of line means for the two traits, and *σ_LSB_* and *σ_LAB_* are, respectively, the standard deviations of the among line variance components for sternopleural and abdominal bristle number^52^. ANOVAs and estimates of variance components were performed using the GLM command (Type III) in SAS™ Studio v3.8 software^147^.

We assessed whether there is genetic variance of micro-environmental variance^90–92^ for the bristle traits. We computed the within line variance for each DGRP3 line for each of the bristle traits and used Levene’s and Brown-Forsythe tests to determine whether there is significant heterogeneity of micro-environmental variance among the lines. We used *ln(σ_ε_*) to quantify micro-environmental variation for each line for further analyses of micro-environmental variance because it is perfectly correlated with in (*ln(σ^2^_ε_*), which has desirable statistical properties and has been used previously to quantify micro-environmental variance^90–92^, and *ln(σ_ε_*) scale as trait means.

### Genome-Wide Association Pipeline

We developed a pipeline for genome wide association (GWA) mapping of line mean phenotypes across the DGRP3. The pipeline first evaluates the effects of *Wolbachia* infection status and large common inversions (minor allele frequency > 0.03). The pipeline preprocesses the PLINK files and covariate files to only include DGRP3 lines for which the trait was measured, and filters them for sites based on a genotype missing rate of 0.1 and a minor allele frequency ≥ 0.01. The pipeline also determines which covariates to include in the linear mixed effects models (LMM) for association testing by fitting a generalized linear model for each covariate against the phenotype. Only covariates with *P*-values less than 0.1 are included in the LMM for the GWA analysis. The Genomic Relationship Matrix (GRM), calculated using GEMMA v0.98.5^148^, is included as a covariate to account for relatedness and population structure. The GWA analyses are performed using GEMMA, and statistical significance is calculated using the Wald test. The β coefficients from the Wald test are estimates of additive effects, calculated as half the difference between means of individuals with the major and minor allele genotypes^52^. The pipeline generates quantile-quantile (Q-Q) plots and Manhattan plots in R using the qqman library^149^. The effects of associated variants are annotated using VEP v111^125^. All annotations for each variant are reported.

We performed GWA analyses for variants affecting loss of DGRP3 lines during inbreeding, host control of microbiome composition, and numbers of abdominal and sternopleural bristles, using the line means for each trait. We used Simpson’s D values and taxa with average relative abundance greater than 5% as phenotypes for the host control of microbiome composition GWA analyses. The relative abundances of these taxa were centered-log ratio transformed^150^ to correct for constant sum constraint^151^ prior to performing GWA testing.

We coded binomial traits as 0,1 for GWA analyses. We only performed the GWA analyses for traits for which the Q-Q plots indicated enrichment of true positives above the nominal —log_10_*P*-value threshold of 5.

## Data Access

The DGRP3 lines are available from the Bloomington Indiana Drosophila Stock Center. The raw genome sequencing data generated in this study have been submitted to the NCBI Short Read Archive and the BioProject database (https://www.ncbi.nlm.nih.gov/bioproject/) under accession number PRJNA1139241. The reviewer link is ftp://ftp-trace.ncbi.nlm.nih.gov/sra/review/SRP521783_20251201_172340_eac2256ff0666fc7e5492d2713c8b229. Individual DGRP3 accession numbers are given in Table S1. All other data generated in this study are given in the supplementary tables and on the DGRP3 website (www.flydgrp.org). The GWA analyses pipeline is hosted on the DGRP3 website. All code is given in the github repositories https://github.com/vshanka23/dgrp_gwas_final and https://github.com/YuefanHuang1998/BRAID (BRAID).

## Acknowledgements

This study was funded by National Institutes of Health grants R01 AG073181 and U01 DA041613 to T.F.C.M. and R.R.H.A. The Clemson University Institute for Human Genetics Statistical and Bioinformatics and Genomics Research Cores were funded in part by P20 GM139769 to T.F.C.M. and R.R.H.A. R.F.L. generated the DGRP3 lines. R.F.L., R.L.L., and N.O.N.Y. maintained the DGRP lines. R.A.L. and L.S. sequenced the DGRP3 lines. V.S., M.E.A., J.B., Y.H., W.H. and T.F.C.M. analyzed the data. V.S., W.H., R.R.H.A. and T.F.C.M. wrote the manuscript. We acknowledge the technical assistance of Dr. Miyoung Shin and Mr. Brian Kesser.

## Supplementary Figures

**Figure S1. Minor allele frequency (MAF) distributions.** MAF frequency distributions are shown for all polymorphic molecular variant categories. Small insertions and deletions are ≤ 50 bp, large structural variants are > 50 bp. (A) SNPs. (B) Small insertions. (C) Small deletions. (D) Large insertions. (E) Large deletions. (F) Large tandem duplications. (G) Large inversions. (H) Large interchromosomal translocations. Inserts magnify the right tails (common variants) of the distributions.

**Figure S2. Distributions of Loss of Function (LoF) genes.** (A) Frequency histogram of the number of LoF genes per DGRP3 line. (B) Frequency histogram of the number of DGRP3 lines harboring LoF genes.

**Figure S3. Mitochondrial haplotypes.** The 16 major mitochondrial haplotypes (black circles) are depicted, as well as the DGRP3 line composition of each haplotype. Orange circles represent the haplotypes and the size of the circles represents the number of lines within each of the haplogroups. The orthogonal notches on the edges represent the number of variants separating the nodes (variant distance). Ellipses encompass large clusters of haplogroups and singlet haplotypes within one edge.

**Figure S4. Composition of the core DGRP3 microbiome.**

**Figure S5. Measures of microbiome diversity and richness**. (A) Shannon’s D^85^. (B) Simpson’s D^86^. (C) Simpson’s E^87^. Vertical lines in the violin plots represent the 25^th^, 50^th^ and 75^th^ quartiles of the distribution.

**Figure S6. Microbiome beta diversity.** (A) Bray-Curtis^88^ multidimensional scaling (MDS). (B) Uniform Manifold Approximation and Projection (UMAP). Ellipses in each plot represent dense clusters of DGRP3 lines.

**Figure S7. Microbiome co-occurrence network.** The nodes (orange circles) denote major microbial species, and the edges represent the Spearman rank correlation coefficients between the abundances of the indicated species. Blue edges indicate negative correlations, and red edges show position correlations, with the magnitude of the correlations given by the scale bar.

**Figure S8. Distributions of the sum of the numbers of bristles on the left and right sternopleural plates.** (A) Distribution of male line means, with lines ordered by increasing mean. (B) Frequency distribution of line means shown in (A). (C) Box plot of line means shown in (A). Distribution of female line means, with lines ordered by increasing mean. (E) Frequency distribution of line means shown in (D). (F) Box plot of line means shown in (D).

**Figure S9. Distributions of the sum of the numbers of bristles on the two most posterior abdominal sternites.** (A) Distribution of male line means, with lines ordered by increasing mean. (B) Frequency distribution of line means shown in (A). (C) Box plot of line means shown in (A). Distribution of female line means, with lines ordered by increasing mean. (E) Frequency distribution of line means shown in (D). (F) Box plot of line means shown in (D).

**Figure S10. Distributions of the difference of the numbers of bristles on the two most posterior abdominal sternites.** (A) Distribution of male line means, with lines ordered by increasing mean. (B) Frequency distribution of line means shown in (A). (C) Box plot of line means shown in (A). Distribution of female line means, with lines ordered by increasing mean. (E) Frequency distribution of line means shown in (D). (F) Box plot of line means shown in (D).

**Figure S11. Relationships between bristle trait means and micro-environmental variances (**In(m_e_)). (A) AB Sum: The sum of the numbers of bristles on the two most posterior abdominal sternites. (B) SB Sum: The sum of the numbers of bristles on the left and right sternopleural plates. (C) AB Diff: The difference between the numbers of bristles on the two most posterior abdominal sternites. F: females. M: Males.

**Figure S12. Receiver Operating Characteristics (ROC) curves from Random Forest discriminant models separating DGRP3 populations.** (A-C) ROC curves and area under the curve (AUC) values for models that compare each group against the rest of the lines using all 758,001 variants. (D-E) ROC curves and AUC values for models that compare each group against the rest of the lines using only the top 216 discriminator variants.

**Figure S13. Genomic distribution of polymorphic variants that discriminate among the three subpopulations in the DGRP3.** The Circos plot depicts the locations of genes, common large polymorphic inversions and the top 216 variants that discriminate among the three DGRP3 subpopulations from the random forest analysis on the *D. melanogaster* genome. The major chromosome arms and genomic positions are shown in the outer ring.

**Figure S14. Distribution of** LD **along the major chromosome arms.** The plots depict the mean *r*^2^ between variants 50-150 bp apart in 100 kb sliding windows in each 1 Mb region. (A) *X* chromosome. (B) Chromosome *2L*. (C) Chromosome *2R*. (D) Chromosome *3L*. (E) Chromosome *3R*.

**Figure S15. Histogram of numbers of high impact homozygous variants in DGRP3 lines.Orange bars: extant lines.** Green bars: extinct lines.

**Figure S16. GWA analysis of inbreeding depression.** (A) Manhattan plot of *—log*_10_(*P*) values. (B) Quantile-quantile plot of observed and expected —*log*_10_(*P*) values.

**Figure S17. GWA analyses of host control of microbiome composition**. The plots on the left are Manhattan plots of —*log*_10_(*P*) values, and on the right are quantile-quantile plots of observed and expected —*log*_10_ (*P*) values. (A) Simpson’s D^86^. (B) *Lactobacillus brevis*. (C) *Komagataeibacter hansenii*. (D) *Acetobacter pomorum*.

**Figure S18. GWA analyses of total sternopleural bristle (SB) numbers.** The plots on the left are Manhattan plots of —*log*_10_(*P*) values, and on the right are quantile-quantile plots of observed and expected —*log*_10_ (*P*) values. (A) Males. (B) Females. (C) Average of both sexes.

**Figure S19. GWA analyses of total abdominal bristle (AB) numbers.** The plots on the left are Manhattan plots of —*log*_10_(*P*) values, and on the right are quantile-quantile plots of observed and expected —*log*_10_ (*P*) values. (A) Males. (B) Females. (C) Average of both sexes. (D) Difference between females and males.

**Figure S20. GWA analyses of the difference in abdominal (AB) bristle number between the two most posterior sternites.** The plots on the left are Manhattan plots of —*log*_10_(*P*) values, and on the right are quantile-quantile plots of observed and expected —*log*_10_ (*P*) values. (A) Males. (B) Females. (C) Average of both sexes. (D) Difference between females and males.

**Figure S21. GWA analyses of** *lnσ_ε_* **total abdominal bristle (AB) number.** The plots on the left are Manhattan plots of —*log*_10_(*P*) values, and on the right are quantile-quantile plots of observed and expected —*log*_10_ (*P*) values. (A) Males. (B) Females.

**Figure S22. Statistical power to detect associations.** The power is calculated based on the 1-df χ^2^ test. For a standardized effect size of g, the explained heritability is 2p(1 — *p*)*β*^2^. The heritability of line means is assumed to be 1. Under these conditions, the test statistic under the alternative hypothesis is distributed as a non-central χ^z^ (*λ*) distribution with non-centrality parameter *λ* = *n* x 2*p*(1 — *p*)*β*^2^, where *n* is the number of lines. With a *P*-value threshold of *α* = 10^—5^, the power to detect the association is *P*(*χ*^2^(*λ*) > *χ*^2^). The power is plotted against effect size for different MAF when the number of lines is (A) 200 or (B) 1,200.

**Figure S23. Discovery of genotype-phenotype associations at increasing sample size.** The subsampling was performed by first randomly sampling 200 lines from the full dataset (*n* = 1,124) and incrementally adding more lines. GWA was performed on quantile-normalized female abdominal bristle number to mitigate effects of outliers. SNPs and genes were discovered at a nominal *P*-value threshold of 10^-5^. The MAF cutoff was 0.05 for 200 lines, 0.02 for 500 lines, and 0.01 for 1,000 and 1,124 lines.

## Supplementary Tables

**Table S1. DGRP3 lines**. (A) Line names, collection date (if known), and current survival status. (B) Number of sequenced reads, coverage (based on 143.7 Mb genome size) and SRA accession numbers.

**Table S2. Short nucleotide variant (SNV) annotation summary.** Summary statistics from the Variant Effect Predictor v111^125^ annotations for the short nucleotide variants. (A) Summary. (B) Coding variants. (C) Chromosomal distribution.

**Table S3. Estimates of nucleotide diversity (**n^61^**) per chromosome arm.** The estimates of π are averages of 10 kb windows. (A) *X* chromosome. (B) Chromosome *2L*. (C) Chromosome *2R*. (D) Chromosome *3L*. (E) Chromosome *3R*. (F) Chromosome *4*. Correlation refers to the correlation between x for SNPs and indels.

**Table S4. Structural variants (> 50 bp) in the DGRP3**. (A) Data for each insertion. POS is the genomic position at which the insertion occurred. (B) Distribution of segregating and homozygous insertions. (C) Data for each deletion. (D) Distribution of segregating and homozygous deletions. (E) Data for each tandem duplication. (F) Distribution of segregating and homozygous tandem duplications. (G) Data for each translocation. (H) Distribution of segregating and homozygous translocations. (I) Data for each inversion. (J) Distribution of segregating and homozygous inversions.

**Table S5. Common (minor allele frequency > 0.03) large (> 1 Mb) polymorphic inversions in the DGRP3.** (A) Data for each inversion. (B) DGRP2 polymorphic inversions. All inversions were mapped cytogenetically^45^. *Known molecular breakpoints called using v5 of the Drosophila genome^65^ were lifted over to the v6 version using the coordinates converter tool in FlyBase^10^. ^†^Approximate physical breakpoints based on cytogenetic breakpoints. (C) Large inversion karyotype status by DGRP3 line. 0: Homozygous reference (standard). 1: Segregating reference/alternate (standard, inversion). 2: Homozygous alternate (inversion).

**Table S6. High impact variants.** Variants were characterized using Variant Effect Predictor^125^. (A) Frameshift variants. (B) Splice acceptor variants. (C) Splice donor variants. (D) Start lost variants. (E) Stop gained variants. (F) Stop lost variants. (G) Transcript ablation variants. (H) Loss of Function genes by DGRP3 line. (I) Loss of Function genes by gene.

**Table S7. Estimates of Tajima’s** D**^68^.** (A-F) Estimates per chromosome arm, average of 10 kb windows. (A) *X* chromosome. (B) Chromosome *2L*. (C) Chromosome *2R*. (D) Chromosome *3L*. (E) Chromosome *3R*. (F) Chromosome *4*. Correlation refers to the correlation between *D* for SNPs and indels. (G) Estimates by gene, average of 1 kb windows. Genes in red font have values of D for SNPs < −2.000.

**Table S8. Mitochondrial variants and haplotypes.** (A) Mitochondrial SNV annotation summary. The summary statistics are from SNPEff v5.4 annotations. (B) Inference of mitochondrial haplotype clusters based on *Mclust* clustering algorithm. Uncertainty represents the complement of class posterior probability.

**Table S9. DGRP3 microbiome.** (A) Wolbachia infection status. (B) Relative abundances of microbial taxa for each DGRP3 line. Lines with 0 total microbial abundances have been removed. Relative abundances are multiplied by 100. (C) Core microbiome composition.

**Table S10. Bristle number data and quantitative genetic analyses.** (A) Raw data. (B) Line means. (C) ANOVAs. (D) Variance heterogeneity tests. (E-L) Analyses of natural selection on bristle number traits. Each tab shows the bristle phenotypes of extant lines and lines that died after their bristle numbers were quantified. The cumulative distributions indicate these phenotypes in purple (extant lines) and orange (extinct lines), and the frequency histograms show the same data for the entire population (full, turquoise) and currently extant (purple) lines. Differences between the means (*t-*tests) of extant and extinct lines indicate directional selection, and differences between the among-line variances (Levene’s tests) of extant and extinct lines indicate stabilizing selection when the variance among the extinct lines is greater than that of extant lines. *j* is a measure of the cumulative strength of stabilizing selection when the population phenotypic variance before selection (*V_P_*) is greater than the population variance after selection (*V_P_*\*)^52^. (E) Female AB sum. (F) Male AB sum. (G) Female *ln*(σ) AB sum. (H) Male *ln*(σ_ε_) AB sum. (I) Female SB sum. (J) Male SB sum. (K) Female *ln*(σ_ε_) SB sum. (L) Male *ln*(σ_ε_) SB sum.

**Table S11. Population structure.** (A) Top discriminating variants from Random Forest analysis separating the three subpopulations of DGRP3 lines identified from the first two Principal Components of all variants. (B) Genomic relatedness of closely related lines. (C-G) LD genome distribution. We computed the mean *r*^2^ between variants 50-150 bp apart in 100 kb sliding windows in each 1 Mb region for all major chromosome arms. (C) *X* chromosome. (D) Chromosome *2L*. (E) Chromosome *2R*. (F) Chromosome *3L*. (G) Chromosome *3L*.

**Table S12. GWA analysis of inbreeding depression.** (A) Top (*P* < 10^-6^) associations from a case-control GWA analysis of DGRP3 lines that died after sequencing and those that are extant. *P*-values < 10^-6^ are indicated in red font and *P*-values less than the conservative Bonferroni significance threshold (*P* < 3.33 x 10^-8^) are indicated in bold red font. (B) Unique variants and genes from the GWA analysis. (C) Gene Ontology enrichment analysis for associated genes.

**Table S13. Host control of microbiome composition.** (A) GWA analyses of variants and genes significantly (*P* < 10^-5^) associated with Simpson’s D^86^ alpha diversity and abundances of major species comprising the microbiome. *P*-values < 10^-6^ are indicated in red font and *P*-values less than the conservative Bonferroni significance threshold (*P* < 3.25 x 10^-8^) are indicated in bold red font. (B) Gene ontologies of significant genes. Genes with *P*-values < 10^-6^ are indicated in red font. Gene Ontology terms consistent with processes thought to be associated with host control of microbial composition are indicated in red font.

**Table S14. Sternopleural bristle GWA analyses.** (A) GWA analyses of variants and genes significantly (*P* < 10^-5^) associated with the sum of bristle numbers on the two sternopleural plates. *P*-values < 10^-6^ are indicated in red font and *P*-values less than the conservative Bonferroni significance threshold (*P* < 3.31 x 10^-8^) are indicated in bold red font. (B) Variants and genes. (C) Gene Ontologies of significant genes. Gene Ontology terms consistent with processes thought to be associated with sternopleural bristles are indicated in red font, and the most relevant terms are indicated in bold red font.

**Table S15. Abdominal bristle GWA analyses.** (A) GWA analyses of variants and genes significantly (*P* < 10^-5^) associated with the sum of bristle numbers on the two most posterior sternites, the difference in bristle numbers on the two most posterior sternites, and *ln*(*σ_ε_*) of the sum of bristles on the two most posterior sternites. *P*-values < 10^-6^ are indicated in red font and *P*-values less than the conservative Bonferroni significance threshold (*P* < 3.31 x 10^-8^) are indicated in bold red font. (B) Variants and genes. (C). Gene Ontology enrichment analyses. (D) Gene Ontologies of significant genes. Gene Ontology terms consistent with processes thought to be associated with abdominal bristles are indicated in red font, and the most relevant terms are indicated in bold red font.

**Table S16. Bristle genes.** Genes affecting SB and/or AB number from *P*-transposable element mutagenesis, complementation testing of naturally occurring alleles, and candidate gene association tests.

**Table S17. Abnormal Abdomen GWA analysis.** (A) GWA analysis of variants and genes significantly (*P* < 10^-5^) associated with the Abnormal Abdomen phenotype. *P*-values < 10^-6^ are indicated in red font and *P*-values less than the conservative Bonferroni significance threshold (*P* < 3.31 x 10^-8^) are indicated in bold red font. (B) Variants and genes.

## References

1. Adams, M. D., Celniker, S. E., Holt, R. A., Evans, C. A., Gocayne,J. D., Amanatides, P. G., Scherer, S. E., Li, P. W., Hoskins, R. A., Galle, R. F., et al. (2000). The genome sequence of *Drosophila melanogaster*. Science 287, 2185–2195. DOI: 10.1126/science.287.5461.2185.

2. International Human Genome Sequencing Consortium. (2001). Initial sequencing and analysis of the human genome. Nature 409, 860–921. DOI: 10.1038/35057062

3. Bellen, H. J., Levis, R. W., Liao, G., He, Y., Carlson, J. W., Tsang, G., Evans-Holm, M., Hiesinger, P. R., Schulze, K. L., Rubin, G. M., et al. (2004). The BDGP gene disruption project: single transposon insertions associated with 40% of Drosophila genes. Genetics 167, 761–781. DOI: 10.1534/genetics.104.026427

4. Bellen, H. J., Levis, R. W., He, Y., Carlson, J. W., Evans-Holm, M., Bae, E., Kim, J., Metaxakis, A., Savakis, C., Schulze, K. L., et al. (2011). The Drosophila gene disruption project: progress using transposons with distinctive site specificities. Genetics 188, 731–743. DOI: 10.1534/genetics.111.126995

5. Brand, A. H., and Perrimon, N. (1993). Targeted gene expression as a means of altering cell fates and generating dominant phenotypes. Development 118, 401–415. DOI: 10.1242/dev.118.2.401

6. Zeidler, M. P., Tan, C., Bellaiche, Y., Cherry, S., Häder, S., Gayko, U., and Perrimon, N. (2004). Temperature-sensitive control of protein activity by conditionally splicing inteins. Nat. Biotechnol. 22, 871–876. DOI: 10.1038/nbt979

7. Dietzl, G., Chen, D., Schnorrer, F., Su, K.-C., Barinova, Y., Fellner, M., Gasser, B., Kinsey, K., Oppel, S., Scheiblauer, S., et al. (2007). A genome-wide transgenic RNAi library for conditional gene inactivation in Drosophila. Nature 448, 151–156. DOI: 10.1038/nature05954

8. Hu, Y., Comjean, A., Rodiger, J., Liu, Y., Gao, Y., Chung, V., Zirin, J., Perrimon, N., and Mohr, S. E. (2021). FlyRNAi.org-the database of the Drosophila RNAi screening center and transgenic RNAi project: 2021 update. Nucleic Acids Res. 49, D908–D915. DOI: 10.1093/nar/gkaa936

9. Li, H., Janssens, J., De Waegeneer, M., Kolluru, S. S., Davie, K., Gardeux, V., Saelens, W., David, F. P. A., Brbić, M., Spanier, K., et al. (2022). Fly Cell Atlas: A single-nucleus transcriptomic atlas of the adult fruit fly. Science 375, eabk2432. DOI: 10.1126/science.abk2432

10. Öztürk-Çolak, A., Marygold, S. J., Antonazzo, G., Attrill, H., Goutte-Gattat, D., Jenkins, V. K., Matthews, B. B., Millburn, G., dos Santos, G., Tabone, C. J., et al. (2024). FlyBase: updates to the Drosophila genes and genomes database. Genetics 227, iyad211. DOI: 10.1093/genetics/iyad211

11. Clayton, G. A., Morris, J. A., and Robertson, A. (1957). An experimental check on quantitative genetical theory. I. Short-term responses to selection. J. Genet. 55, 131–151.

12. Clayton, G. A., and Robertson, A. (1957). An experimental check on quantitative genetical theory. II. The long-term effects of selection. J. Genet. 55, 152–170.

13. Frankham, R., Jones, L. P., and Barker, J. S. (1968). The effects of population size and selection intensity in selection for a quantitative character in Drosophila. I. Short-term response to selection. Genet. Res. 12, 237–248. DOI: 10.1017/s0016672300011848

14. Jones, L. P., Frankham, R., and Barker, J. S. (1968). The effects of population size and selection intensity in selection for a quantitative character in Drosophila. II. Long-term response to selection. Genet. Res. 12, 249–266. DOI: 10.1017/s001667230001185x

15. Yoo, B. H. (1980). Long-term selection for a quantitative character in large replicate populations of *Drosophila melanogaster*. I. Responses to selection. Genet. Res. 35, 1–17. DOI: 10.1007/BF00253881

16. Weber, K. E. (1990). Increased selection response in larger populations. I. Selection for wing-tip height in Drosophila melanogaster at three population sizes. Genetics 125, 579–584. DOI: 10.1093/genetics/125.3.579

17. Mukai, T., Chigusa, S. I., Mettler, L. E., and Crow, J. F. (1972). Mutation rate and dominance of genes affecting viability in *Drosophila melanogaster*. Genetics 72, 335–355. DOI: 10.1093/genetics/72.2.335

18. Houle, D., Hoffmaster, D. K., Assimacopoulos, S., and Charlesworth, B. (1992). The genomic mutation rate for fitness in Drosophila. Nature 359, 58–60. DOI: 10.1038/359058a0

19. Santiago, E., Albornoz, J., Domínguez, A., Toro, M. A., and López-Fanjul, C. (1992). The distribution of spontaneous mutations on quantitative traits and fitness in *Drosophila melanogaster*. Genetics 132, 771–781. DOI: 10.1093/genetics/132.3.771

20. Clayton, G. A., and Robertson, A. (1955). Mutation and quantitative variation. Am. Nat. 89, 151–158.

21. Caballero, A., Toro, M. A., and López-Fanjul, C. (1991). The response to artificial selection from new mutations in *Drosophila melanogaster*. Genetics 128, 89–102. DOI: 10.1093/genetics/128.1.89

22. Mackay, T. F. C., Lyman, R. F., Jackson, M. S., Terzian, C., and Hill, W. G. (1992). Polygenic mutation in *Drosophila melanogaster*: estimates from divergence among inbred strains. Evolution 46, 300–316. DOI: 10.1111/j.1558-5646.1992.tb02039.x

23. Mackay, T. F. C., Fry, J. D., Lyman, R. F., and Nuzhdin, S. V. (1994). Polygenic mutation in *Drosophila melanogaster*: estimates from response to selection of inbred strains. Genetics 136, 937–951. DOI: 10.1093/genetics/136.3.937

24. Houle, D., Hughes, K. A., Hoffmaster, D. K., Ihara, J., Assimacopoulos, S., Canada, D., and Charlesworth, B. (1994). The effects of spontaneous mutation on quantitative traits. I. Variances and covariances of life history traits. Genetics 138, 773–785. DOI: 10.1093/genetics/138.3.773

25. Clark, A. G., Wang, L., and Hulleberg, T. (1995). Spontaneous mutation rate of modifiers of metabolism in Drosophila. Genetics 139, 767–779. DOI: 10.1093/genetics/139.2.767

26. Aquadro, C. F., Desse, S. F., Bland, M. M., Langley, C. H., and Laurie-Ahlberg, C. C. (1986). Molecular population genetics of the *alcohol dehydrogenase* gene region of *Drosophila melanogaster*. Genetics 114, 1165–1190. DOI: 10.1093/genetics/114.4.1165

27. Langley, C. H., and Aquadro, C. F. (1987). Restriction-map variation in natural populations of *Drosophila melanogaster*: *white*-locus region. Mol. Biol. Evol. 4, 651–663. DOI: 10.1093/oxfordjournals.molbev.a040467

28. Schaeffer, S. W., Aquadro, C. F., and Langley, C. H. (1988). Restriction-map variation in the *Notch* region of *Drosophila melanogaster*. Mol. Biol. Evol. 5, 30–40. DOI: 10.1093/oxfordjournals.molbev.a040475

29. Miyashita, N., and Langley, C. H. (1988). Molecular and phenotypic variation of the *white* locus region in *Drosophila melanogaster*. Genetics 120, 199–212. DOI: 10.1093/genetics/120.1.199

30. Aguadé, M., Miyashita, N., and Langley, C. H. (1989). Reduced variation in the *yellow-achaete-scute* region in natural populations of *Drosophila melanogaster*. Genetics 122, 607–615. DOI: 10.1093/genetics/122.3.607

31. Begun, D. J., and Aquadro, C. F. (1992). Levels of naturally occurring DNA polymorphism correlate with recombination rates in *D. melanogaster*. Nature 356, 519–520. DOI: 10.1038/356519a0

32. Breese, E. L., and Mather, K. (1957). The organization of polygenic activity within a chromosome in Drosophila. I. Hair characters. Heredity 11, 373–395.

33. Thoday, J. M. (1961). Location of polygenes. Nature 191, 368–370.

34. Wolstenholme, D. R., and Thoday, J. M. (1963). Effects of disruptive selection. VII. A third chromosome polymorphism. Heredity 10, 413–431. DOI: 10.1038/hdy.1963.48

35. Shrimpton, A. E., and Robertson, A. (1988). The isolation of polygenic factors controlling bristle score in *Drosophila melanogaster*. I. Allocation of third chromosome sternopleural bristle effects to chromosome sections. Genetics 118, 437–443. DOI: 10.1093/genetics/118.3.437

36. Shrimpton, A. E., and Robertson, A. (1988). The isolation of polygenic factors controlling bristle score in *Drosophila melanogaster*. II. Distribution of third chromosome bristle effects within chromosome sections. Genetics 118, 445–459. DOI: 10.1093/genetics/118.3.445

37. Long, A. D., Mullaney, S. L., Reid, L. A., Fry, J. D., Langley, C. H., and Mackay, T. F. C. (1995). High resolution mapping of genetic factors affecting abdominal bristle number in *Drosophila melanogaster*. Genetics 139, 1273–1291. DOI: 10.1093/genetics/139.3.1273

38. Nuzhdin, S. V., Pasyukova, E. G., Dilda, C. L., Zeng, Z. B., and Mackay, T. F. C. (1997). Sex-specific quantitative trait loci affecting longevity in *Drosophila melanogaster*. Proc. Natl. Acad. Sci. U.S.A. 94, 9734–9739. DOI: 10.1073/pnas.94.18.9734

39. Mackay, T. F. C., and Langley, C. H. (1990). Molecular and phenotypic variation in the *achaete-scute* region of *Drosophila melanogaster*. Nature 348, 64–66. DOI: 10.1038/348064a0

40. Lai, C., Lyman, R. F., Long, A. D., Langley, C. H., and Mackay, T. F. C. (1994). Naturally occurring variation in bristle number and DNA polymorphisms at the *scabrous* locus of *Drosophila melanogaster*. Science 266, 1697–1702. DOI: 10.1126/science.7992053

41. Long, A. D., Lyman, R. F., Langley, C. H., and Mackay, T. F. C. (1998). Two sites in the *Delta* gene region contribute to naturally occurring variation in bristle number in *Drosophila melanogaster*. Genetics 149, 999–1017. DOI: 10.1093/genetics/149.2.999

42. De Luca, M., Roshina, N. V., Geiger-Thornsberry, G. L., Lyman, R. F., Pasyukova, E. G., and Mackay, T. F. C. (2003). *Dopa decarboxylase* (*Ddc*) affects variation in Drosophila longevity. Nat. Genet. 34, 429–433. DOI: 10.1038/ng1218

43. Carbone, M. A., Jordan, K. W., Lyman, R. F., Harbison, S. T., Leips, J., Morgan, T. J., DeLuca, M., Awadalla, P., and Mackay, T. F. C. (2006). Phenotypic variation and natural selection at *Catsup*, a pleiotropic quantitative trait gene in Drosophila. Curr. Biol. 16, 912–919. DOI: 10.1016/j.cub.2006.03.051

44. Mackay, T. F. C., Richards, S., Stone, E. A., Barbadilla, A., Ayroles, J. F., Zhu, D., Casillas, S., Han, Y., Magwire, M. M., Cridland, J. M., et al. (2012). The *Drosophila melanogaster* Genetic Reference Panel. Nature 482, 173–178. DOI: 10.1038/nature10811

45. Huang, W., Massouras, A., Inoue, Y., Peiffer, J., Ràmia, M., Tarone, A. M., Turlapati, L., Zichner, T., Zhu, D., Lyman, R. F., et al. (2014). Natural variation in genome architecture among 205 *Drosophila melanogaster* Genetic Reference Panel lines. Genome Res. 24, 1193–1208. DOI: 10.1101/gr.171546.113

46. Sieberts, S. K., and Schadt, E. E. (2007). Moving toward a system genetics view of disease. Mamm. Genome 18, 389–401. DOI: 10.1007/s00335-007-9040-6

47. Civelek, M., and Lusis, A. J. (2014). Systems genetics approaches to understand complex traits. Nat. Rev. Genet. 15, 34–48. DOI: 10.1038/nrg3575

48. Everett, L. J., Huang, W., Zhou, S., Carbone, M. A., Lyman, R. F., Arya, G. H., Geisz, M. S., Ma, J., Morgante, F., St Armour G, et al. (2020). Gene expression networks in the Drosophila Genetic Reference Panel. Genome Res. 30, 485–496. DOI: 10.1101/gr.257592.119

49. Zhou, S., Morgante, F., Geisz, M. S., Ma, J., Anholt, R. R. H., and Mackay, T. F. C. (2020). Systems genetics of the Drosophila metabolome. Genome Res 30, 392–405. DOI: 10.1101/gr.243030.118

50. Huang, W., Carbone, M. A., Lyman, R. F., Anholt, R. R. H., and Mackay, T. F. C. (2020). Genotype by environment interaction for gene expression in *Drosophila melanogaster*. Nat. Commun 11, 5451. DOI: 10.1038/s41467-020-19131-y

51. Huang, W., Campbell, T., Carbone, M. A., Jones, W. E., Unselt, D., Anholt, R. R. H., and Mackay, T. F. C. (2020). Context-dependent genetic architecture of Drosophila life span. PLoS Biol. 18, e3000645. DOI: 10.1371/journal.pbio.3000645

52. Falconer, D. S., and Mackay, T. F. C. (1996). Introduction to Quantitative Genetics. Longman Group Limited, Burnt Hill, Harlow, Essex.

53. Cockerham, C. C. (1954). An extension of the concept of partitioning hereditary variance for analyses of covariance among relatives when epistasis is present. Genetics 39, 859–882. DOI: 10.1093/genetics/39.6.859

54. Huang, W., Richards, S., Carbone, M. A., Zhu, D., Anholt, R. R. H., Ayroles, J. F., Duncan, L., Jordan, K. W., Lawrence, F., Magwire, M. M., et al. (2012). Epistasis dominates the genetic architecture of Drosophila quantitative traits. Proc. Natl. Acad. Sci. U.S.A. 109, 15553–15559. DOI: 10.1073/pnas.1213423109

55. He, X., Zhou, S., St Armour, G. E., Mackay, T. F..C, and Anholt, R. R. H. (2016). Epistatic partners of neurogenic genes modulate Drosophila olfactory behavior. Genes Brain Behav. 15, 280–290. DOI: 10.1111/gbb.12279

56. He, B. Z., Ludwig, M. Z., Dickerson, D. A., Barse, L., Arun, B., Vilhjálmsson, B. J., Jiang, P., Park, S. Y., Tamarina, N. A., Selleck, S. B., et al. (2014). Effect of genetic variation in a Drosophila model of diabetes-associated misfolded human proinsulin. Genetics 196, 557–567. DOI: 10.1534/genetics.113.157800

57. Mackay, T. F. C., and Huang, W. (2018). Charting the genotype-phenotype map: lessons from the *Drosophila melanogaster* Genetic Reference Panel. Wiley Interdiscip. Rev. Dev. Biol. 7, 10.1002/wdev.289. DOI: 10.1002/wdev.289

58. Lavoy, S., Chittoor-Vinod, V. G., Chow, C. Y., and Martin, I. (2018). Genetic modifiers of neurodegeneration in a Drosophila model of Parkinson’s Disease. Genetics 209, 1345–1356. DOI: 10.1534/genetics.118.301119

59. Talsness, D. M., Owings, K. G., Coelho, E., Mercenne, G., Pleinis, J. M., Partha, R., Hope, K. A., Zuberi, A. R., Clark, N. L., Lutz, C. M., et al. (2020). A Drosophila screen identifies NKCC1 as a modifier of NGLY1 deficiency. Elife 9, e57831. DOI: 10.7554/eLife.57831

60. Campos, J. L., Halligan, D. L., Haddrill, P. R., and Charlesworth, B. (2014). The relation between recombination rate and patterns of molecular evolution and variation in *Drosophila melanogaster*. Mol. Biol. Evol. 31, 1010–1028. DOI: 10.1093/molbev/msu056

61. Nei, M., and Li, W.-H. (1979). Mathematical model for studying genetic variation in terms of restriction endonucleases. Proc. Natl. Acad. Sci. U.S.A. 76, 5269–5273. DOI: 10.1073/pnas.76.10.5269

62. Bridges, C. B. (1935). Salivary chromosome maps with a key to the banding of the chromosomes of *Drosophila melanogaster*. J. Hered. 26, 60–64.

63. Sorsa, V. (1988). Chromosome Maps of Drosophila, Volume II. CRC Press Inc, Boca Raton, Florida.

64. Lemeunier, F., David, J. R., Tsacas, L., and Ashburner, M. (1986). The melanogaster species group. Pp 147–256 in The Genetics and Biology of Drosophila Volume 3c, edited by Ashburner M, Carson HL, Thompson Jr JN. Academic Press Inc, London.

65. Corbett-Detig, R. B., and Hartl, D. L. (2012). Population genomics of inversion polymorphisms in *Drosophila melanogaster*. PLoS Genet. 8, e1003056. DOI: 10.1371/journal.pgen.1003056

66. Charlesworth, B., Morgan, M. T., and Charlesworth, D. (1993). The effect of deleterious mutations on neutral molecular variation. Genetics 134, 1289–1303. DOI: 10.1093/genetics/134.4.1289

67. Kryukov, G. V., Pennacchio, L. A., and Sunyaev, S. R. (2007). Most rare missense alleles are deleterious in humans: implications for complex disease and association studies. Am. J. Hum. Genet. 80, 727–739. DOI: 10.1086/513473

68. Tajima, F. (1989). Statistical method for testing the neutral mutation hypothesis by DNA polymorphism. Genetics 123, 585–595. DOI: 10.1093/genetics/123.3.585

69. David, J. R., and Capy, P. (1988). Genetic variation of *Drosophila melanogaster* natural populations. Trends Genet. 4, 106–111. DOI: 10.1016/0168-9525(88)90098-4

70. Keller, A. (2007). *Drosophila melanogaster*’s history as a human commensal. Curr. Biol. 17, R77–R81. DOI: 10.1016/j.cub.2006.12.031

71. Campo, D., Lehmann, K., Fjeldsted, C., Souaiaia, T., Kao, J., and Nuzhdin, S. V. (2013). Whole-genome sequencing of two North American *Drosophila melanogaster* populations reveals genetic differentiation and positive selection. Mol. Ecol. 22, 5084–5097. DOI: 10.1111/mec.12468

72. Leigh, J. W., and Bryant, D. (2015). POPART: full-feature software for haplotype network construction. Methods Ecol. Evol. 6, 1110–1116. DOI10.1111/2041-210X.12410

73. Scrucca, L., Fop, M., Murphy, T. B., and Raftery, A. E. (2016). Mclust5: Clustering, classification, and density estimation using Gaussian finite mixture models. R. J. 8, 289–317.

74. Shin, S. C., Kim, S.-H., You, H., Kim, B., Kim, A. C., Lee, K.-A., Yoon, J.-H., Ryu, J.-H., and Lee, W.-J. (2011). Drosophila microbiome modulates host developmental and metabolic homeostasis via insulin signaling. Science 334, 670–674. DOI: 10.1126/science.1212782

75. Keebaugh, E. S., Yamada, R., Obadia, B., Ludington, W. B., and Ja, W. W. (2018). Microbial quantity impacts Drosophila nutrition, development, and lifespan. iScience 4, 247–259. DOI: 10.1016/j.isci.2018.06.004

76. Kamareddine, L., Robins, W. P., Berkey, C. D., Mekalanos, J. J., and Watnick, P. I. (2018). The Drosophila immune deficiency pathway modulates enteroendocrine function and host metabolism. Cell Metab. 28, 449–462. DOI: 10.1016/j.cmet.2018.05.026

77. Cho, K. H., and Kang, S. O. (2025). The gut microbiota of *Drosophila melanogaster*: A model for host–microbe interactions in metabolism, immunity, behavior, and disease. Microorganisms 13, 2515. DOI: 10.3390/microorganisms13112515

78. Storelli, G., Defaye, A., Erkosar, B., Hols, P., Royet, J., and Leulier, F. (2011). *Lactobacillus plantarum* promotes Drosophila systemic growth by modulating hormonal signals through TOR-dependent nutrient sensing. Cell Metab. 14, 403–414.

79. Storelli, G., Strigini, M., Grenier, T., Bozonnet, L., Schwarzer, M., Daniel, C., Matos, R., and Leulier, F. (2018). Drosophila perpetuates nutritional mutualism by promoting the fitness of its intestinal symbiont *Lactobacillus plantarum*. Cell Metab. 27, 362–377. DOI: 10.1016/j.cmet.2011.07.012

80. Consuegra, J., Grenier, T., Akherraz, H., Rahioui, I., Gervais, H., da Silva, P., and Leulier, F. (2020). Metabolic cooperation among commensal bacteria supports Drosophila juvenile growth under nutritional stress. iScience 23, 101232. DOI: 10.1016/j.isci.2020.101232

81. Newell, P. D., and Douglas, A. E. (2014). Interspecies interactions determine the impact of the gut microbiota on nutrient allocation in *Drosophila melanogaster*. Appl. Environ. Microbiol. 80, 788–796. DOI: 10.1128/AEM.02742-13

82. Téfit, M. A., and Leulier, F. (2017). *Lactobacillus plantarum* favors the early emergence of fit and fertile adult Drosophila upon chronic undernutrition. J. Exp. Biol. 220, 900–907. DOI: 10.1242/jeb.151522

83. Beribaka, M., Jelić, M., Tanasković, M., Lazić, C., and Stamenković-Radak, M. (2021). Life history traits in two Drosophila species differently affected by microbiota diversity under lead exposure. Insects 12, 1122. DOI: 10.3390/insects12121122

84. Suito, T., Nagao, K., Juni, N., Hara, Y., Sokabe, T., Atomi, H., and Umeda, M. (2022). Regulation of thermoregulatory behavior by commensal bacteria in Drosophila. Biosci. Biotechnol. Biochem. 86, 1060–1070. DOI: 10.1093/bbb/zbac087

85. Shannon, C. E. (1948). A mathematical theory of communication. Bell System Technical Journal 27, 379–423, 623–656. DOI.org/10.1002/j.1538-7305.1948.tb01338.x

86. Simpson, E. H. (1949). Measurement of diversity. Nature 163, 688. DOI10.1038/163688a0

87. Hill, M. O. (1973). Diversity and evenness: A unifying notation and its consequences. Ecology 54, 427–432. DOI10.2307/1934352

88. Bray, J. R., and Curtis, J. T. (1957). An ordination of the upland forest communities of southern Wisconsin. Ecological Monographs 27, 325–349. DOI10.2307/1942268

89. Henriques, S. F., Dhakan, D. B., Serra, L., Francisco, A. P., Carvalho-Santos, Z., Baltazar, C., Elias, A. P., Anjos, M., Zhang, T., Maddocks, O. D. K., et al. (2020). Metabolic cross-feeding in imbalanced diets allows gut microbes to improve reproduction and alter host behaviour. Nat. Commun. 11, 4236. DOI: 10.1038/s41467-020-18049-9

90. Rowe, S. J., White, I. M. S., Avendaño, S., and Hill, W. G. (2006). Genetic heterogeneity of residual variance in broiler chickens. Genet. Sel. Evol. 38, 617–635. DOI: 10.1186/1297-9686-38-6-617

91. Ordas, B., Malvar, R. A., and Hill, W. G. (2008). Genetic variation and quantitative trait loci associated with developmental stability and the environmental correlation between traits in maize. Genet. Res. 90, 385–395. DOI: 10.1017/S0016672308009762

92. Morgante, F., Sørensen, P., Sorensen, D. A., Maltecca, C., and Mackay, T. F. C. (2015). Genetic architecture of micro-environmental plasticity in *Drosophila melanogaster*. Sci. Rep. 5, 9785. DOI: 10.1038/srep09785

93. Sobels, S. H. (1952). Genetics and morphology of the genotype *Asymmetric* with special reference to its *Abnormal Abdomen* character (*Drosophila melanogaster*). Genetica 26, 117–279.

94. Mackay, T. F. C., Lyman, R. F., and Hill, W. G. (1995). Polygenic mutation in *Drosophila melanogaster*: non-linear divergence among unselected strains. Genetics 139, 849–859. DOI: 10.1093/genetics/139.2.849

95. Mackay, T. F. C., and Lyman, R. F. (2005). Drosophila bristles and the nature of quantitative genetic variation. Philos. Trans. R. Soc. Lond. B Biol. Sci. 360, 1513–1527. DOI: 10.1098/rstb.2005.1672

96. VanRaden, P. M. (2008). Efficient methods to compute genomic predictions. J. Dairy Sci. 91, 4414–4423. DOI: 10.3168/jds.2007-0980

97. Ober, U., Ayroles, J. F., Stone, E. A., Richards, S., Zhu, D., Gibbs, R. A., Stricker, C., Gianola, D., Schlather, M., Mackay, T. F. C., et al. (2012). Using whole-genome sequence data to predict quantitative trait phenotypes in *Drosophila melanogaster*. PLoS Genet 8, e1002685. DOI: 10.1371/journal.pgen.1002685

98. Hill, W. G., and Robertson, A. (1966). The effect of linkage on limits to artificial selection. Genet. Res. 8, 269–294. DOI: 10.1017/S001667230800949X

99. Naitza, S., and Ligoxygakis, P. (2004). Antimicrobial defences in Drosophila: the story so far. Mol. Immunol. 40, 887–896. DOI: 10.1016/j.molimm.2003.10.008

100. Kaneko, T., and Silverman, N. (2005). Bacterial recognition and signalling by the Drosophila IMD pathway. Cell Microbiol. 7, 461–469. DOI: 10.1111/j.1462-5822.2005.00504.x

101. Mallick, S., and Eleftherianos, I. (2024). Role of cuticular genes in the insect antimicrobial immune response. Front. Cell Infect. Microbiol. 14, 1456075. DOI: 10.3389/fcimb.2024.1456075

102. zur Lage, P., Shrimpton, A. D., Flavell, A. J., Mackay, T. F. C., and Brown, A. J. (1997). Genetic and molecular analysis of *smooth*, a quantitative trait locus affecting bristle number in *Drosophila melanogaster*. Genetics 146, 607–618. DOI: 10.1093/genetics/146.2.607

103. Lai, C., McMahon, R., Young, C., Mackay, T. F. C., and Langley, C. H. (1998). *quemao*, a Drosophila bristle locus, encodes geranylgeranyl pyrophosphate synthase. Genetics 149, 1051–1061. DOI: 10.1093/genetics/149.2.1051

104. Norga, K. K., Gurganus, M. C., Dilda, C. L., Yamamoto, A., Lyman, R. F., Patel, P. H., Rubin, G. M., Hoskins, R. A., Mackay, T. F. C., and Bellen, H. J. (2003). Quantitative analysis of bristle number in Drosophila mutants identifies genes involved in neural development. Curr. Biol. 13, 1388–1396. DOI: 10.1016/s0960-9822(03)00546-3

105. Long, A. D., Mullaney, S. L., Mackay, T. F. C., and Langley, C. H. (1996). Genetic interactions between naturally occurring alleles at quantitative trait loci and mutant alleles at candidate loci affecting bristle number in *Drosophila melanogaster*. Genetics 144, 1497–1510. DOI: 10.1093/genetics/144.4.1497

106. Lyman, R. F., and Mackay, T. F. C. (1998). Candidate quantitative trait loci and naturally occurring phenotypic variation for bristle number in *Drosophila melanogaster*: the *Delta-Hairless* gene region. Genetics 149, 983–998. DOI: 10.1093/genetics/149.2.983

107. Long, A. D., Lyman, R. F., Morgan, A. H., Langley, C. H., and Mackay, T. F. C. (2000). Both naturally occurring insertions of transposable elements and intermediate frequency polymorphisms at the *achaete-scute* complex are associated with variation in bristle number in *Drosophila melanogaster*. Genetics 154, 1255–1269. DOI: 10.1093/genetics/154.3.1255

108. Robin, C., Lyman, R. F., Long, A. D., Langley, C. H., and Mackay, T. F. C. (2002). *hairy*: A quantitative trait locus for Drosophila sensory bristle number. Genetics 162, 155–164. DOI: 10.1093/genetics/162.1.155

109. modENCODE Consortium, Roy, S., Ernst, J., Kharchenko, P. V., Kheradpour, P., Negre, N., Eaton, M. L., Landolin, J. M., Bristow, C. A., Ma, L., et a. (2010). Identification of functional elements and regulatory circuits by Drosophila modENCODE. Science 330, 1787–1797. DOI 10.1126/science.1198374.

110. Bryant, C. D., Smith, D. J., Kantak, K. M., Nowak, T. S. Jr., Williams, R. W., Damaj, M. I., Redei, E. E., Chen, H., and Mulligan, M. K. (2020). Facilitating complex trait analysis via reduced complexity crosses. Trends Genet. 36, 549–562. DOI: 10.1016/j.tig.2020.05.003

111. Mackay, T. F. C., Lyman, R. F., and Jackson, M. S. (1992). Effects of *P* element insertions on quantitative traits in *Drosophila melanogaster*. Genetics 130, 315–332. DOI: 10.1093/genetics/130.2.315

112. Lyman, R. F., Lawrence, F., Nuzhdin, S. V., and Mackay, T. F. C. (1996). Effects of single *P*-element insertions on bristle number and viability in *Drosophila melanogaster*. Genetics 143, 277–292. DOI: 10.1093/genetics/143.1.277

113. Kearsey, M. J., and Barnes, B. W. (1970). Variation for metrical characters in Drosophila populations. II. Natural selection. Heredity 25, 11–21. DOI: 10.1038/hdy.1970.2

114. Robertson, A. (1955). Selection in animals: synthesis. Cold Spring Harbor Symp. Quant. Biol. 20, 225–229. DOI: 10.1101/sqb.1955.020.01.021

115. Robertson, A. (1967). The nature of quantitative variation. pp. 265-280 in Heritage From Mendel, edited by A. Brink. The University of Wisconsin Press, Madison, WI.

116. Spiers, J. G. C. (1974). The effects of larval competition on a quantitative character in *Drosophila melanogaster*. Ph.D. Thesis, University of Edinburgh, Edinburgh, Scotland.

117. Nadeau, J. H., Singer, J. B., Matin, A., and Lander, E. S. (2000). Analysing complex genetic traits with chromosome substitution strains. Nat. Genet. 24, 221–225. DOI: 10.1038/73427

118. Singer, J. B., Hill, A. E., Burrage, L. C., Olszens, K. R., Song, J., Justice, M., O’Brien, W. E., Conti, D. V., Witte, J. S., Lander, E. S., et al. (2004). Genetic dissection of complex traits with chromosome substitution strains of mice. Science 304, 445–448. DOI: 10.1126/science.1093139

119. Chen, S., Zhou, Y., Chen, Y., and Gu, J. (2018). fastp: an ultra-fast all-in-one FASTQ preprocessor. Bioinformatics 34, i884–i890. DOI: 10.1093/bioinformatics/bty560

120. Li, H., and Durbin, R. (2009). Fast and accurate short read alignment with Burrows-Wheeler transform. Bioinformatics 25, 1754–1760. DOI: 10.1093/bioinformatics/btp324

121. Li, H., Handsaker, B., Wysoker, A., Fennell, T., Ruan, J., Homer, N., Marth, G., Abecasis, G., and Durbin, R. (2009). 1000 Genome Project Data Processing Subgroup. The Sequence Alignment/Map format and SAMtools. Bioinformatics 25, 2078–2079. DOI: 10.1093/bioinformatics/btp352

122. DePristo, M. A., Banks, E., Poplin, R., Garimella, K. V., Maguire, J. R., Hartl, C., Philippakis, A. A., del Angel, G., Rivas, M. A., Hanna, M., et al. (2011). A framework for variation discovery and genotyping using next-generation DNA sequencing data. Nat. Genet. 43, 491–498. DOI: 10.1038/ng.806

123. Stone, E. A. (2012). Joint genotyping on the fly: identifying variation among a sequenced panel of inbred lines. Genome Res. 22, 966–974. DOI: 10.1101/gr.129122.111

124. Purcell, S., Neale, B., Todd-Brown, K., Thomas, L., Ferreira, M. A., Bender, D., Maller, J., Sklar, P., De Bakker, P. I., Daly, M. J., et al. (2007). PLINK: a tool set for whole-genome association and population-based linkage analyses. Am. J. Hum. Genet. 81, 559–575. DOI: 10.1086/519795

125. McLaren, W., Gil, L., Hunt, S. E., Riat, H. S., Ritchie, G. R. S., Thormann, A., Flicek, P., and Cunningham, F. (2016). The Ensembl Variant Effect Predictor. Genome Biol. 17, 122. DOI: 10.1186/s13059-016-0974-4

126. Chen, X., Schulz-Trieglaff, O., Shaw, R., Barnes, B., Schlesinger, F., Källberg, M., Cox, A. J., Kruglyak, S., and Saunders, C. T. (2016). Manta: rapid detection of structural variants and indels for germline and cancer sequencing applications. Bioinformatics 32, 1220–1222. DOI: 10.1093/bioinformatics/btv710

127. Jeffares, D. C., Jolly, C., Hoti, M., Speed, D., Shaw, L., Rallis, C., Balloux, F., Dessimoz, C., Bähler, J., and Sedlazeck, F. J. (2017). Transient structural variations have strong effects on quantitative traits and reproductive isolation in fission yeast. Nat. Commun. 8, 14061. DOI: 10.1038/ncomms14061

128. Gu, Z., Gu, L., Eils, R., Schlesner, M., and Brors B. 2014. Circlize implements and enhances circular visualization in R. Bioinformatics 30, 2811–2812. DOI: 10.1093/bioinformatics/btu393

129. Danecek, P., Bonfield, J. K., Liddle, J., Marshall, J., Ohan, V., Pollard, M. O., Whitwham, A., Keane, T., McCarthy, S. A., Davies, R. M., et al. (2021). Twelve years of SAMtools and BCFtools. Gigascience 10, giab008. DOI: 10.1093/gigascience/giab008

130. Katoh, K., Misawa, K., Kuma, K., and Miyata, T. (2002) MAFFT: a novel method for rapid multiple sequence alignment based on fast Fourier transform. Nucleic Acids Res. 30, 3059–3066. DOI: 10.1093/nar/gkf436

131. Liang, G., Mi, D., Chang, J., Yau, T. O., Xu, G., Ruan, J., Bu, W., and Gao, S. (2022). Precise annotation of Drosophila mitochondrial genomes leads to insights into AT-rich regions. Mitochondrion 65, 145–149. DOI: 10.1016/j.mito.2022.06.006

132. Bushnell, B., Rood, J., and Singer, E. (2017). BBMerge - Accurate paired shotgun read merging via overlap. PLoS One 12, e0185056. DOI: 10.1371/journal.pone.0185056

133. Visconti, A., Martin, T. C., and Falchi, M. (2018). YAMP: a containerized workflow enabling reproducibility in metagenomics research. Gigascience 7, giy072. DOI: 10.1093/gigascience/giy072

134. Truong, D. T., Franzosa, E. A., Tickle, T. L., Scholz, M., Weingart, G., Pasolli, E., Tett, A., Huttenhower, C., and Segata, N. (2015). MetaPhlAn2 for enhanced metagenomic taxonomic profiling. Nat. Methods 12, 902–903. DOI: 10.1038/nmeth.3589

135. McKinney, W. (2010). Data structures for statistical computing in Python. SciPy Proceedings 2010, 51–56. DOI10.25080/Majora-92bf1922-00a

136. Kruchten, N., Seier, A., and Parmer, C. (2026). An interactive, open-source, and browser-based graphing library for Python. Version 6.6.0. DOI:10.5281/zenodo.14503524.

137. Benjamini, Y., and Hochberg, Y. (1995). Controlling the false discovery rate: a practical and powerful approach to multiple testing. J. Roy. Stat. Soc. B (Methodological) 57, 289–300. DOI10.1111/j.2517-6161.1995.tb02031.x

138. Shannon, P., Markiel, A., Ozier, O., Baliga, N. S., Wang, J. T., Ramage, D., Amin, A., Schwikowski, B., and Ideker, T. (2003). Cytoscape: A software environment for integrated models of biomolecular interaction networks. Genome Res. 13, 2498–2504. DOI: 10.1101/gr.1239303

139. Virtanen, P., Gommers, R., Oliphant, T. E., Haberland, M., Reddy, T., Cournapeau, D., Burovski, E., Peterson, P., Weckesser, W., Bright, J., et al. (2020). SciPy 1.0: fundamental algorithms for scientific computing in Python. Nat. Methods 17, 261–272. DOI: 10.1038/s41592-019-0686-2

140. Aton, M., McDonald, D., Cañardo Alastuey, J., Azom, R., Batra, P., Bezshapkin, V., Bolyen, E., Cagle, A., Caporaso, J. G., Debelius, J. W., et al. (2026). Scikit-bio: a fundamental Python library for biological omic data analysis. Nat. Methods 23, 274–276. DOI: 10.1038/s41592-025-02981-z

141. Kramer, O. (2016). Scikit-Learn, pp 45-53 in Machine Learning for Evolution Strategies. Springer International Publishing, Switzerland.

142. Waskom, M. L. (2021). seaborn: statistical data visualization. J. Open Source Software 6, 3021. DOI10.21105/joss.03021

143. Zheng, X., Levine, D., Shen, J., Gogarten, S. M., Laurie, C., and Weir, B. S. (2012). A high-performance computing toolset for relatedness and principal component analysis of SNP data. Bioinformatics 28, 3326–3328. DOI: 10.1093/bioinformatics/bts606

144. Liaw, A., and Wiener, M. 2002. Classification and regression by randomForest. R News 2/3, 18–22.

145. Endelman, J. B. (2011). Ridge regression and other kernels for genomic selection with R Package rrBLUP. The Plant Genome 4, 250–255. DOI: 10.3835/plantgenome2011.08.0024

146. Sturn, A., Quackenbush, J., and Trajanoski, Z. (2022). Genesis: Cluster analysis of microarray data. Bioinformatics 18, 207–208. DOI: 10.1093/bioinformatics/18.1.207

147. SAS Studio® Statistical Analysis Software Version 3.8. 2018. SAS® Institute Inc., Cary, NC.

148. Zhou, X., and Stephens, M. (2012). Genome-wide efficient mixed-model analysis for association studies. Nat. Genet. 44, 821–824. DOI: 10.1038/ng.2310

149. Turner, S. D. (2018). qqman: an R package for visualizing GWAS results using QQ and Manhattan plots. Journal of Open Source Software 3, 731. DOI: 10.21105/joss.00731

150. Aitchison, J. (1986). The Statistical Analysis of Compositional Data. London: Chapman and Hall.

151. Gloor, G. B., Macklaim, J. M., Pawlowsky-Glahn, V., and Egozcue, J. J. (2017). Microbiome datasets are compositional: and this is not optional. Front. Microbiol. 8, 2224. DOI10.3389/fmicb.2017.02224

