## Supplementary Figures for "The *Drosophila melanogaster* Genetic Reference Panel, Version 3 (DGRP3)"

**Figure S1. Minor allele frequency (MAF) distributions.** MAF frequency distributions are shown for all polymorphic molecular variant categories. Small insertions and deletions are ≤ 50 bp, large structural variants are > 50 bp. (A) SNPs. (B) Small insertions. (C) Small deletions. (D) Large insertions. (E) Large deletions. (F) Large tandem duplications. (G) Large inversions. (H) Large interchromosomal translocations. Inserts magnify the right tails (common variants) of the distributions.

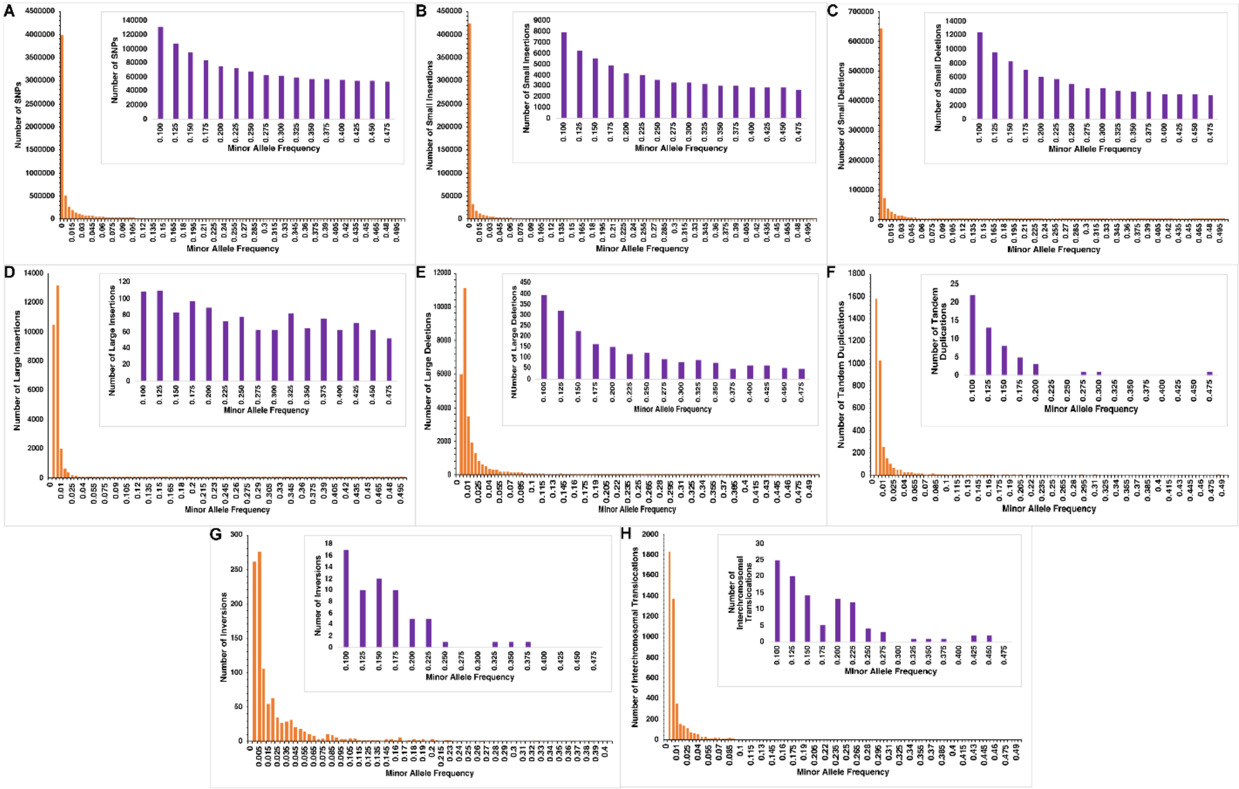

**Figure S2. Distributions of Loss of Function (LoF) genes.** (A) Frequency histogram of the number of LoF genes per DGRP3 line. (B) Frequency histogram of the number of DGRP3 lines harboring LoF genes.

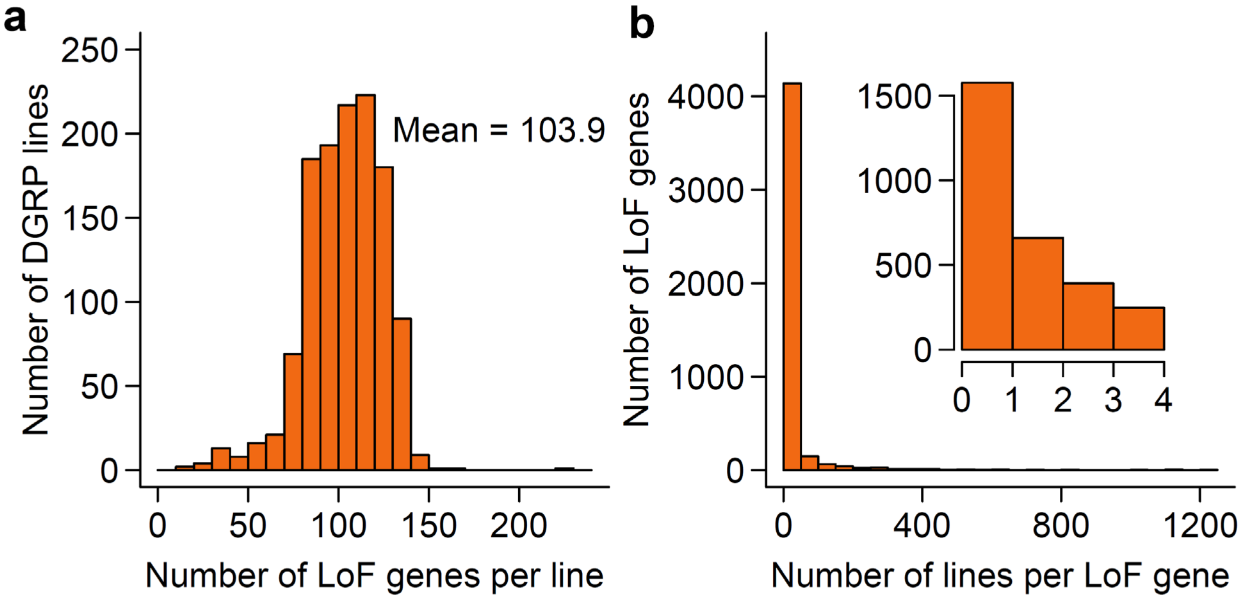

**Figure S3. Mitochondrial haplotypes.** The 16 major mitochondrial haplotypes (black circles) are depicted, as well as the DGRP3 line composition of each haplotype. Orange circles represent the haplotypes and the size of the circles represents the number of lines within each of the haplogroups. The orthogonal notches on the edges represent the number of variants separating the nodes (variant distance). Ellipses encompass large clusters of haplogroups and singlet haplotypes within one edge.

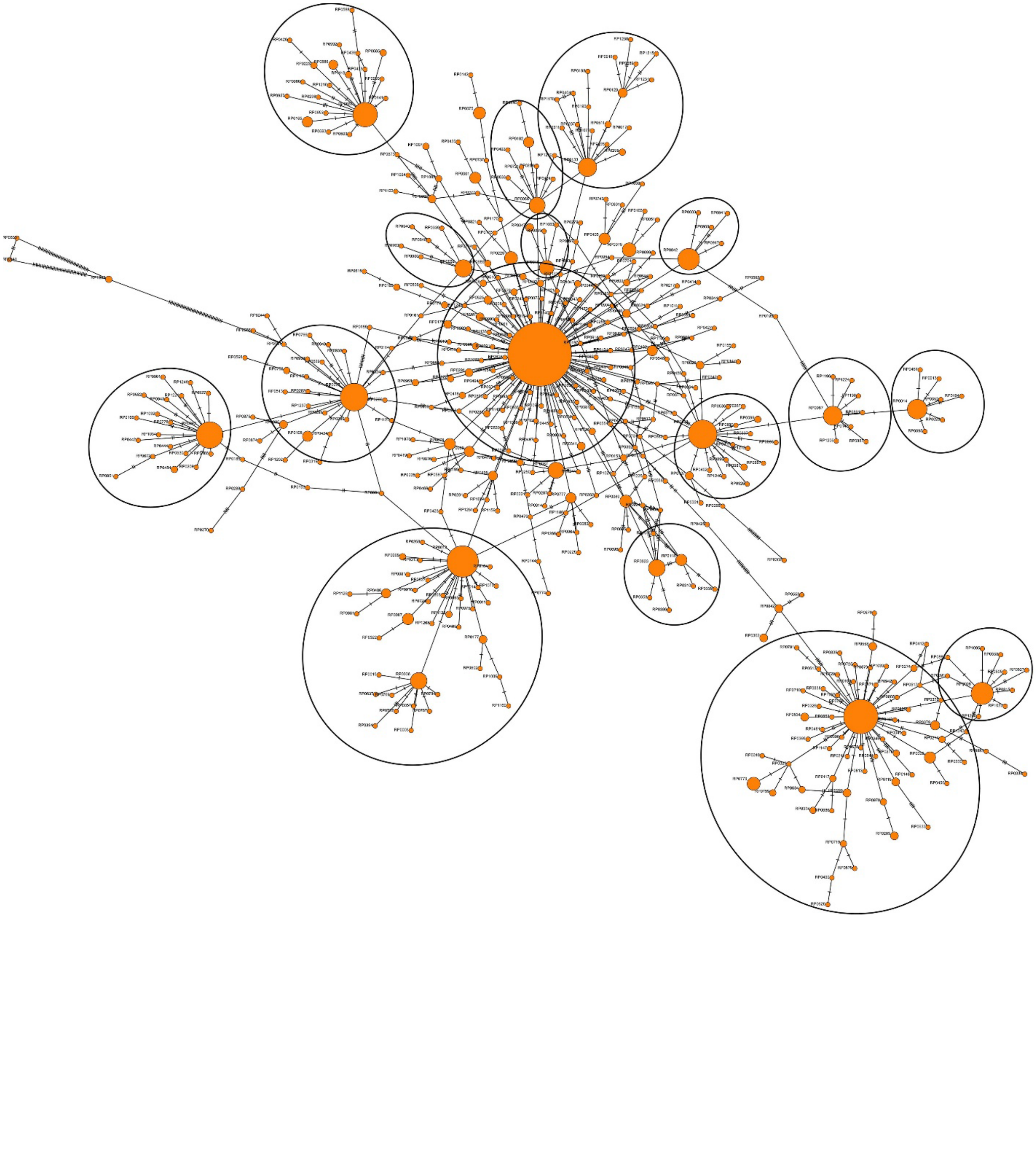

**Figure S4. Composition of the core DGRP3 microbiome.**

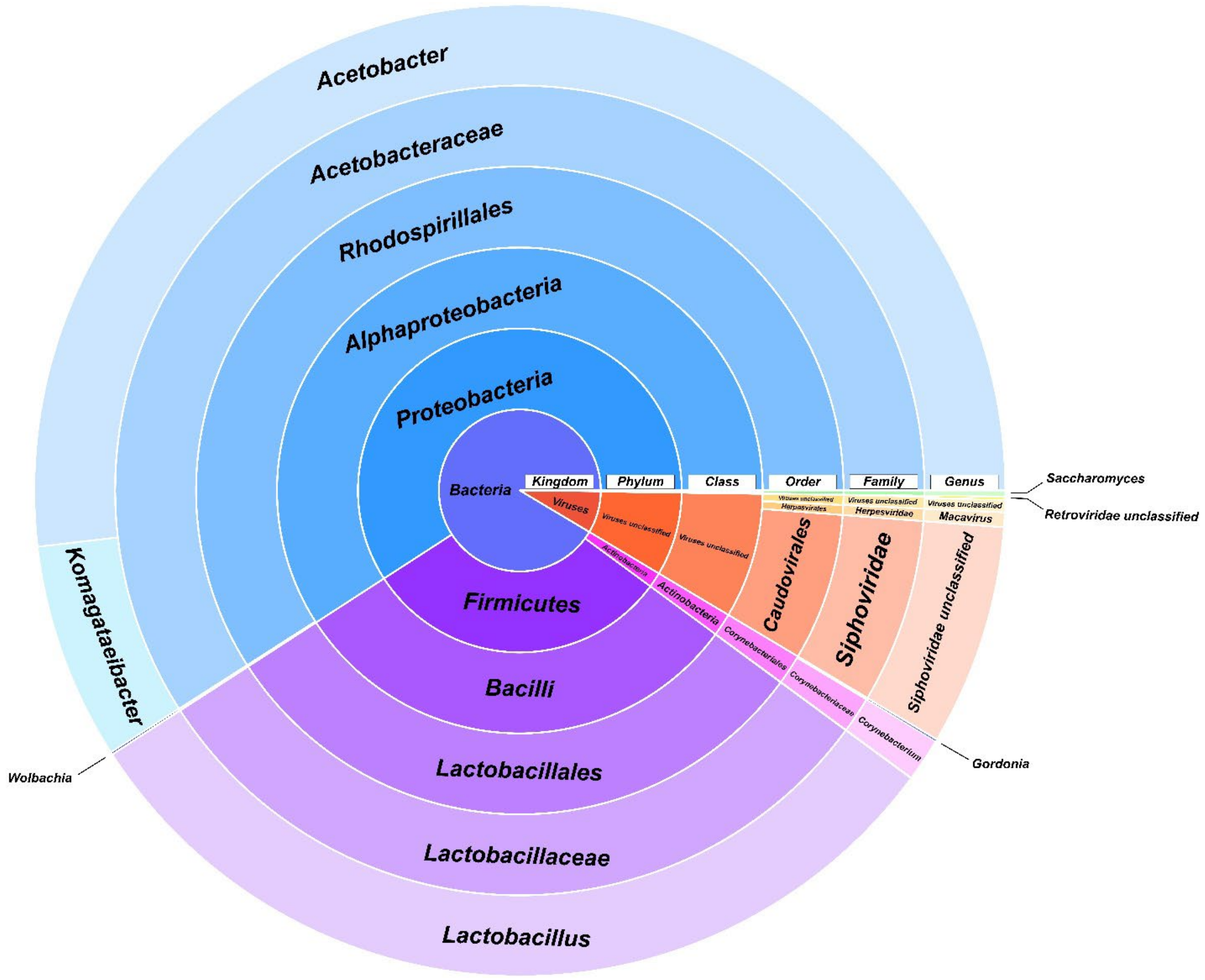

**Figure S5. Measures of microbiome diversity and richness**. (A) Shannon's $D^{85}$. (B) Simpson's $D^{86}$. (C) Simpson's $E^{87}$. Vertical lines in the violin plots represent the 25th, 50th and 75th quartiles of the distribution.

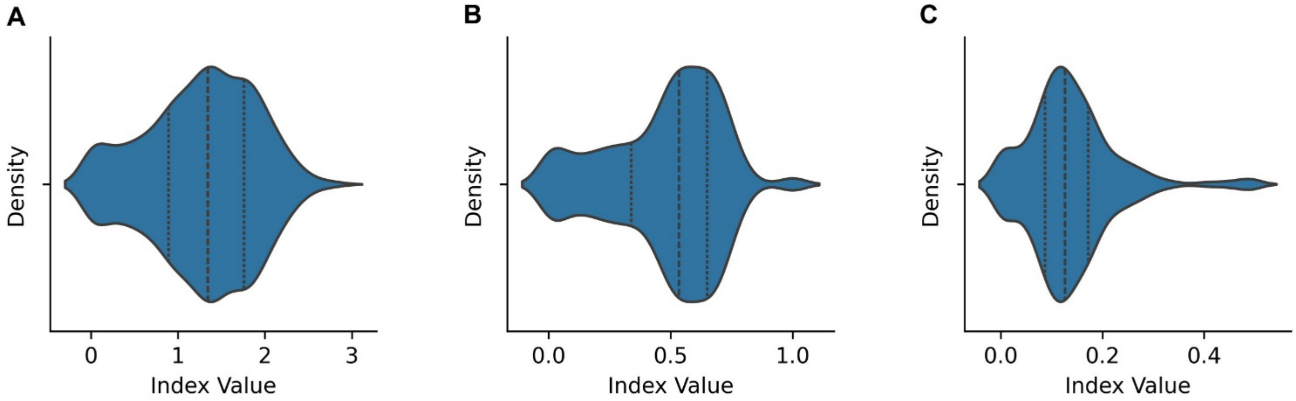

**Figure S6. Microbiome beta diversity.** (A) Bray-Curtis[88] multidimensional scaling (MDS). (B) Uniform Manifold Approximation and Projection (UMAP). Ellipses in each plot represent dense clusters of DGRP3 lines.

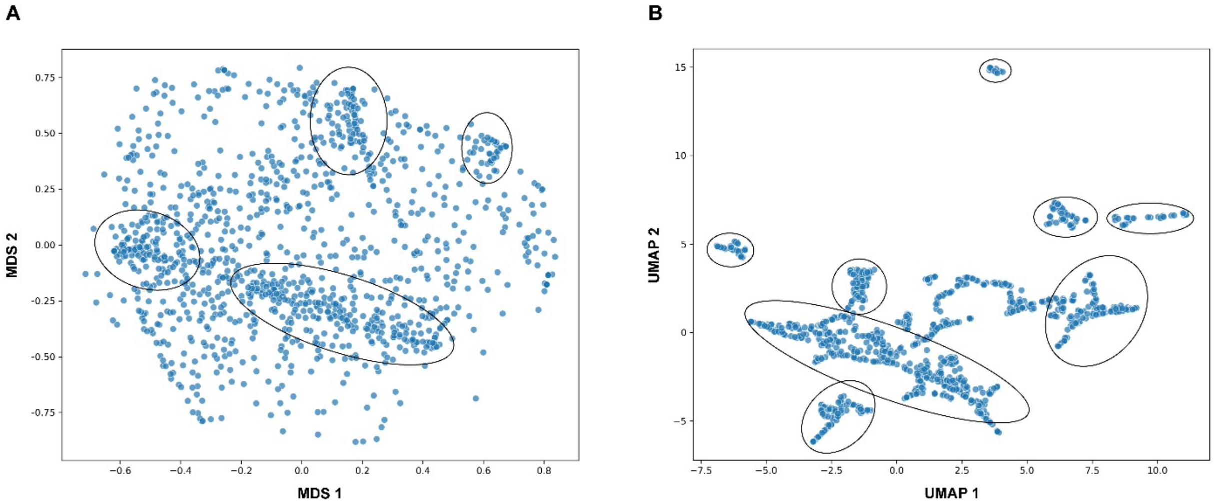

**Figure S7. Microbiome co-occurrence network.** The nodes (orange circles) denote major microbial species, and the edges represent the Spearman rank correlation coefficients between the abundances of the indicated species. Blue edges indicate negative correlations, and red edges show position correlations, with the magnitude of the correlations given by the scale bar.

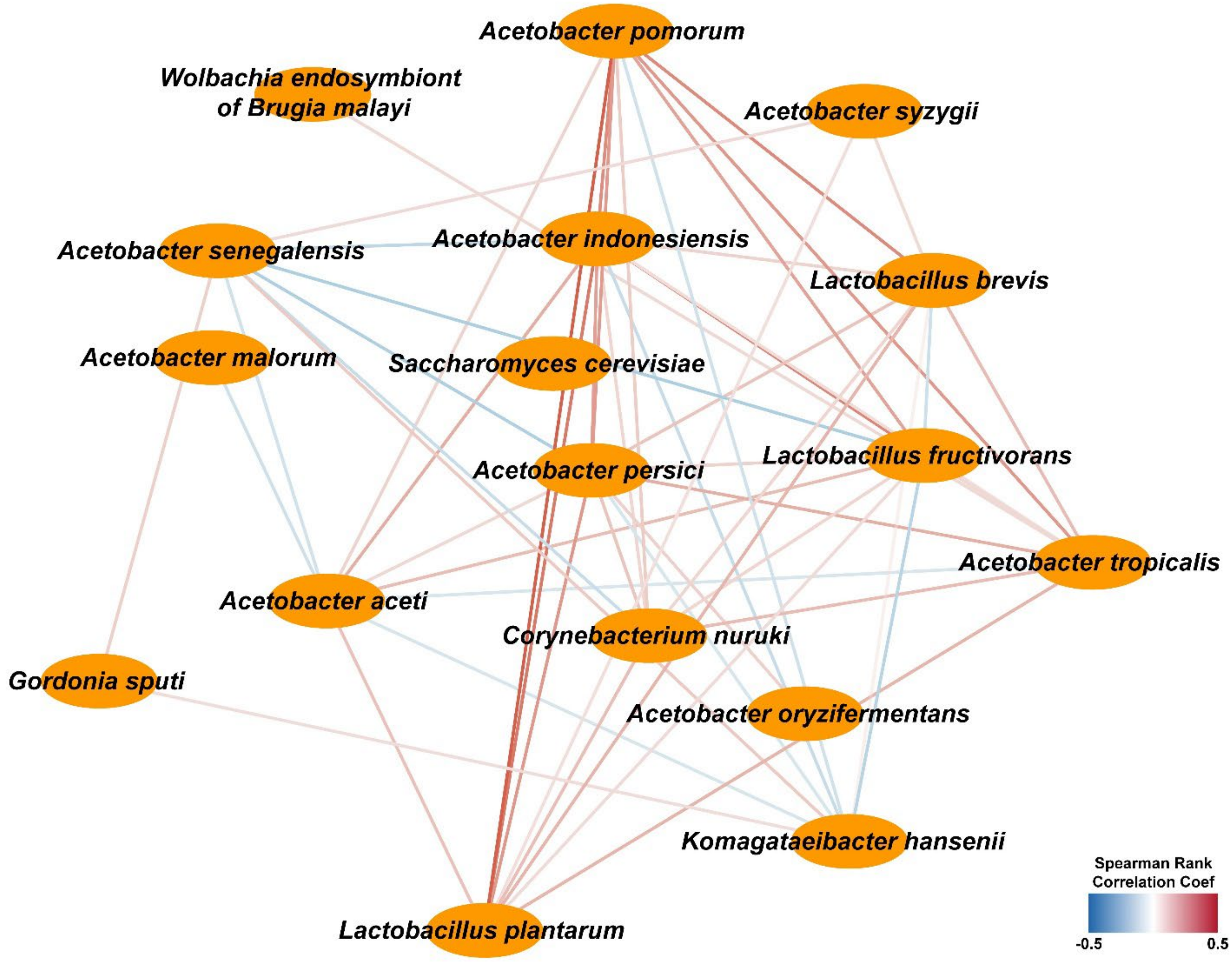

**Figure S8. Distributions of the sum of the numbers of bristles on the left and right sternopleural plates.** (A) Distribution of male line means, with lines ordered by increasing mean. (B) Frequency distribution of line means shown in (A). (C) Box plot of line means shown in (A). Distribution of female line means, with lines ordered by increasing mean. (E) Frequency distribution of line means shown in (D). (F) Box plot of line means shown in (D).

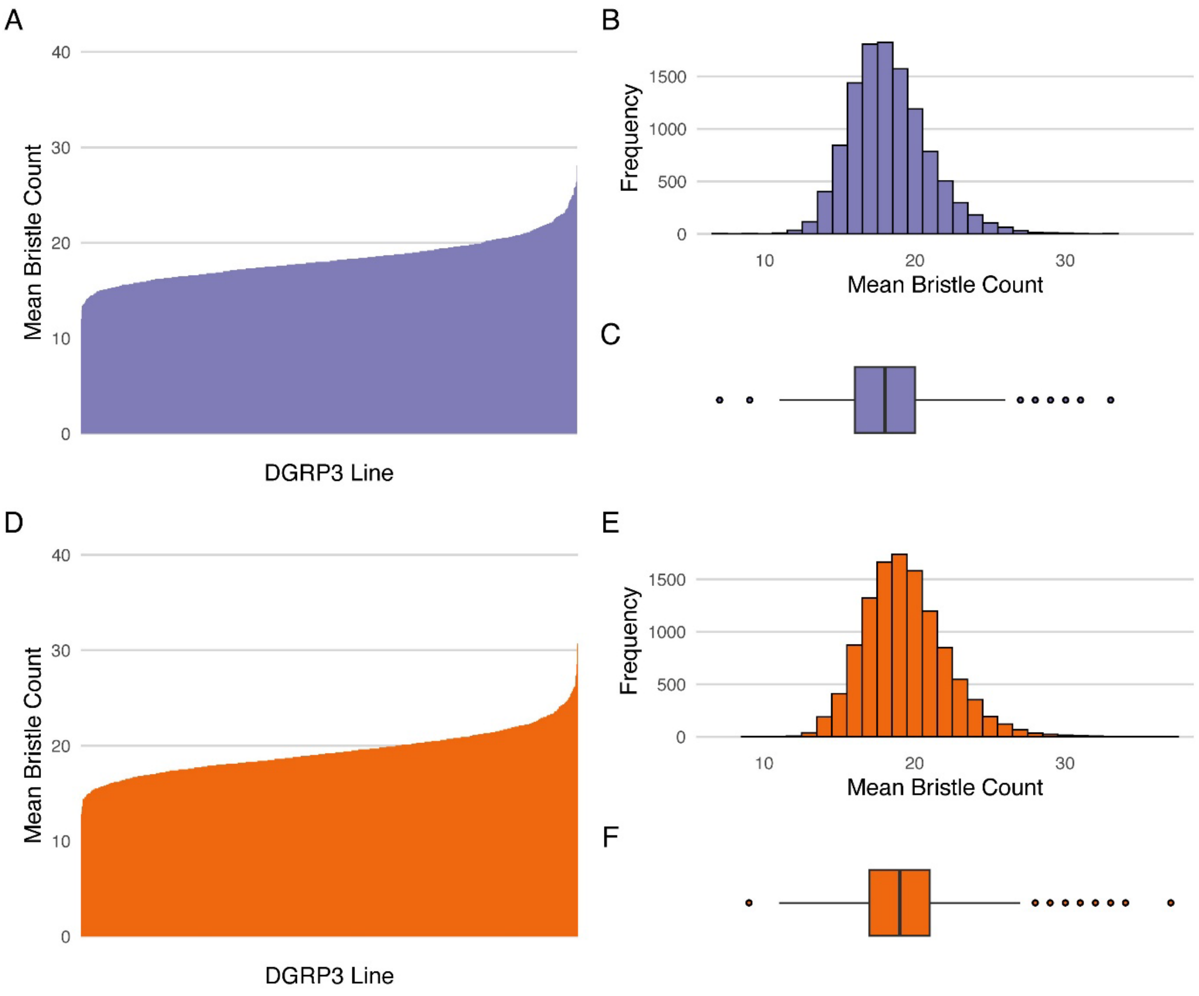

**Figure S9. Distributions of the sum of the numbers of bristles on the two most posterior abdominal sternites.** (A) Distribution of male line means, with lines ordered by increasing mean. (B) Frequency distribution of line means shown in (A). (C) Box plot of line means shown in (A). Distribution of female line means, with lines ordered by increasing mean. (E) Frequency distribution of line means shown in (D). (F) Box plot of line means shown in (D).

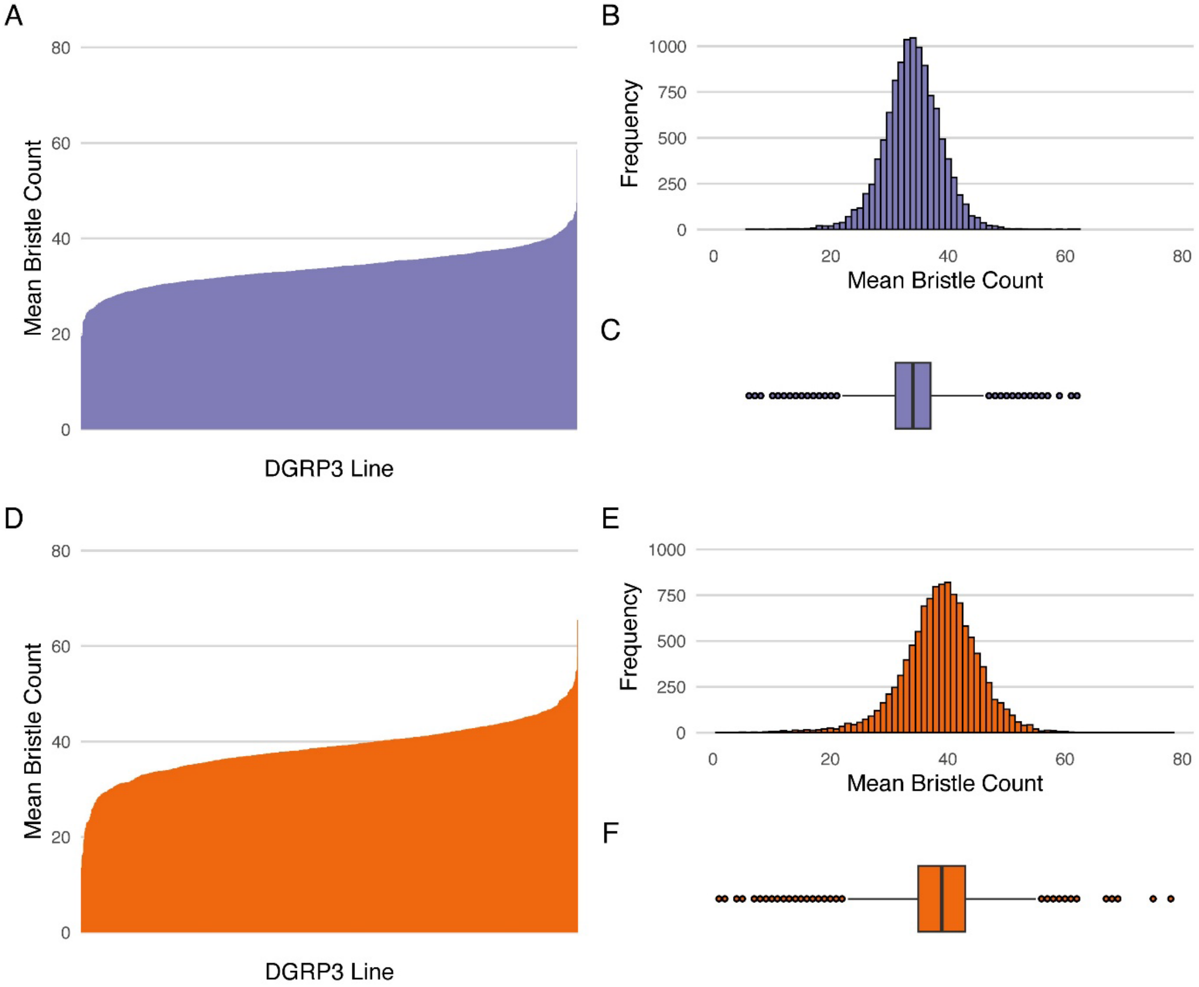

**Figure S10. Distributions of the difference of the numbers of bristles on the two most posterior abdominal sternites.** (A) Distribution of male line means, with lines ordered by increasing mean. (B) Frequency distribution of line means shown in (A). (C) Box plot of line means shown in (A). Distribution of female line means, with lines ordered by increasing mean. (E) Frequency distribution of line means shown in (D). (F) Box plot of line means shown in (D).

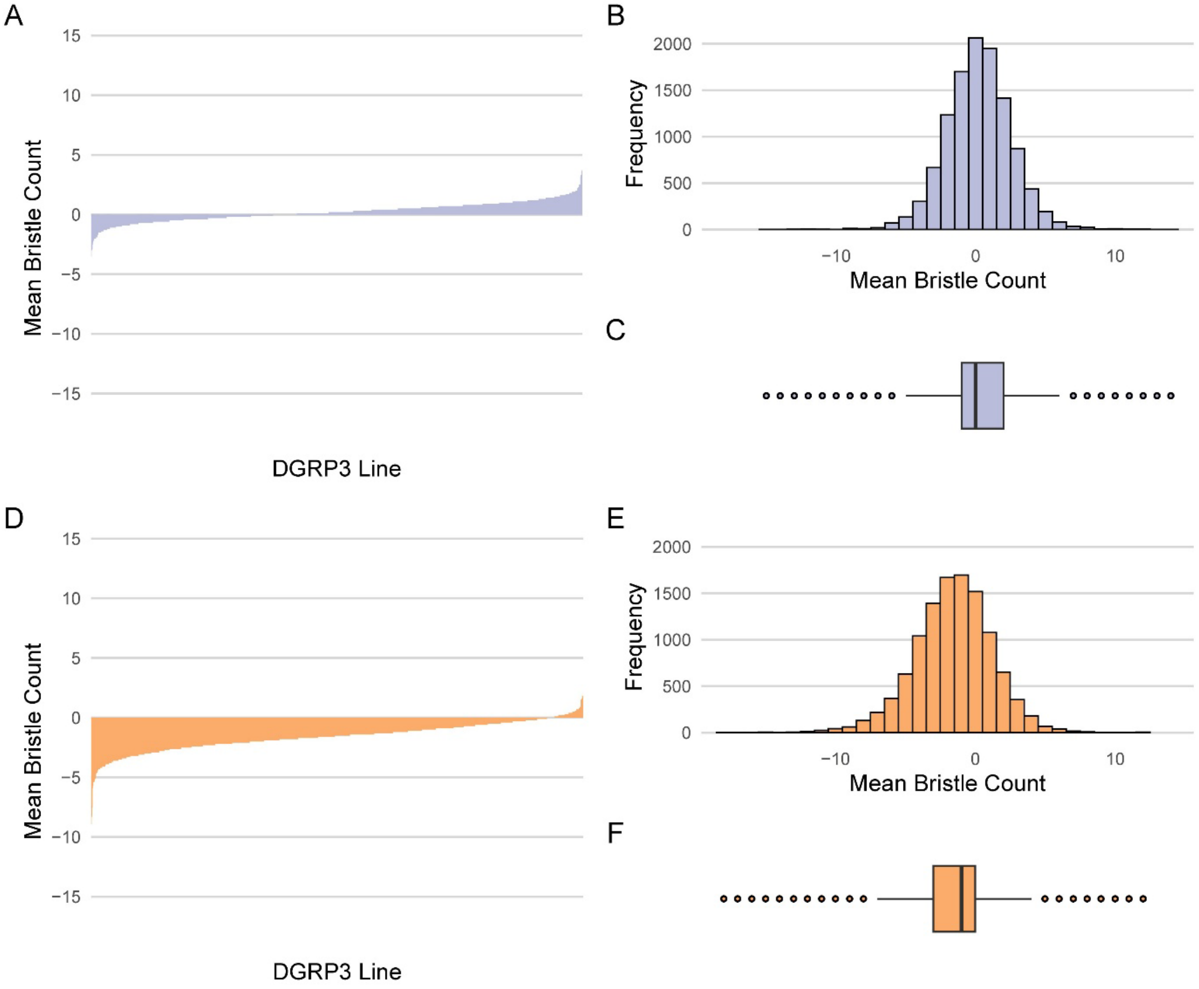

**Figure S11. Relationships between bristle trait means and micro-environmental variances ($ln(\sigma_\varepsilon)$).** (A) AB Sum: The sum of the numbers of bristles on the two most posterior abdominal sternites. (B) SB Sum: The sum of the numbers of bristles on the left and right sternopleural plates. (C) AB Diff: The difference between the numbers of bristles on the two most posterior abdominal sternites. F: females. M: Males.

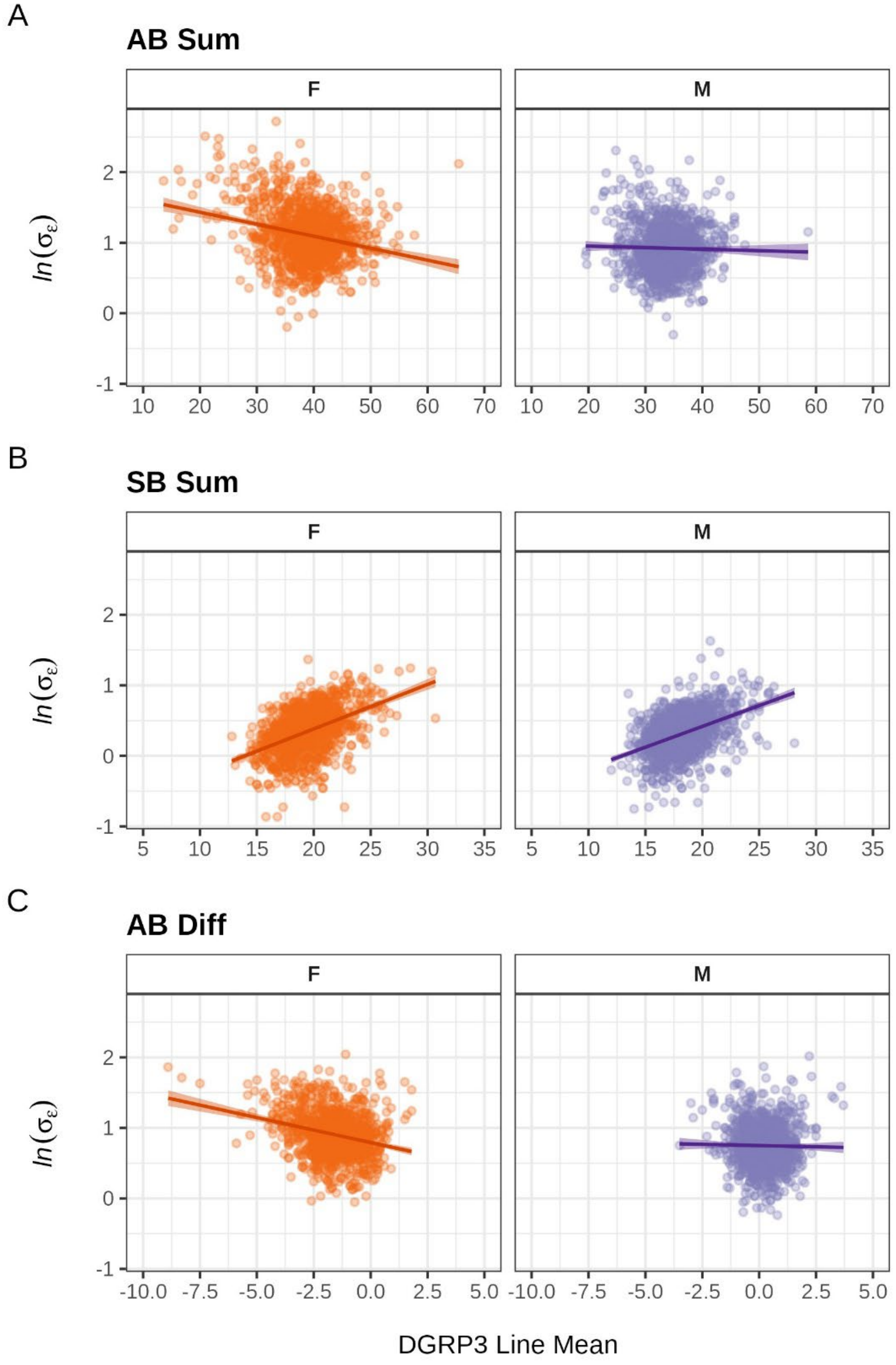

**Figure S12. Receiver Operating Characteristics (ROC) curves from Random Forest discriminant models separating DGRP3 populations.** (A-C) ROC curves and area under the curve (AUC) values for models that compare each group against the rest of the lines using all 758,001 variants. (D-E) ROC curves and AUC values for models that compare each group against the rest of the lines using only the top 216 discriminator variants.

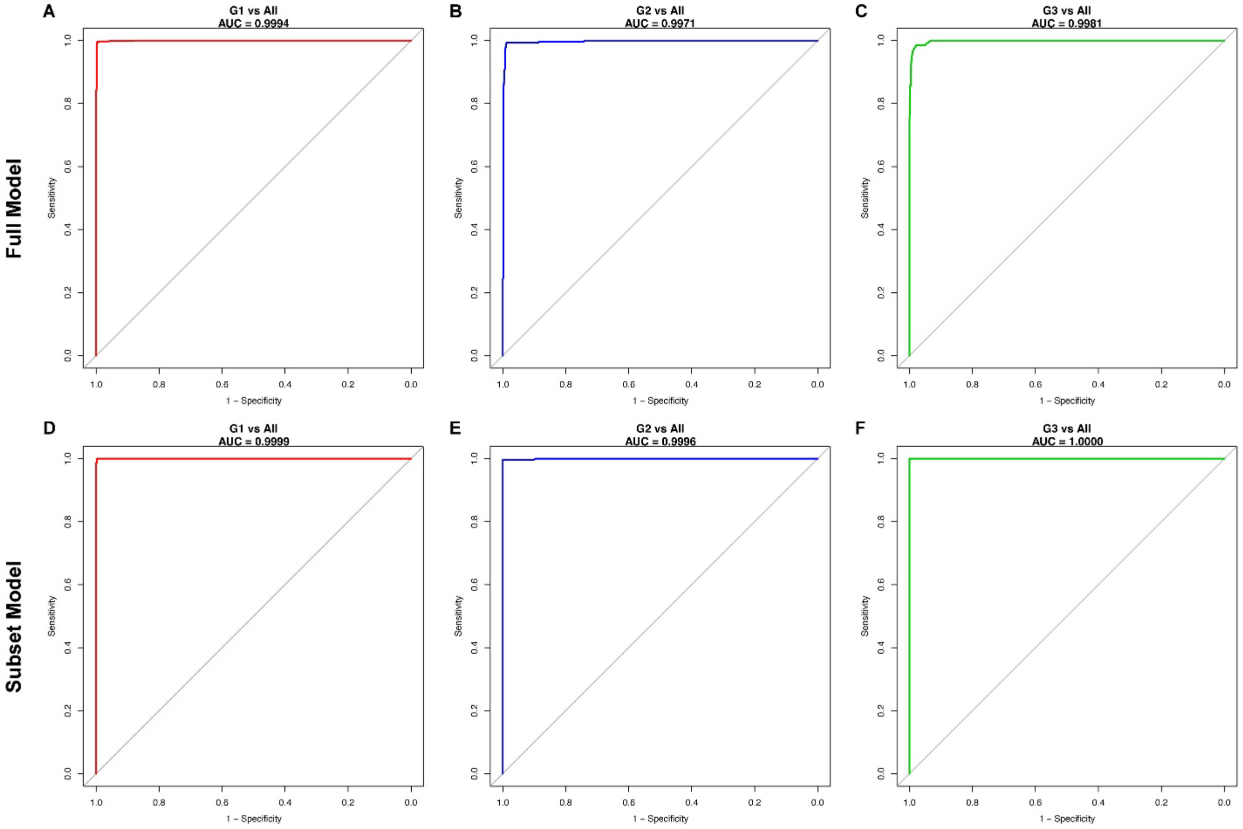

**Figure S13. Genomic distribution of polymorphic variants that discriminate among the three subpopulations in the DGRP3.** The Circos plot depicts the locations of genes, common large polymorphic inversions and the top 216 variants that discriminate among the three DGRP3 subpopulations from the random forest analysis on the *D. melanogaster* genome. The major chromosome arms and genomic positions are shown in the outer ring.

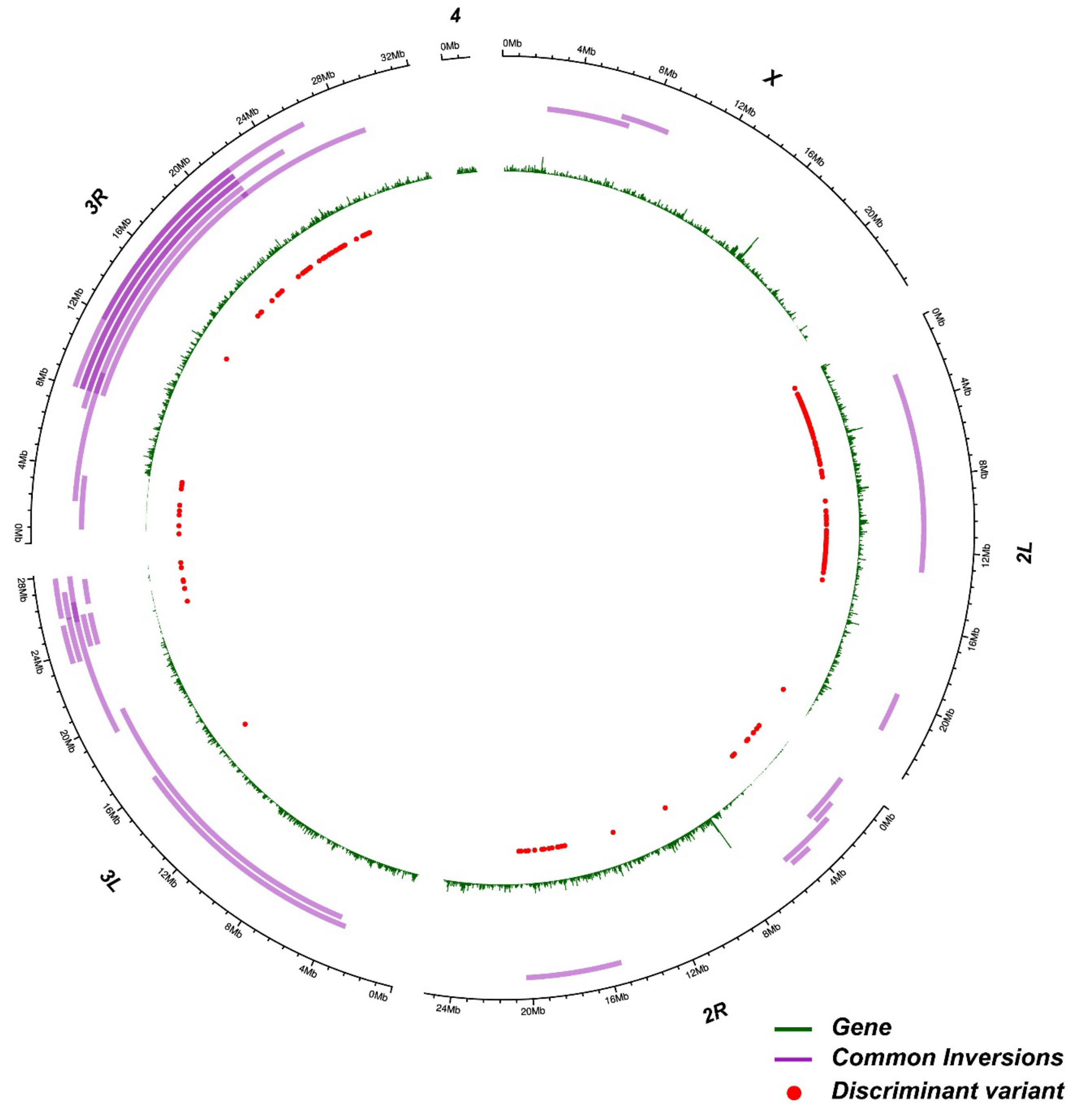

**Figure S14. Distribution of $LD$ along the major chromosome arms.** The plots depict the mean $r^2$ between variants 50-150 bp apart in 100 kb sliding windows in each 1 Mb region. (A) *X* chromosome. (B) Chromosome *2L*. (C) Chromosome *2R*. (D) Chromosome *3L*. (E) Chromosome *3R*.

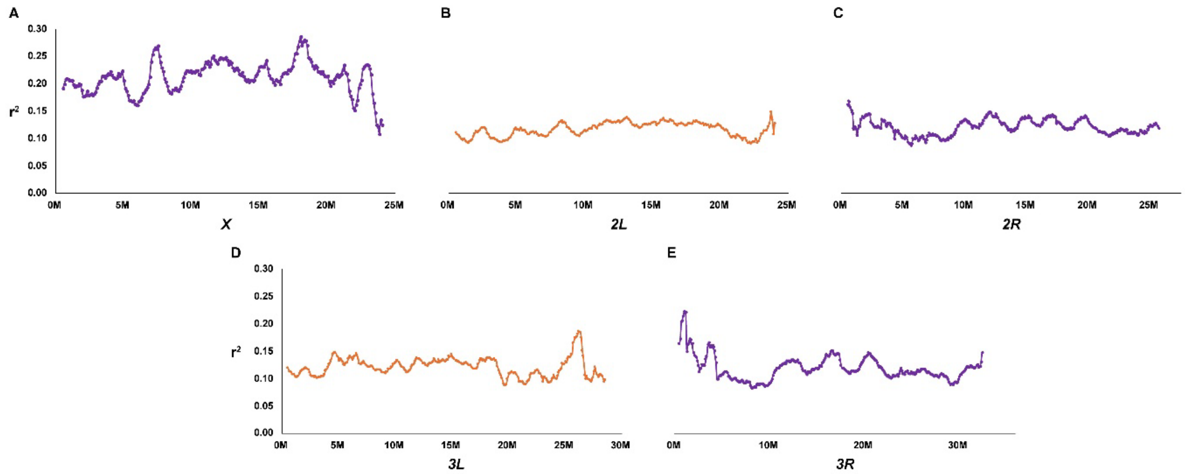

**Figure S15. Histogram of numbers of high impact homozygous variants in DGRP3 lines.**
Orange bars: extant lines. Green bars: extinct lines.

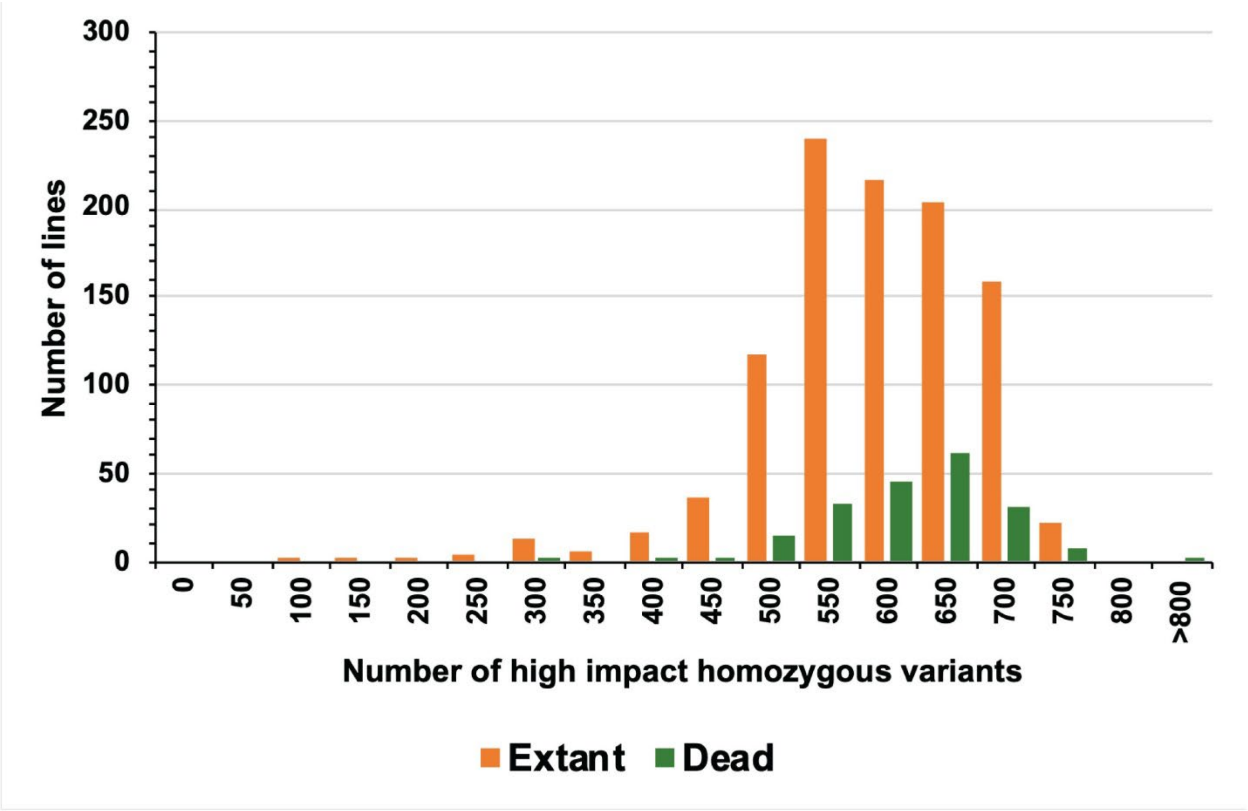

**Figure S16. GWA analysis of inbreeding depression.** (A) Manhattan plot of $-log_{10}(P)$ values. (B) Quantile-quantile plot of observed and expected $-log_{10}(P)$ values.

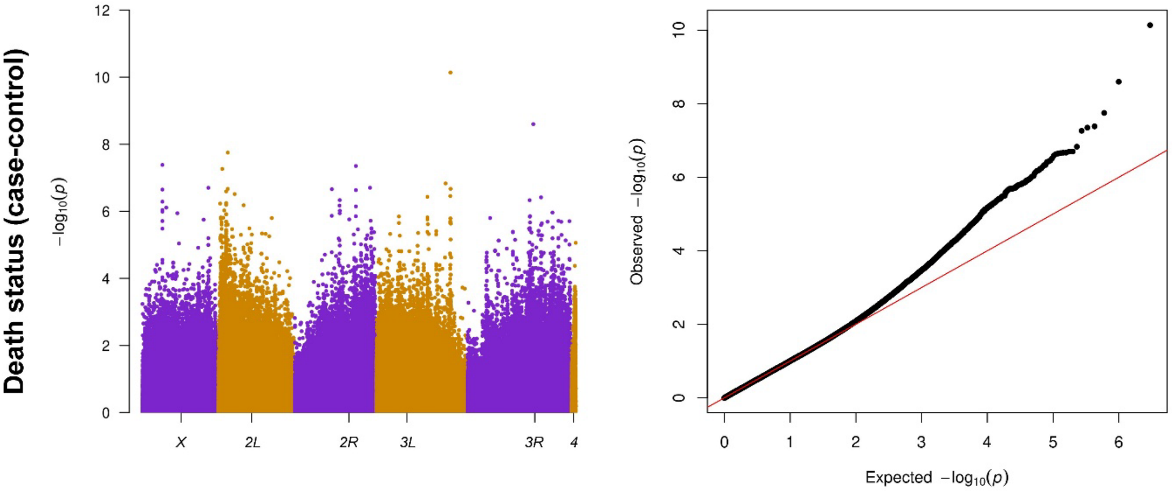

**Figure S17. GWA analyses of host control of microbiome composition**. The plots on the left are Manhattan plots of $-log_{10}(P)$ values, and on the right are quantile-quantile plots of observed and expected $-log_{10}(P)$ values. (A) Simpson's $D$[86]. (B) *Lactobacillus brevis*. (C) *Komagataeibacter hansenii*. (D) *Acetobacter pomorum*.

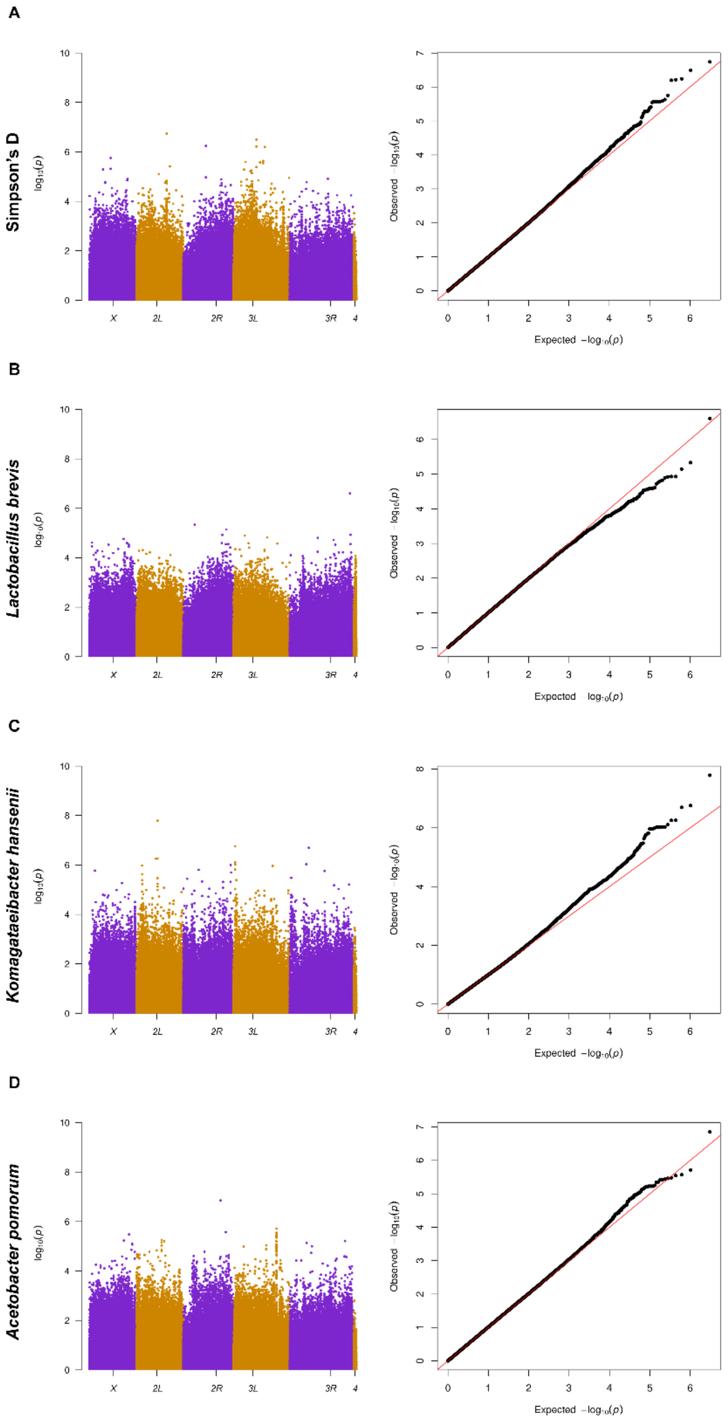

**Figure S18. GWA analyses of total sternopleural bristle (SB) numbers.** The plots on the left are Manhattan plots of $-log_{10}(P)$ values, and on the right are quantile-quantile plots of observed and expected $-log_{10}(P)$ values. (A) Males. (B) Females. (C) Average of both sexes.

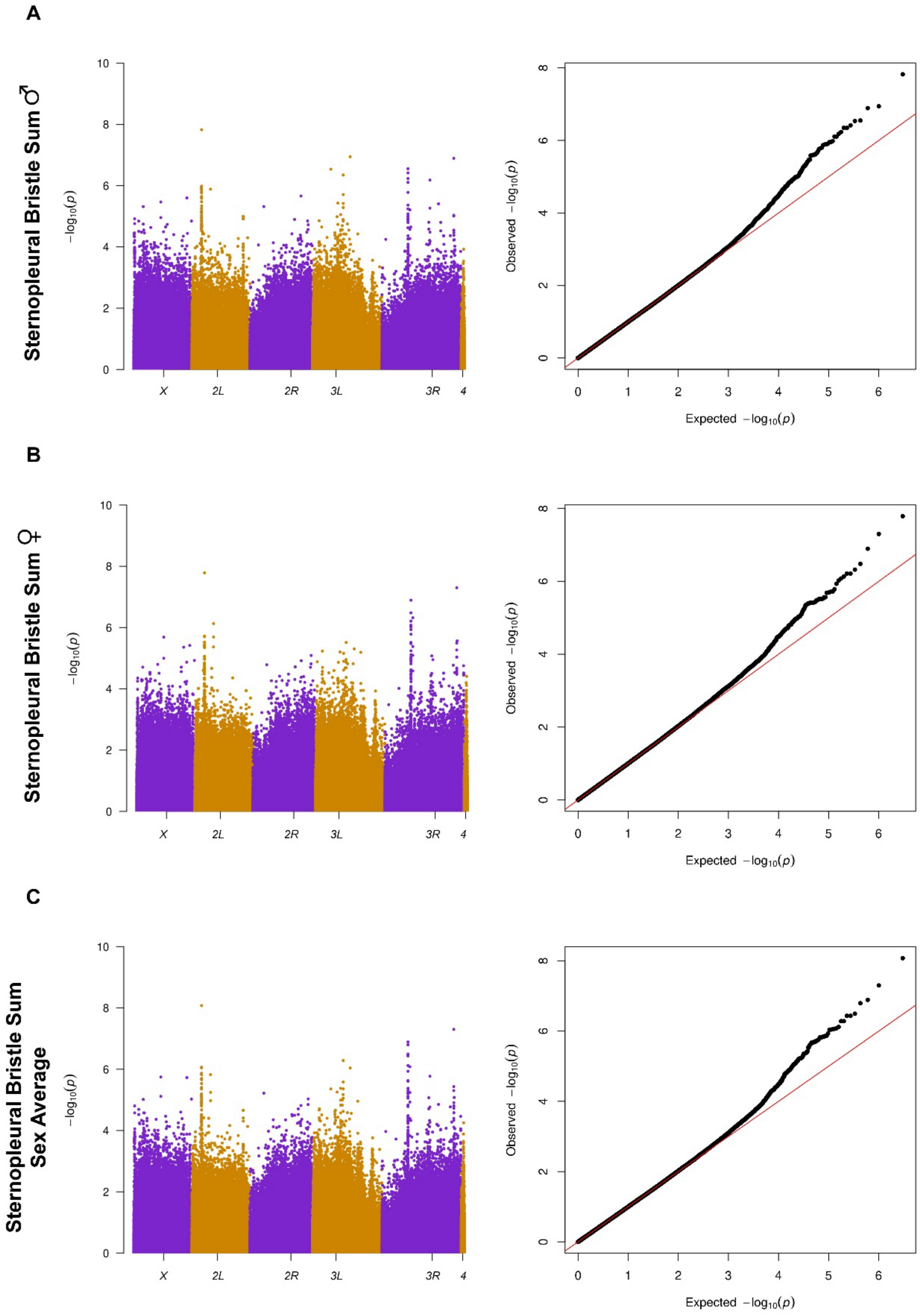

**Figure S19. GWA analyses of total abdominal bristle (AB) numbers.** The plots on the left are Manhattan plots of $-log_{10}(P)$ values, and on the right are quantile-quantile plots of observed and expected $-log_{10}(P)$ values. (A) Males. (B) Females. (C) Average of both sexes. (D) Difference between females and males.

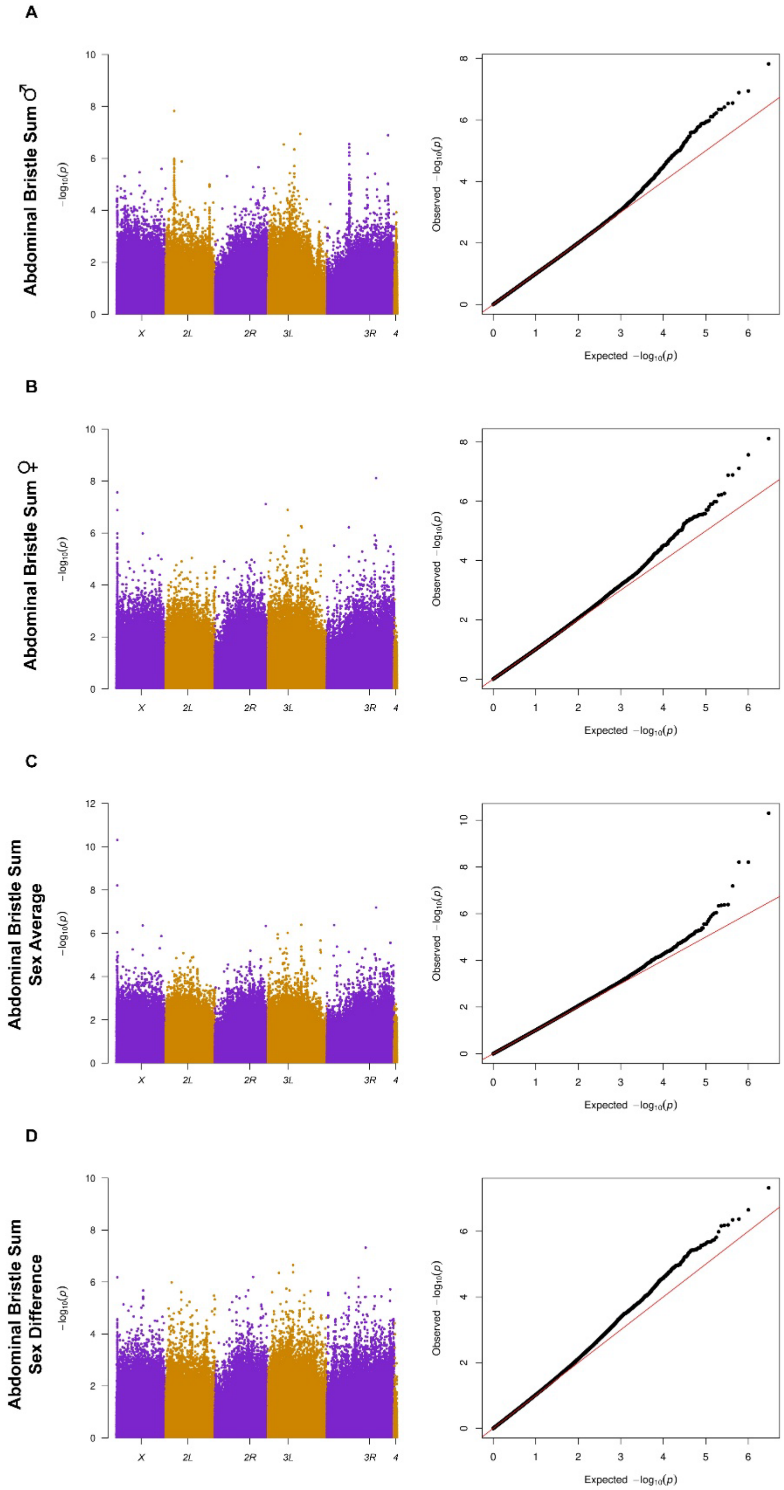

**Figure S20. GWA analyses of the difference in abdominal (AB) bristle number between the two most posterior sternites.** The plots on the left are Manhattan plots of $-log_{10}(P)$ values, and on the right are quantile-quantile plots of observed and expected $-log_{10}(P)$ values. (A) Males. (B) Females. (C) Average of both sexes. (D) Difference between females and males.

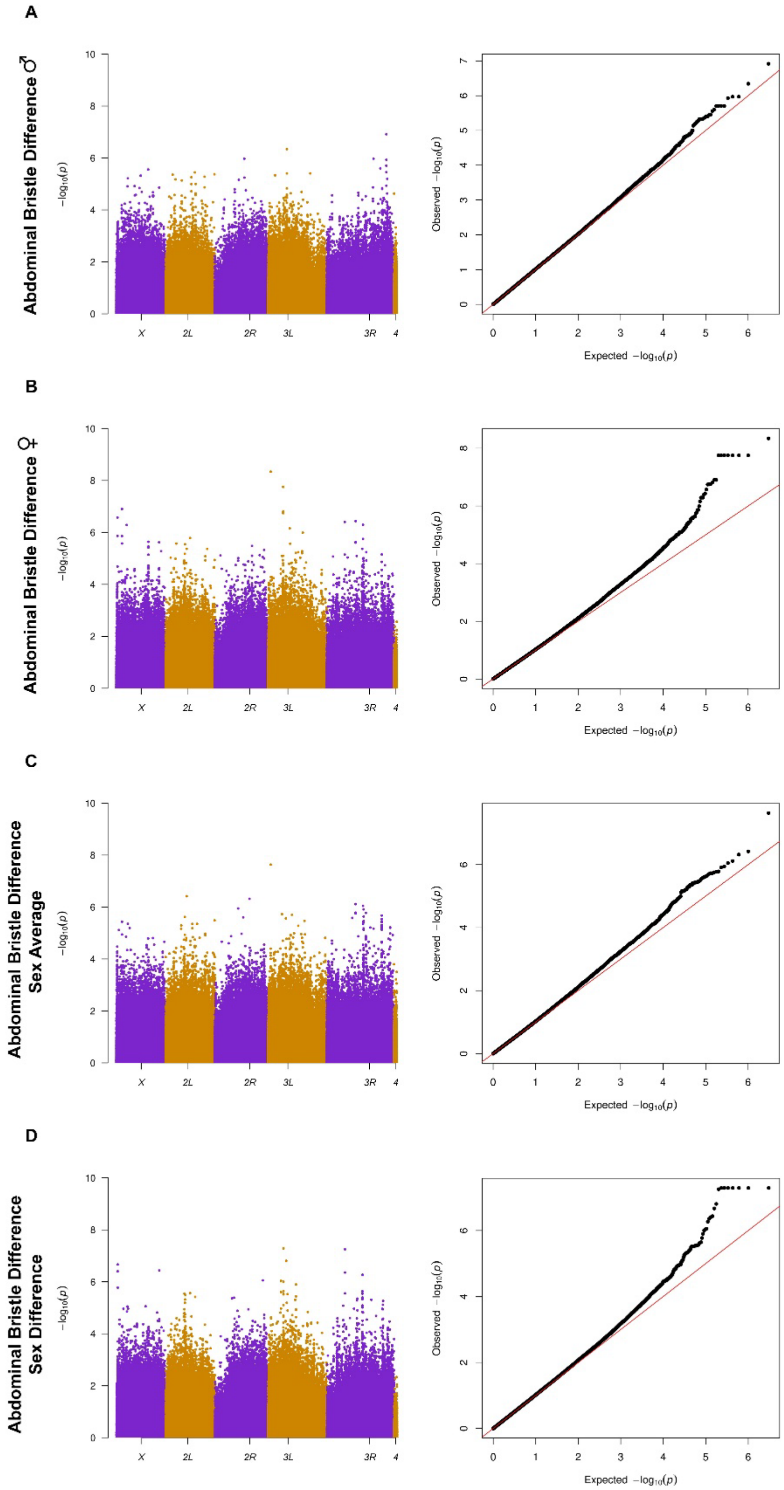

**Figure S21. GWA analyses of $ln\sigma_{\varepsilon}$ total abdominal bristle (AB) number.** The plots on the left are Manhattan plots of $-log_{10}(P)$ values, and on the right are quantile-quantile plots of observed and expected $-log_{10}(P)$ values. (A) Males. (B) Females.

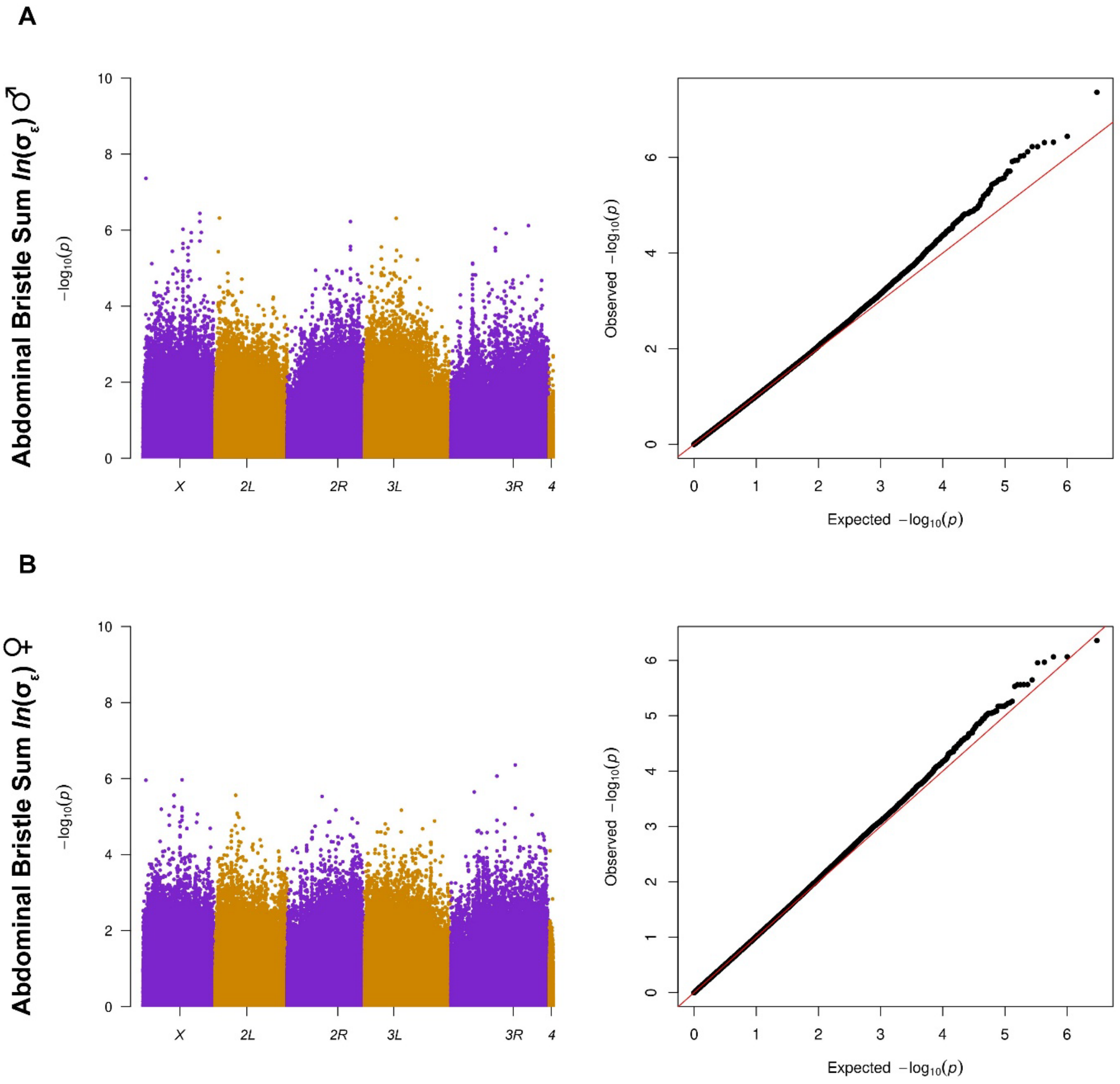

**Figure S22. Statistical power to detect associations.** The power is calculated based on the 1-df $\chi^2$ test. For a standardized effect size of $\beta$, the explained heritability is $2p(1-p)\beta^2$. The heritability of line means is assumed to be 1. Under these conditions, the test statistic under the alternative hypothesis is distributed as a non-central $\chi^2(\lambda)$ distribution with non-centrality parameter $\lambda = n \times 2p(1-p)\beta^2$, where *n* is the number of lines. With a *P*-value threshold of $\alpha = 10^{-5}$, the power to detect the association is $P(\chi^2(\lambda) > \chi^2)$. The power is plotted against effect size for different MAF when the number of lines is (A) 200 or (B) 1,200.

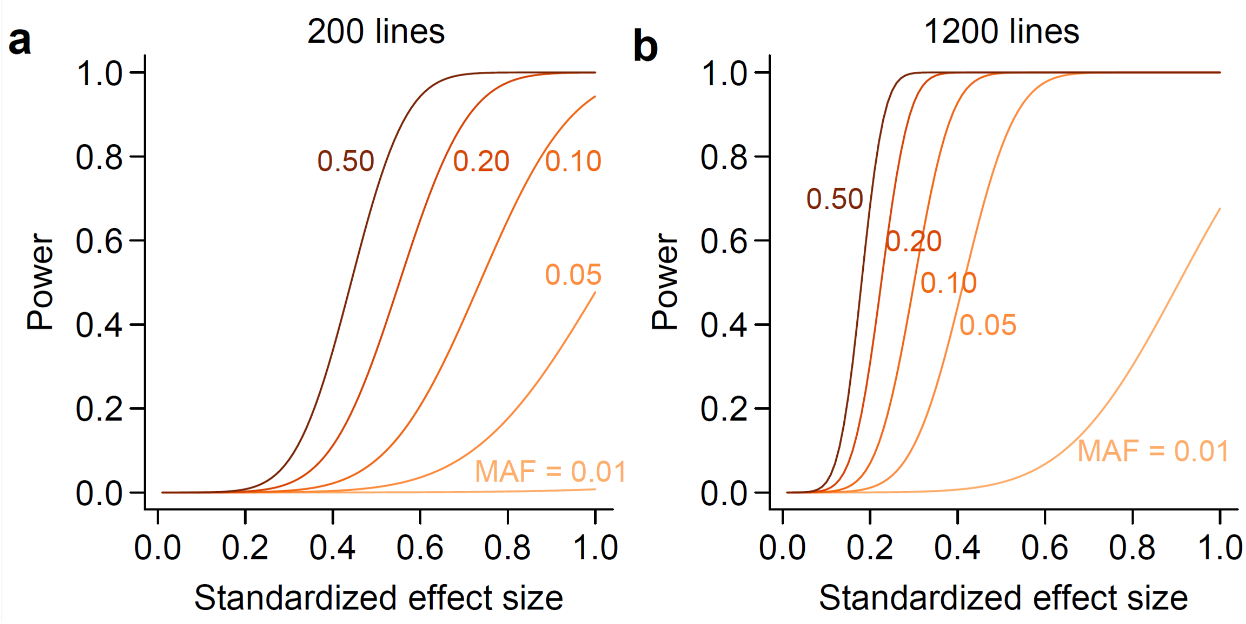

**Figure S23. Discovery of genotype-phenotype associations at increasing sample size.** The subsampling was performed by first randomly sampling 200 lines from the full dataset ($n$ = 1,124) and incrementally adding more lines. GWA was performed on quantile-normalized female abdominal bristle number to mitigate effects of outliers. SNPs and genes were discovered at a nominal $P$-value threshold of $10^{-5}$. The MAF cutoff was 0.05 for 200 lines, 0.02 for 500 lines, and 0.01 for 1,000 and 1,124 lines.

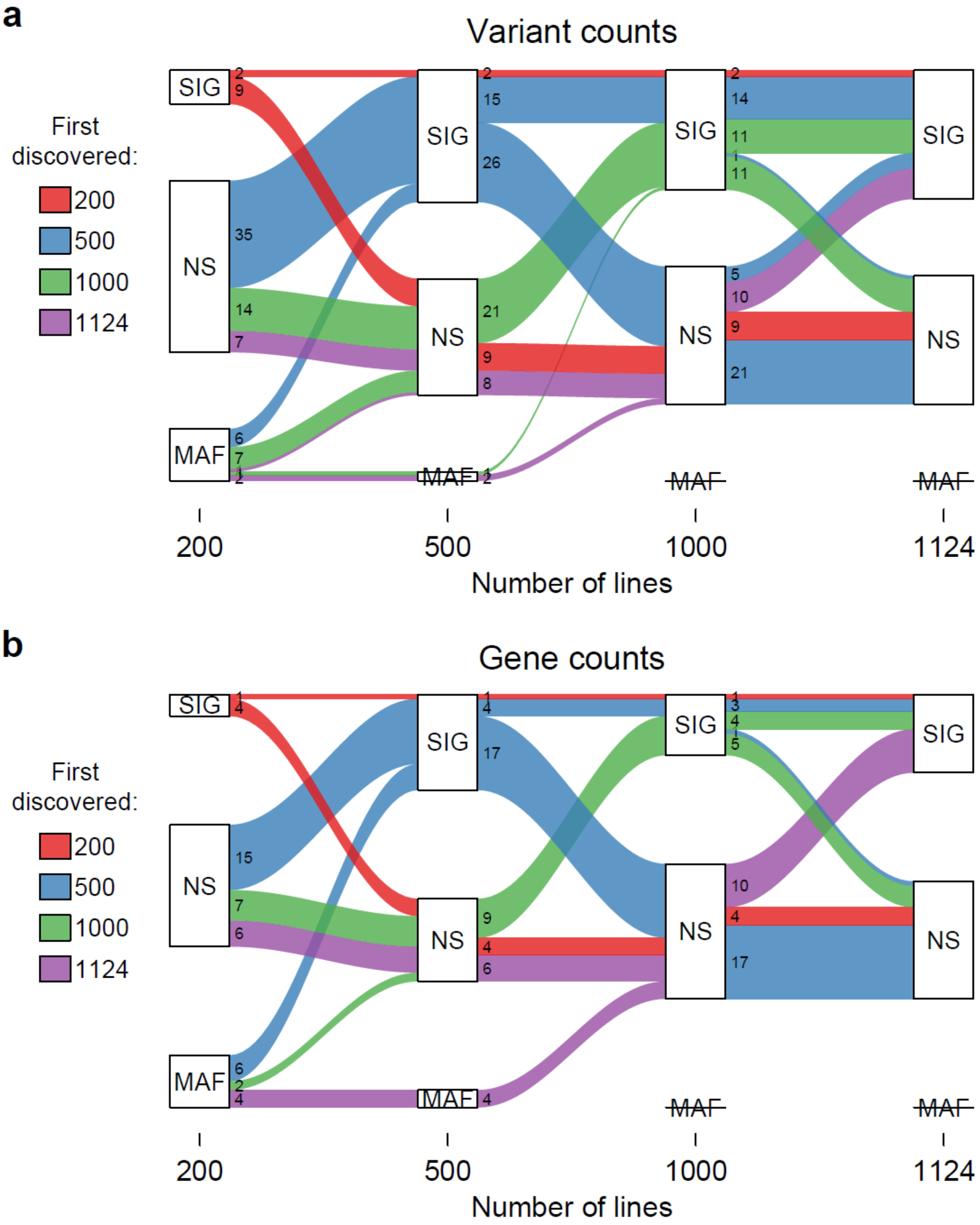
